# Mutualism-enhancing mutations dominate early adaptation in a microbial community

**DOI:** 10.1101/2021.07.07.451547

**Authors:** Sandeep Venkataram, Huan-Yu Kuo, Erik F. Y. Hom, Sergey Kryazhimskiy

## Abstract

Species interactions drive evolution while evolution shapes these interactions. The resulting eco-evolutionary dynamics, their outcomes and their repeatability depend on how adaptive mutations available to community members affect fitness and ecologically relevant traits. However, the diversity of adaptive mutations is not well characterized, and we do not know how this diversity is affected by the ecological milieu. Here we use barcode lineage tracking to address this gap in a competitive mutualism between the yeast *Saccharomyces cerevisiae* and the alga *Chlamydomonas reinhardtii*. We find that yeast has access to many adaptive mutations with diverse ecological consequences, in particular, those that increase and reduce the yields of both species. The presence of the alga does not change which mutations are adaptive in yeast (i.e., there is no fitness trade-off for yeast between growing alone or with alga), but rather shifts selection to favor yeast mutants that increase the yields of both species and make the mutualism stronger. Thus, in the presence of the alga, adaptations contending for fixation in yeast are more likely to enhance the mutualism, even though cooperativity is not directly favored by natural selection in our system. Our results demonstrate that ecological interactions not only alter the trajectory of evolution but also dictate its repeatability; in particular, weak mutualisms can repeatably evolve to become stronger.

## Introduction

Ecological communities are often perturbed by environmental shifts^1,2^, demographic noise^3^ and species turnover^4,5^. Such perturbations can not only displace communities from their ecological equilibria but also precipitate adaptive evolution^6–9^. Evolutionary changes within one species can be rapid and can alter its ecological interactions with other community members, which can cause further evolution^8–10^. Although such eco-evolutionary feedbacks appear to be widespread^7,9–25^, the population genetic mechanisms that underlie them are not well understood^26^. For example, how does the spectrum of adaptive mutations available to a species (i.e. the genomic locations and fitness effects of mutations that provide fitness benefits) depend on the composition of the surrounding community? In particular, how does it change when a species is lost from the community or a new species invades it? How many and which of the adaptive mutations available to a species affect its interactions with the rest of the community? Which of these mutations are likely to spread and fix? And thus, how diverse and repeatable are the ecological outcomes of evolution?

Empirical data supporting answers to these questions would help us develop a better theoretical understanding of eco-evolutionary dynamics. For example, many existing models assume that any combination of traits can be produced by mutations so that the eco-evolutionary trajectories and outcome are determined exclusively by natural selection^27^. However, recent evidence suggests that the availability of mutations can significantly impact evolution^28–32^. Yet, we know very little about the distributions of ecological and fitness effects of new mutations in multi-species communities and how these distributions shift when the ecological milieu changes—for example, due to the addition or extinction of community members.

Here, we address this gap in one of the simplest experimentally tractable microbial communities. Our community consists of two species, the alga *Chlamydomonas reinhardtii* and the yeast *Saccharomyces cerevisiae*, that interact in our environment via competition and mutualism^33^. Although communities in nature often contain more members, understanding eco-evolutionary dynamics in simple model communities is helpful for developing an intuition and expectations for the behaviors of more complex ecosystems^34–36^. We measure how adaptive mutations arising in one member of our community, the yeast, affect its competitive fitness (a metric that determines the evolutionary success of a mutant lineage), the absolute abundances of both species in the community (a metric that informs us about the type of interactions between species and the stability of the community) as well as basic life-history traits of yeast (growth rates and carrying capacities) that contribute to both fitness and abundances. We specifically ask whether and how the statistical distribution of effects of adaptive mutations in yeast are altered by the presence/absence of the alga. To this end, we use the barcode lineage tracking (BLT) technology^37,38^ to isolate hundreds of adaptive mutations arising in yeast when it evolves alone or in community with the alga. Our data offer us a detailed view on how inter-species interactions affect the evolutionary dynamics of new mutations, and how these mutations in turn alter the ecology of our community.

## Results

### Yeast and alga form a facultative competitive mutualism

In a previous study, Hom and Murray showed that in a sealed environment in which nitrite is provided as a sole source of nitrogen and glucose is provided as a sole source of organic carbon, the yeast *Saccharomyces cerevisiae* and the alga *Chlamydomonas reinhardtii* spontaneously form an obligate mutualism^33^. Under such conditions, *C. reinhardtii* consumes nitrite and produces ammonium that is secreted and utilized by *S. cerevisiae*, which consumes glucose and produces CO_2_ that is in turn utilized by the alga. When the environment is opened to ambient gas exchange and ammonium is supplied in the medium, both the yeast and the alga can survive without each other. In this study, we grow the yeast and the alga alone and together over multiple 5-day growth and dilution cycles in well-mixed and well-lit conditions, open to gas exchange, in a medium supplemented with 0.5 mM of ammonium (see Methods for details).

Time-course cell-density measurements of the wild-type yeast and alga over a single cycle confirm that both species grow significantly differently in each other’s presence than alone (Figure 1, Extended Data Figure 1, Data S1, repeated-measures ANOVA *P* = 10^−4^ for the yeast and *P* = 5×10^−8^ for the alga), indicating that they ecologically interact in our experimental environment. Specifically, the alga achieves higher densities over the entire growth cycle in the community compared to growth alone. The presence of the alga alters yeast growth dynamics in a more complex way. When yeast grows alone, it reaches peak cell density at 48 hours, after which point its density gradually declines (Figure 1A), suggesting that it exhausts the initial supply of ammonium after about 48 hours. When yeast grows in community with the alga, the two species initially compete, likely for ammonium. This can be seen by a reduction of the peak yeast cell density at 48 hours (Figure 1A). Upon the depletion of supplemented ammonium, the alga subsequently reduces nitrite to ammonium that it then secretes^33^. This nitrogen provisioning by the alga reduces the yeast’s rate of population decline between days 3 and 4 (t-test *P* = 5×10^−4^, Extended Data Figure 1, Table S1). As a result, yeast reaches approximately the same density by the end of the cycle in the community as it does alone, despite having a lower peak density on day 2. Thus, in the latter portion of the growth cycle, the yeast experiences a benefit from its interaction with the alga. Since yeast and alga initially compete and later cooperate in our conditions, we refer to our system as a competitive mutualism^39^.

**Figure 1.**
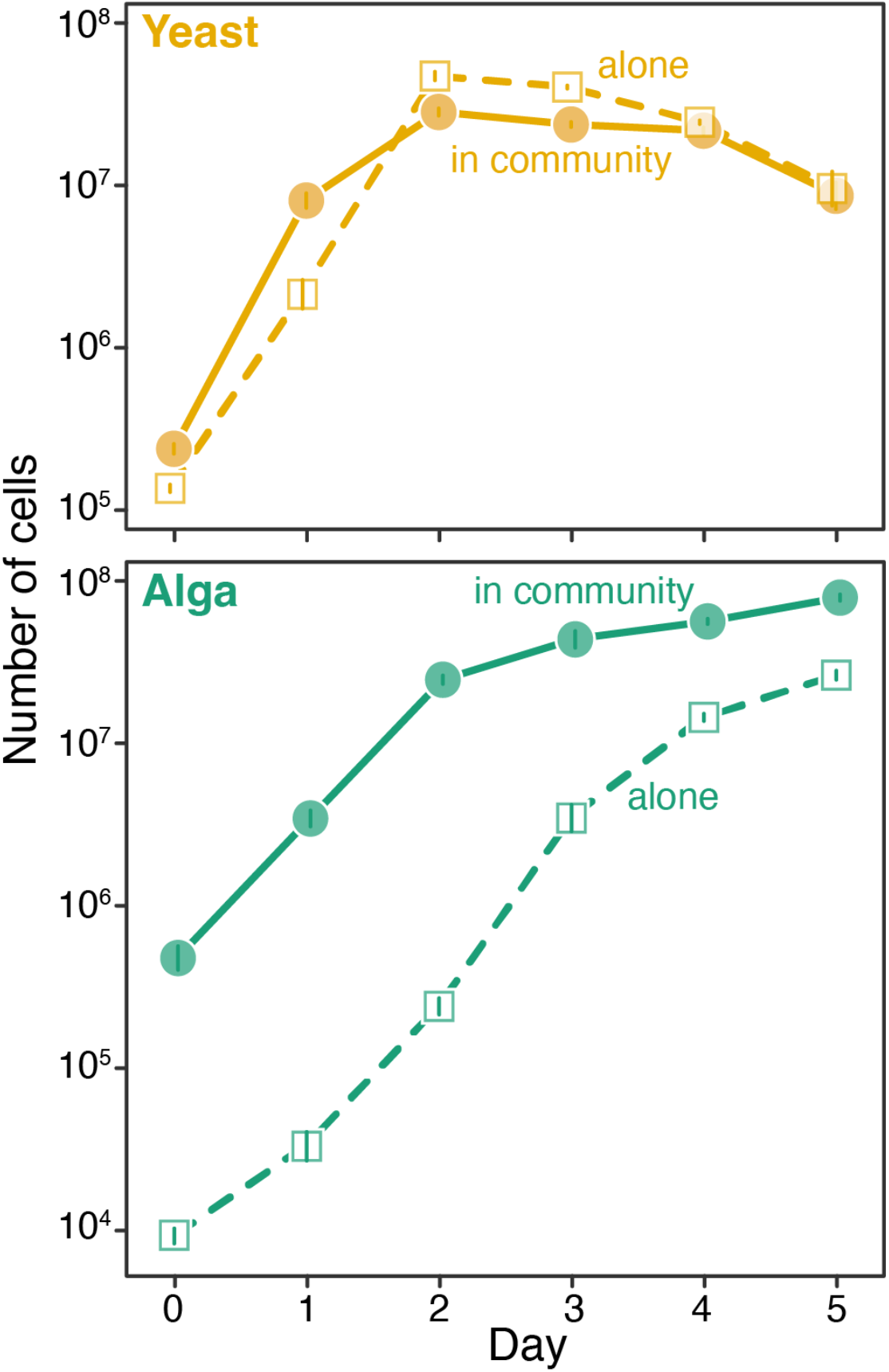
Growth of ancestral yeast and alga. Growth of the ancestral yeast and alga alone (solid lines) and in a community (dashed lines) over the 5-day cycle. Error bars show ±1 standard error of the mean. Growth curve data can be found in Data S1.

Although the ecological interactions in our system are quite complex, it would be convenient to quantify them with some simple summary statistics. We can do so, for example, by comparing the final densities that each species achieves at the end of the 5-day cycle when growing in community versus alone (Figure 1). We refer to these final densities as “yields”. Specifically, we compute the ratio of yeast yield in community (YYC) to its yield alone (YYA) and the ratio of alga yield in community (AYC) to its yield alone (AYA). Both ratios exceeding unity indicate that cooperation is on the whole more important than competition. Conversely, when both ratios are less than one, competition is more important on the whole than cooperation. For our wildtype community, we find that YYC to YYA ratio is not significantly different from one, while the AYC to AYA ratio equals 3.00 (95% Confidence Interval (CI) [2.64, 3.36], Student’s t-test *t* = 13.24, df=7.12, *P* = 3×10^−6^; Figure 1). This indicates that, on the whole, the alga benefits from its interactions with the yeast, while the yeast, on the whole, neither benefits nor suffers from its interactions with the alga. Thus, according to this metric, yeast and alga form a net commensal relationship. However, we emphasize that this net commensalism is a result of a balance between the underlying competitive and cooperative interactions.

The fact that the yeast and the alga can grow in our conditions either alone or together as a community allows us to inquire how evolution of one species is affected by its ecological interactions with the other, under otherwise identical environmental conditions. We focus on the initial phase of adaptive evolution. Since the yeast is likely to adapt faster than the alga (see Supplementary Information), we characterize the distribution of ecological and fitness effects of adaptive mutations arising in yeast and examine how these distributions depend on the presence/absence of the alga.

### The presence of the alga alters the fitness effects of beneficial mutations in yeast without apparent trade-offs

We first asked whether and how ecological interactions with the alga change the distribution of fitness effects of adaptive mutations arising in yeast. To this end, we carried out five replicate BLT experiments in yeast evolving <u>a</u>lone (the “A-condition”) and in a <u>c</u>ommunity with the alga (the “C-condition”; Methods). In each population, we tracked the frequencies of ~5×10^5^ neutral DNA barcodes integrated into the yeast genome^40^ for 17 growth and dilution cycles. We identified on average 2,820 and 2,905 adapted barcode lineages per culture in A- and C-conditions, respectively (Methods; Extended Data Figures 2, 3, Figures S1–S8). The similarity of these numbers suggests that the presence of the alga does not dramatically change the rate at which beneficial mutations arise in yeast.

Each adapted lineage is expected to initially carry a single beneficial driver mutation^37,38^. Thus, by tracking barcode frequencies, we can estimate the competitive fitness benefits of many simultaneously segregating driver mutations relative to the ancestral yeast strain (Methods). The estimated distributions of fitness effects of beneficial mutations (bDFEs) are much broader than expected from measurement noise alone in both A- and C-conditions (Supplementary Information), indicating that yeast has access to multiple mutations with different fitness benefits, consistent with previous work^37,41^. The bDFEs in A- and C-conditions are different in both median effect and breadth (Figure 2A, S6 and S7). Specifically, the presence of the alga reduced the bDFE median (1.60 in A vs. 1.50 in C; *P* < 10^−4^, two-sided permutation test, see Methods) and increased its width (interquartile range (IQR) = 0.31 in A vs. IQR = 0.37 in C; *P* < 10^−4^, two-sided permutation test). This increase in width is associated with the appearance of two peaks with higher relative fitness values around 2.0 and 2.5 (Figures 2A and S6). Since the dynamics of adaptation depend on the shape of the bDFE^42,43^, these results indicate that the presence of the alga alters evolutionary dynamics in the yeast population.

**Figure 2.**
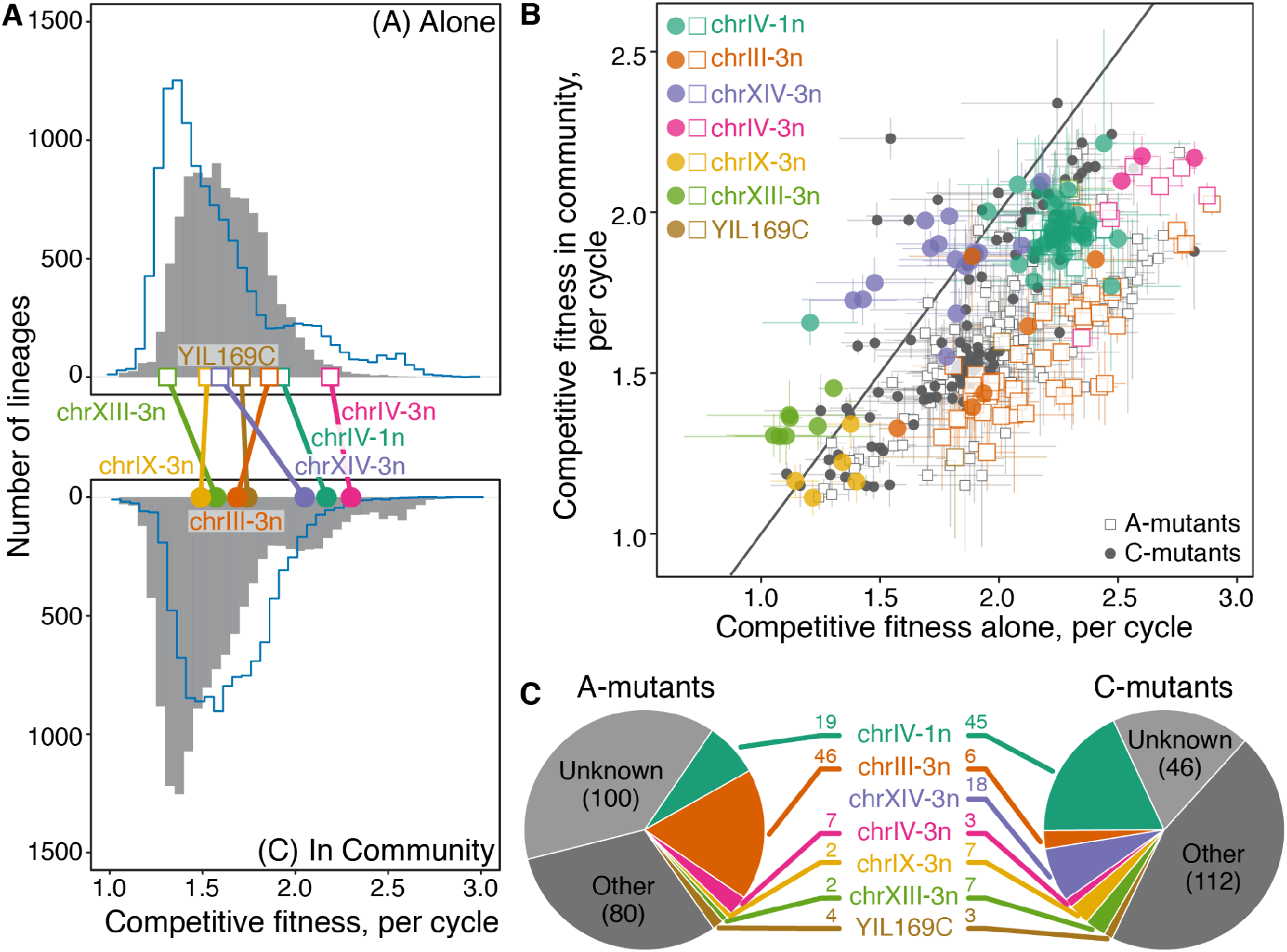
The presence of the alga affects adaptation in yeast at fitness and genetic levels. **(A)** Yeast bDFE when evolving alone (top) and in the community with the alga (bottom). Each histogram is constructed from data pooled across five replicate BLT experiments. Blue outlines show the bDFE in the other condition. Each colored point indicates the average fitness of a mutant carrying a mutation at the indicated adaptive locus (same data as in panel B). Large CNVs are referred to as chr*x*-*y*n, where *x* is the chromosome number and *y* is the number of copies. (**B)** Fitness of A- and C-mutants measured in competition assays in the A- and C-conditions. Mutants carrying mutations at the 7 most common driver loci are colored. Error bars show ±1 standard error. The solid line indicates the diagonal. (**C)** Distribution of adaptive mutations among A- and C-mutants. Seven most common driver loci are shown individually; all other mutations are grouped into “Other” (see Data S3 for full distributions). Adaptive mutations that are expected to be present in our mutants but were not identified in the genome data are labeled as “Unknown” (see Methods for details). Numbers indicate how many A- and C-mutants carry each type of driver mutation.

The presence of the alga can alter the yeast bDFE by changing which mutations are beneficial (i.e., by imposing a fitness trade-off relative to the A-condition) or by changing the fitness benefits provided by adaptive mutations, or both. To discriminate between these possibilities, we randomly sampled 221 yeast clones from distinct adapted lineages in the A-condition (“A-mutants”) and 189 yeast clones from distinct adapted lineages in the C-condition (“C-mutants”). Clones were sampled at cycle nine, a time point at which most adapted lineages are expected to still be driven by a single beneficial mutation (Methods). We then used competition assays to measure the fitness of all A- and C-mutants relative to their ancestor in both A- and C-conditions (Methods; Figures S9–S11; Data S2). These direct measurements of competitive fitness are concordant with our estimates from the BLT experiment (see Supplementary Information; Figure S12). We found that the C-mutants had significantly higher fitness in their “home” C-condition than A-mutants, by on average 10.8% (95% CI [7.7%, 14.0%], *P* = 2×10^−11^, ANOVA model: fitness ~ environment, *F* = 45.83, *df* = 1) and that A-mutants were more fit in their home A-condition than C-mutants by on average 14.4% (95% CI [10.2%, 18.5%], *P* = 2×10^−11^; *F* = 45.42, *df* = 1), consistent with a signature of local adaptation. However, the fitness distributions of both A- and C-mutants are wide and overlapping in both conditions, so that some A-mutants are more fit than some C-mutants in the C-condition and vice versa.

Interactions with the alga significantly alter the fitness of 88% (362/410) of all sampled mutants (false discovery rate (FDR) = 11%, obtained by permutation). However, fitness is positively correlated between the two conditions across all mutants (Figure 2B; Pearson *R* = 0.70, 95% CI [0.65, 0.75], two-sided *P* = 10^−63^, *t* = 19.9, *df* = 408). Importantly, none of the A- or C-mutations are deleterious in their “non-home” condition. Thus, the presence of the alga changes the fitness benefits provided by adaptive mutations in yeast but does not impose a measurable fitness trade-off, in the sense that it does not alter which mutations are beneficial.

### The presence of the alga alters the distribution of mutations contending for fixation in yeast

Given that all sampled yeast mutants are beneficial both in the presence and in the absence of the alga and that mutant fitness is correlated between the two conditions, we expected that the sets of A- and C-mutants would be genetically indistinguishable. To test this expectation, we sequenced the genomes of 181 out of 189 C-mutants, 215 out of 221 A-mutants, as well as 24 ancestral isolates as controls (Methods; Figures S13–S16; Data S3). We found 176 large copy-number variants (CNVs) across 14 loci in the genomes of A- and C-mutants and none in the ancestral isolates. All of these large CNVs are thus likely adaptive (see Supplementary Information for an extended discussion), with a typical A- and C-mutant carrying on average 0.39 ± 0.04 and 0.51 ± 0.04 of these mutations, respectively. 85/176 large CNVs are whole-chromosome aneuploidies. Out of 91 remaining ones, 64 are partial losses on Chr IV, with breakpoints concordant with known LTR elements (Data S3). This large number of aneuploidies is consistent with the fact that they occur in *S. cerevisiae* at a rate of about 10^−4^ per diploid genome per generation^44^ and the fact that they can be adaptive in some conditions^45–47^.

In addition to large CNVs, we discovered 185 small indels and point mutations at 63 loci for which mutations are found in A- and C-mutants significantly more often than expected by chance (see Methods and Supplementary Information), suggesting that these small mutations are also adaptive. A typical A- and C-mutant carries on average 0.35 ± 0.03 and 0.60 ± 0.06 of these small adaptive mutations, respectively. Overall, we identified 361 beneficial mutations across 77 loci in 250 out of 396 adapted mutants, with each A- and C-mutant carrying on average 0.74 ± 0.04 and 1.11 ± 0.07 such mutations, respectively (Table S2, Data S3).

We quantified the diversity of adaptive mutations carried by A- and C-mutants with the probability of genetic parallelism, *P_g_*, which is the probability that two random clones share a mutation at the same driver locus (lower *P_g_* values imply higher diversity). We found that *P_g_* is slightly higher among A-mutants than among C-mutants (*P_g_* = 6.0 ± 0.6% and 8.5 ± 0.8%, respectively), although this difference was not statistically significant (*P* = 0.06, two-sided permutation test; see Methods). Nevertheless, there were large differences in the frequency distribution of driver mutations among A- and C-mutants (Figures 2C and Extended Data Figure 4, Data S4; *P* = 5×10^−8^, two-sided χ^2^-test, χ^2^ = 160.51, *df* = 76), suggesting that the chance for any given beneficial mutation to rise to a high enough frequency and be sampled varied dramatically between A- and C-conditions. The starkest difference was observed for the amplification of chromosome XIV (mutation chrXIV-3n): 10% (18/181) of C-mutants carried it but none of the 215 A-mutants did (Figures 2C and Extended Data Figure 4), despite chrXIV-3n mutations being beneficial in both conditions (Figure 2A and B).

This initially puzzling observation could be explained by the dynamics of adaptation in populations evolving in the clonal interference regime^42,48,49^. Clonal interference prevents weak and/or rare beneficial mutations from reaching even moderately high frequencies^32,42,50,51^. Thus, the same beneficial mutation can have dramatically different chances of being sampled in the two conditions if it is located in different parts of the respective bDFEs. Consistent with this explanation, we find that a typical chrXIV-3n mutant is 2.05 ± 0.04 times more fit per cycle than the ancestral wild-type and ranks in the top 11 ± 1.3% of most fit mutants in the C-condition, while being only 1.59 ± 0.03 times more fit than the ancestor and ranking at 53 ± 5.3% in the A-condition (Figure 2A). Other mutations with strong discrepancies in their representation among A- and C-mutants show similar shifts in their fitness effects and rank order (Figure 2A, Data S4). In general, interactions with the alga shifted the fitness ranks of mutants between conditions by 14.1 ± 0.8% on average. We used simulations to confirm that such differences in fitness effect and rank are sufficient to explain the observed genetic differences between A- and C-mutants (see Supplementary Information, Extended Data Figure 5).

The fact that the sets of A- and C-mutants are genetically different implies that the yeast populations in these two conditions are about to embark on distinct evolutionary trajectories. Indeed, A- and C-mutants primarily represent high-frequency lineages in the respective populations. Therefore, the mutations that they carry are more likely to win the clonal competition towards fixation in the condition from which they were sampled. Then, the fact that A- and C-mutants carry statistically distinct sets of mutations implies that mutations contending for fixation in yeast in the A-versus C-conditions are also different. In other words, by altering the fitness benefits of mutations, the ecological interactions with the alga change the evolutionary trajectory of yeast, at least over the short-term.

### Adaptive mutations in yeast have diverse ecological consequences for the community

A mutation that spreads and fixes in the yeast population could subsequently alter the ecological dynamics of the yeast-alga community; and adaptive mutations at different loci may have different ecological consequences. For example, some mutations could increase yeast’s competitive ability and ultimately lead to the exclusion of the alga from the community. Others could increase yeast’s cooperativity and thereby strengthen the mutualism. To assess the prevalence and the magnitude of different ecological effects of adaptive mutations in yeast, we selected 28 C-mutants and 31 A-mutants that are representative of the genetic diversity of contending mutations (Methods). We formed 59 “mutant communities” by culturing each of these yeast mutants with the ancestral strain of the alga. As all A-mutants and C-mutants are adaptive in both the A- and C-conditions, we pool all of them to increase our power to identify their ecological effects. We quantified the ecological effect of each mutation by measuring YYC (yeast yield in community) and AYC (algal yield in community), as well as YYA (yeast yield alone; Methods, Figure S18, S19). As before, we define yield as the species density at the end of the 5-day growth cycle. We focused on yields for two reasons. Yield is a measure of absolute species abundance which determines the robustness of our community to demographic fluctuations: communities with higher yields of both species are more ecologically stable^3^. In addition, as discussed above, the ratio of YYC to YYA as well as the ratio of AYC to AYA (alga yield alone) inform us about the net balance between competition versus cooperation in our communities.

We found that many of the adaptive mutations significantly affected both YYA and YYC (Figure 3A, S19A,B). The majority of tested mutations, 53% (31/59), significantly decrease YYA while 15% (9/59) of them significantly increase it (two-sided FDR = 28%, see Methods). At the same time, we found no mutations that significantly decreased YYC, but 34% (20/59) of them significantly increased it (FDR = 26%). Mutations have uncorrelated effects on YYA and YYC (Figure 3A), suggesting that yeast yield in the two conditions is determined by different underlying traits. Mutations in yeast also significantly alter AYC, with 24% (14/59) of mutations increasing it and 8% (5/59) decreasing it (FDR = 32%), with the effects on AYC and YYC being positively correlated (Figures 3B and S19C).

**Figure 3.**
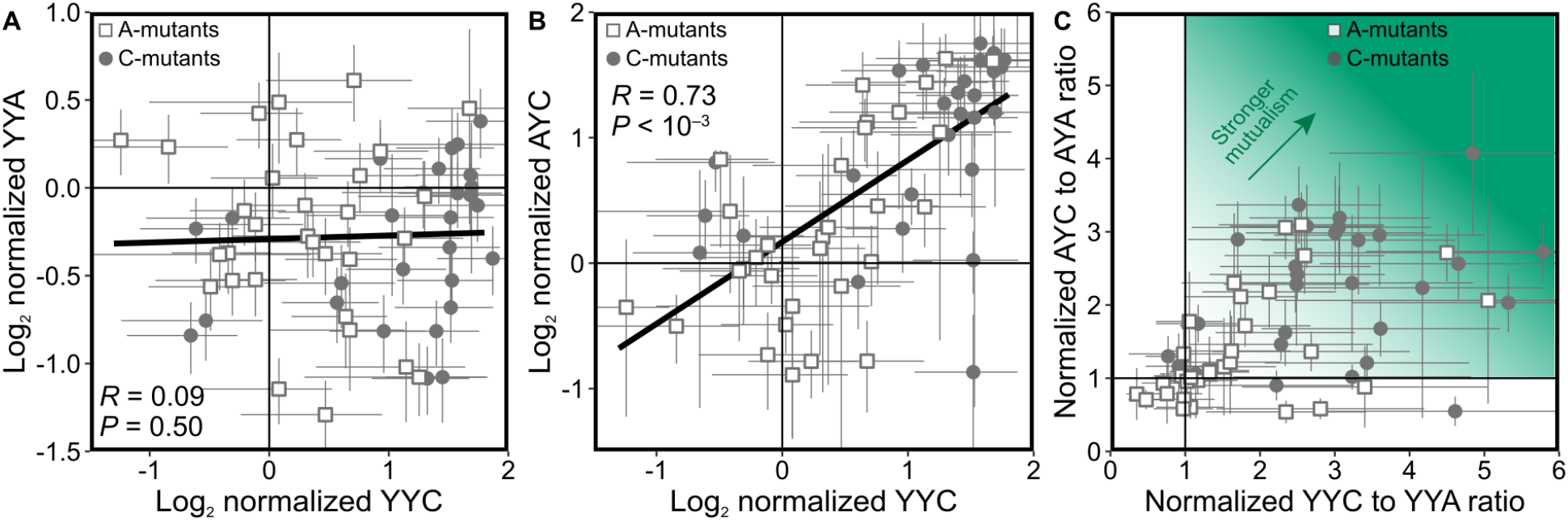
Adaptive mutations in yeast have diverse ecological consequences. **(A)** Correlation between yeast yield in community (YYC) and yeast yield alone (YYA), normalized by ancestral values, across sampled mutants. (**B**) Correlation between yeast yield in community (YYC) and alga yield in community (AYC), normalized by ancestral values, across sampled mutants. (**C)** The ratio of YYC to YYA and the ratio of AYC to AYA (algal yield alone), normalized by ancestral values, across sampled mutants. Darker shades of green indicate stronger mutualism. In all panels, error bars represent ±1 standard error of the mean. In panels A and B, *P*-values are obtained by permutation (Methods) and solid lines are fitted by linear regression.

The fact that mutations alter the yields of both species suggests that some of them may also tip the balance between cooperation and competition in one or the other direction. Indeed, we found that 24% (14/59) of mutants have significantly increased both AYC/AYA and YYC/YYA ratios, 39% (23/59) have a significantly increased the YYC/YYA ratio only, and 12% (7/59) have significantly decreased one or both ratios (two-tailed FDR = 5%, Figure 3C). Thus, seven mutants acquired mutations that weaken the yeast-algal mutualism, at least for one of the partners, and 37 mutants acquired mutations that enhance the mutualism.

These results show that yeast has access to beneficial mutations with ecologically diverse consequences and suggest that our community has the potential to embark on a variety of eco-evolutionary trajectories with possibly different ecological outcomes.

### Mutations favored by selection in the presence of the alga strengthen the mutualism

Given that adaptive mutations in yeast have a variety of ecological consequences and that yeast populations in the absence or presence of the alga are likely destined to fix different adaptive mutations, we next asked whether mutations that contend for fixation in the A- or C-conditions might have systematically different ecological effects.

We first noticed that the C-mutants clustered in the top right corner of the YYC versus AYC plot (Figure 3B). A formal statistical test confirmed that YYC and AYC values for the C-mutant communities were on average 103% and 79% higher than those for the A-mutant communities, respectively (YYC: 95% CI [62%, 143%], *P* = 3×10^−4^, permutation test; AYC: 95% CI [35%, 122%]; *P* = 4×10^−3^; *n*_1_ = 28 C-mutants and *n*_2_ = 31 A-mutants). These differences remained large and significant even after accounting for the frequencies with which different driver mutations were observed in our yeast populations (YYC: *P* = 10^−5^; AYC: *P* = 3×10^−3^, permutation test, Extended Data Figure 6, see Methods), indicating that this trend is not an accidental byproduct of our choice of A- and C-mutants. Instead, the observed differences in yield must be caused by systematic genetic differences between the A- and C-mutants. In other words, in the presence of the alga, natural selection favors yeast mutants that produce higher yields in the community. This conclusion is further corroborated by the fact that both YYC and AYC are correlated strongly and significantly with competitive fitness in the C-condition, but only weakly with fitness in the A-condition (Extended Data Figure 7).

A mutation in yeast that increases YYC may concomitantly lead to either a gain or a loss of YYA (Figure 3A). Therefore, such a mutation may either increase or decrease the net benefit that yeast derives from its interactions with the alga over the growth cycle. We found that 86% (24/28) of the C-mutants had a significantly higher YYC/YYA ratio than the ancestor (FDR = 21%), of which many, 32% (9/28), had also a significantly higher AYC/AYA ratio (FDR = 6%; Figure 3C) but only 1/28 had a significantly lower AYC/AYA ratio (FDR = 58%). In contrast, a smaller fraction, only 52% (16/31), of the A-mutants had a higher YYC/YYA ratio (FDR = 35%), of which 16% (5/31) also had a higher AYC/AYA (FDR = 12%) and 2/31 had lower AYC/AYA (FDR = 29%). Thus, C-mutants both benefit more often from the presence of the alga and reciprocally more often provide benefits to the alga compared to the A-mutants. In other words, adaptive mutations that dominate yeast adaptation in the presence of the alga are more likely to make yeast more cooperative and/or less competitive and thereby strengthen the mutualism, compared to mutations that dominate adaptation in the absence of the alga.

An interesting potential consequence of this shift in selection on yeast precipitated by the presence of the alga is that it can change the repeatability of yeast evolution along the competition-mutualism continuum. We quantified such ecological repeatability by the probability that two randomly drawn yeast mutants that contend for fixation in a given condition both increase or both decrease the YYC/YYA ratio and simultaneously both increase or both decrease the AYC/AYA ratio. The probability of ecological parallelism would be 25% under a uniform null model. For the A-mutants, this probability is 33 ± 2.8%, indistinguishable from the null expectation (*P* = 0.21, two-sided χ^2^-test, χ^2^ = 1.55, *df* = 1). In contrast, it is 68 ± 5.5% for the C-mutants, which is significantly higher than expected (*P* = 6×10^−8^, χ^2^ = 29.3, *df* = 1) and also significantly higher than for the A-mutants (*P* = 0.031, two-sided permutation test; see Methods). Thus, yeast evolves more repeatably (towards stronger mutualism) in the presence of the alga than in its absence.

In summary, our results show that mutations contending for fixation in yeast populations evolving alone have relatively diverse effects on the ecology of the yeast-algal community, with some strengthening and some weakening the mutualism. In contrast, mutations contending for fixation in yeast evolving in the presence of the alga predominantly lead to higher yields of both species, which strengthens the yeast-alga mutualism and makes evolution more repeatable at the ecological level.

### Mutualism enhancement is not selected directly but is likely a byproduct of selection for other yeast life-history traits

We next asked how stronger mutualism could possibly evolve in our community. Specifically, does natural selection in the presence of the alga favor mutations that increase cooperativity and/or decrease competitiveness in yeast directly or is this bias a byproduct of selection for other traits^25,52^? Natural selection can directly favor rare mutualism-enhancing (i.e., more cooperative and/or less competitive) yeast mutants only if such mutants preferentially receive fitness benefits from their algal partners, that is, if there is a partner-fidelity feedback^53^. Our system is well-mixed, so that all diffusible benefits are shared by the entire culture, eliminating any potential fitness advantage of rare mutualism-enhancing mutants^25,54^. The only way to prevent such diffusion and ensure preferential benefit exchange with an algal partner is for such a mutant to form a physical association with the partner^21^. However, we found no evidence for such associations in any of the sampled mutants (Extended Data Figure 8; Methods). Given the absence of a plausible partner-fidelity feedback, the increased cooperativity and/or decreased competitiveness of the C-mutants must be a byproduct (pleiotropic effect) of selection for one or more other traits.

We sought to identify traits under selection in the C-condition that could cause yields to increase. Both competitive fitness and yield depend on fundamental physiological and life-history traits embodied by yeast and alga, such as their growth rates, mortalities, nutrient consumption efficiencies, etc^55^. Since measuring all potentially relevant traits was not feasible in this study, we focused on two key traits that are known to be under selection in environments with variable nutrient availability. The maximum population growth rate, *r*, is important for competitive fitness when resources are abundant^56,57^, a condition that takes place at the beginning of each growth cycle in our cultures. The carrying capacity, *K*, is an indicator of nutrient utilization efficiency, which is important for competitive fitness when resources are scarce^56,57^, a condition that takes place at the later phase of each growth cycle. We estimate *r* and *K* in the A-condition, reasoning that these intrinsic traits would be relevant for fitness and yield in both A- and C-conditions. We estimate *r* by regressing the natural logarithm of the yeast cell density against time during the initial phase of the growth cycle (Methods). We estimate *K* as the maximum yeast cell density during the growth cycle, which is usually achieved on day 2.

We estimated *r* and *K* for all 59 sampled mutants (Figures S20, S21; Methods) and found that many mutations significantly increased and decreased either one or both traits (Figure 4A). We found a negative correlation between the effects of mutations on *r* and *K* (Pearson *R* = –0.38, 95% CI: [–0.59, –0.13], two-sided permutation *P* = 0.004, Figure 4A), indicating a trade-off between growth rate and nutrient utilization efficiency, which is often observed in other systems^58–65^. More specifically, we found 16 C-mutants and 4 A-mutants have a significantly higher *K* and a significantly lower *r* than the ancestor (FDR = 18%), an observation that is rare in experimental evolution studies^60^ where selection usually favors higher *r^55,59,66–69^*. However, theory suggests that high-*K*/low-*r* mutations can be favored in the presence of an *r*-*K* trade-off in populations near starvation^56,57^. We confirmed that 60% (12/20) of our significant high-*K*/low-*r* mutants can in fact invade the ancestral yeast population in simulations of a logistic growth model (see Supplementary Information and Figure S22). While this model demonstrates the plausibility of selection favoring high-*K*/low-*r* mutants, it does not capture all the important complexities of our system. Thus, we next explicitly tested whether *r* and *K* are under selection in our A- and C-conditions. To this end, we examined the correlation between these traits and competitive fitness among all 59 assayed mutants.

**Figure 4.**
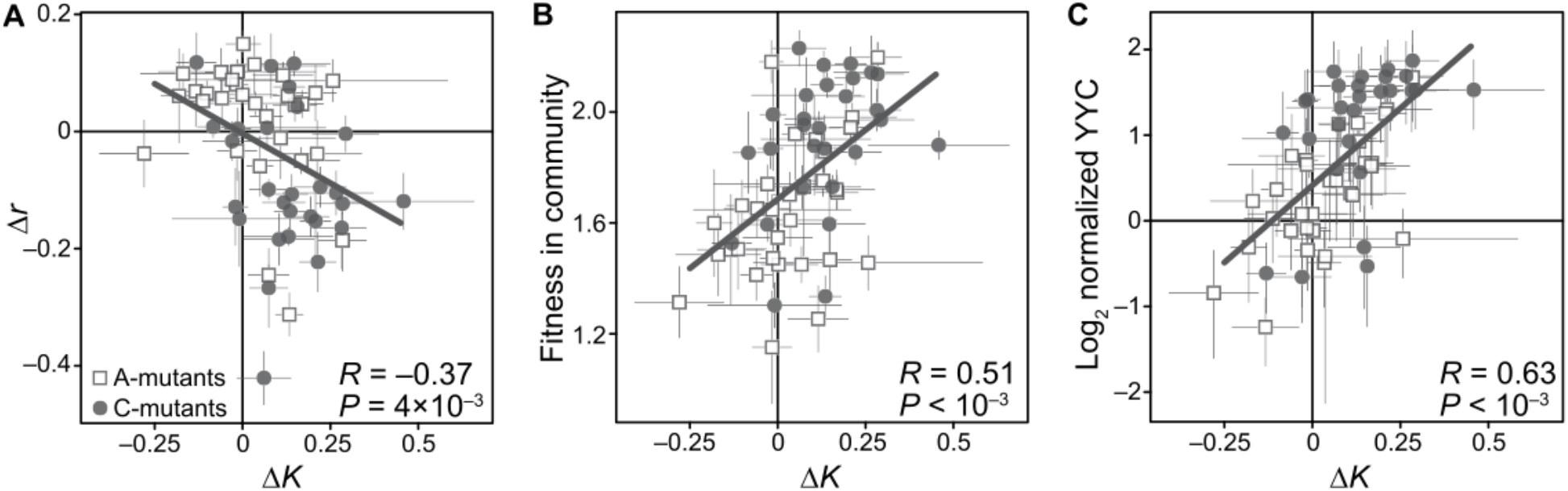
The effects of adaptive mutations on life-history traits, fitness and yield. The effects of mutations on growth rate (*r*) and peak density (*K*) in the A-condition, Δ*r* and Δ*K*, are reported as fractional differences relative to the ancestral values (Methods). **(A)** Correlation between Δ*K* and Δ*r* among all sampled adapted mutants. (**B)** Correlation between Δ*K* and competitive fitness in the community among all sampled adapted mutants. (**C)** Correlation between Δ*K* and YYC (yeast yield in community), normalized to ancestral values, among all sampled adapted mutants. Error bars show ±1 standard error of the mean. Solid lines are fitted by linear regression. *P*-values are computed by permutation (Methods).

We found that neither *r* nor *K* are significantly correlated with fitness in the A-condition (Supplementary Information; Extended Data Figures 9B,10A; Table S4). As a result, the A-mutants have *r* and *K* values that are indistinguishable from the ancestor (average Δ*r* = 2 ± 1.9%, *P* = 0.27; average Δ*K* = 3 ± 2.4%; *P* = 0.29; *n* = 31, permutation test). These observations suggest that other traits that we have not measured must be more important for fitness in the A-condition than either *r* or *K*. In contrast, fitness in the C-condition is positively correlated with *K* (Figure 4B, *R* = 0.51, 95% CI [0.29, 0.68], *P* < 10^−4^) and negatively correlated with *r* (Extended Data Figure 9A, Pearson’s *R* = –0.51, 95% CI [–0.68, –0.29], *P* < 10^−4^), consistent with the observed *r*-*K* trade-off, and both of these traits together explain 37% of variation in competitive fitness in the C-condition. Interestingly, a negative correlation between fitness and *r* persists even after controlling for *K* (Table S4), suggesting that other unmeasured traits must also be important for fitness in the C-condition. Regardless, C-mutants reach *K* values on average 12.7% higher than the ancestor (*P* = 4×10^−4^, permutation test) and 9.5% higher than A-mutants (95% CI [2.6%, 16.3%], *P* = 0.03). A typical C-mutant also has a significantly lower *r* than both the ancestor (average Δ*r* = –8.7 ± 2.3%, *P* = 0.001) and a typical A-mutant (Δ*r* = –11%, 95% CI [– 17.0%, –5.1%], *P* = 7×10^−4^). These observations suggest that nutrient efficiency is an important component of fitness in the C-condition, and that yeast high-*K*/low-*r* mutants are favored by selection in the presence of the alga.

We next asked whether higher-*K* mutants achieve higher yields. We might expect a strong positive correlation between these quantities because, all else being equal, mutants that reach higher density in the middle of the growth cycle due to their higher carrying capacity are more likely to maintain higher density at the end of the growth cycle. However, we found no correlation between *K* and YYA (Extended Data Figure 10C). This lack of correlation further confirms that adaptive mutations that we sampled must affect other unmeasured traits which are more important for yield than *K*. In contrast, we found that *K* and YYC were positively correlated (Figure 4C), suggesting that higher nutrient efficiency is important for achieving higher yields in the community with the alga.

Our observations suggest a plausible model for how adaptive evolution can favor mutualism enhancement in the absence of partner-fidelity feedbacks: ecological interactions with the alga intensify selection for yeast mutants that use resources more efficiently (i.e., those that reach higher *K* even at the expense of reduced *r*); once these mutants spread in the yeast population, they support higher yields of both members of the community. Whether mutualistic partners generally induce selection for lower *r* and/or higher *K*, and whether such selection consistently leads to increased yields of both species remains an open question.

Similar to our analysis of ecological parallelism, we asked whether the presence of the alga alters the probability of parallelism at the level of life-history traits, which we define as the probability that *r* would be affected in the same direction in two randomly sampled mutants and that *K* would also be affected in the same direction in these mutants (see Methods). We find that the probability of trait parallelism is 30.7 ± 1.7% for the A-mutants, which is not significantly different from 25% expected under the uniform null model (*P* = 0.47, two-sided χ^2^-test, χ^2^ = 0.53, *df* = 1). In contrast, the probability of trait parallelism is 46.3 ± 4.8% for the C-mutants, which significantly exceeds 25% (*P* = 0.009, χ^2^ = 6.7, *df* = 1), suggesting that evolution in the presence of the alga becomes more repeatable not only at the ecological level, as shown in the previous section, but also at the level of underlying life-history traits.

In summary, our results show that interactions with alga shift natural selection on yeast to favor mutants that increase *K* and decrease *r*, which in turn leads to increasing yields of both species in the community. The shift in selection imposed by the alga makes evolution more repeatable both at the level of life-history traits and even more so at the ecological level.

## Discussion

We characterized early adaptation in the experimental yeast-alga community and made three main observations. First, we found that yeast have access to adaptive mutations that are not only genetically diverse but also have diverse ecological effects. Second, even though there are no measurable fitness trade-offs for yeast between growing alone or with the alga, the presence of the alga modifies the fitness benefits provided by many mutations. This shift in selection pressures is sufficient to change the set of mutations that contend for fixation in yeast and thereby to alter the course of its evolution. Third, mutations that are strongly favored by selection in the presence versus absence of the alga have different ecological consequences. Specifically, the presence of the alga shifts selection on yeast to favor mutations that enhance the yeast-alga mutualism (as measured by the yield of both species at the end of the growth cycle), making evolution at the ecological level more repeatable.

Insofar as our yeast-alga community is representative of other ecological communities, our results suggest that (i) organisms have access to a variety of adaptive mutations with diverse ecological consequences and (ii) ecological perturbations, such as removal or addition of species, can change the fitness effects of many of these mutations, thereby altering future outcomes of evolution not only at the genetic but also at the ecological level. As a result, the eco-evolutionary dynamics of multi-species communities are likely historically contingent on both prior evolution^70^ and ecology^71^. Thus, we might expect that ecological communities would generically have the potential to embark on a variety of divergent eco-evolutionary trajectories and approach different ecological attractors. For example, mutations that are beneficial to yeast in our community can either increase or decrease the yields of both species suggesting that our community has the potential to evolve either towards stronger mutualism or towards mutualism breakdown, with probabilities of these outcomes being dependent on whether yeast previously evolved in the presence or absence of the alga.

Given this potential for ecological and evolutionary historical contingency, one might expect *a priori* that replicate communities would often diverge towards different ecological states. However, recent laboratory studies have found that replicate communities tend to evolve towards similar ecological states with notable repeatability^13,15,16,20,21,24,25,72–76^. Our results show that, while yeast has access to a set of adaptive mutations that are quite diverse in terms of their ecological effects, natural selection acting on yeast growing in the community strongly favors a biased subset of these mutations, namely those that produce higher yields of both yeast and alga. When viewed in the context of these prior observations, our findings suggest that ecological interactions may limit the space of the most likely evolutionary trajectories. In our system in particular, the presence of the alga modifies the effects of mutations in yeast in such a way that yeast evolution becomes more repeatable at the ecological level, at least over the short-term. In other words, ecological interactions may canalize evolution. Whether such canalization is a general feature of evolution in a community context remains to be determined.

In our competitive mutualistic community, canalization appears to occur in the direction of enhanced mutualism in the sense that the presence of the alga shifts selection on yeast in favor of mutations that benefit both species. There are no demonstrated mechanisms that would favor such enhanced mutualism in our community directly, but our results suggest another plausible scenario for how it can evolve. Mutations in yeast favored in the presence of the alga tend to increase yeast’s carrying capacity in our medium and reduce its growth rate. Increased carrying capacity could provide the competitive advantage necessary for such mutants to spread. Once these mutations dominate, increased *K* and/or decreased *r* could enhance cooperation or reduce competition with the alga. Specifically, increased *K* implies that there are more yeast cells to generate CO_2_, which stimulates algal growth. Reduction in *r* could also benefit the alga via the “competitive restraint” mechanism^77^ in which slower growing yeast compete less for the initial supply of ammonium and thereby offer the alga an opportunity to grow more and supply more ammonium during the latter portions of the growth cycle. However, competition for the initial ammonium can probably not be reduced to zero solely by mutations in yeast because yeast lacks the molecular machinery for metabolizing the only other nitrogen source, nitrite. Therefore, a single mutation or even a few mutations cannot alleviate yeast’s basic requirement for ammonium. Furthemore, traits other than *r* and *K* most certainly contribute to both fitness and yield. Thus, additional experiments will be needed to determine how adaptive mutations in yeast modify the competitive and cooperative phases of the growth cycle to provide an evolutionary advantage and increase the yields of both species.

How the presence of the alga amplifies the fitness advantage of high-*K* mutants is currently unclear. An analysis of the genetic and biochemical basis of yeast adaptation may help us answer this question and assess how general the ecological mechanisms of mutualism enhancement might be. However, one challenge is that many mutations driving adaptation in yeast are large chromosomal amplifications and deletions, and it is unclear which amplified/deleted genes actually cause the fitness gains and changes in the ecologically relevant traits. At this point, we can only speculate on this subject. For example, it is known that ChrXIV-3n amplifications are adaptive under ammonium limitation, possibly driven by the copy number of the gene *MEP2* that encodes a high affinity ammonium transporter^78^. We suspect that these adaptations are particularly beneficial to yeast in the C-condition because the alga provides a continuous but low flux of ammonia. Another interesting example are mutations in genes *HEM1, HEM2* and *HEM3*, which provide much larger fitness benefits in the C-condition compared to the A-condition (Data S4) possibly because they shift the metabolic balance towards fermentation at higher concentrations of dissolved oxygen produced by the alga (see Supplementary Information). Elucidating these and other mechanisms of physiological adaptation in our competitive mutualistic systems is the subject of future work.

To conclude, our results suggest that microbial adaptation in the community context is driven by many mutations that are genetically and phenotypically diverse and have diverse ecological consequences. Changes in the ecological milieu, such as loss of some species or invasions by others, may not necessarily alter which mutations are beneficial to community members. Nevertheless, such ecological changes can quantitatively alter the benefits of mutations, so that evolutionary trajectories become canalized towards certain ecological outcomes.

## Methods

### Barcode lineage tracking (BLT) experiment and data analysis

#### Strains

We used the strain CC1690 of the alga *Chlamydomonas reinhardtii*, which can also be obtained from the Chlamydomonas Resource Center. The barcoded library of the diploid yeast Saccharomyces cerevisiae strain GSY6699^40^ was kindly provided by Prof. Gavin Sherlock. This is a diploid, prototrophic strain derived from the BY genetic background, homozygous throughout the genome, except for locus YBR209W, where one copy of a DNA barcode was integrated^37^. Our starting library consists of about 5 × 10^5^ clones, each of which carries a unique DNA barcode at this locus. In principle, the genomes of all clones should be identical everywhere else prior to our barcode lineage tracking (BLT) experiment. However, as discussed in Supplementary Information (Sections 1.3 and 3), we found that our initial population already contains some pre-existing polymorphisms, which arose prior to our BLT experiment.

#### Growth conditions

Both yeast monocultures and yeast-alga communities were cultured in a defined minimal medium^33^ (“CYM medium”) supplemented with 2% dextrose, 10mM KNO_2_ and 0.5mM NH_4_Cl, which we thereafter refer to as the “growth medium”. All cultures were grown in 10mL of the growth medium in 50mL flasks (FisherSci #FS2650050) capped with 50mL plastic beakers (VWR #414004-145) at room temperature (21°C) on a platform shaker with 70 foot-candles of constant light (three Feit Electric #73985 suspended approximately 24 inches above the platform shaker) shaking at 125 RPM, unless noted otherwise.

#### BLT pre-cultures

Prior to the BLT experiment, yeast and alga were pre-cultured in 50mL of growth medium in 250mL delong baffled flasks (PYREX #C4446250) for two and 10 days respectively. Alga pre-cultures were started from colonies. To start yeast pre-cultures, the barcoded yeast library was thawed from frozen stock at room temperature, then 500μL were transferred into 50mL of the growth medium.

#### BLT initiation and propagation

We conducted five replicate BLT experiments for each of two treatments, yeast monoculture (the A-condition) and yeast + algae community (the C-condition). Each monoculture BLT experiment was initiated from 100μL of the yeast pre-culture. Each community BLT experiment was initiated from 100μL of the 1:1 (v/v) yeast and alga mixture. Cultures were grown for 5 days before being diluted 1:100 for the next growth cycle (100μL into 10mL fresh media). A total of 17 growth/dilution cycles were completed. A detailed discussion on the number of generations per growth cycle is provided in Supplementary Information (Section 1.1). Throughout this work, we ignore adaptation in the alga, as discussed in Supplementary Information (Section 1.2).

#### Culture preservation

Glycerol stocks were taken of the yeast pre-culture and yeast + algae inoculum mixture, as well as at the end of every odd growth cycle. Separate stocks were stored for DNA extraction and cell isolation purposes with two replicates each, for a total of 4 stocks per culture per time point. Cell isolation stocks were created by aliquoting 1.5mL of culture into 500μL of 80% glycerol, mixing by vortex and storing at −80°C. DNA stocks were created by removing the supernatant of the remaining 7mL of culture via centrifugation and resuspending in 2mL of 20% glycerol (80% glycerol diluted with 1x PBS), which was then stored as two separate 1mL stocks at −80°C.

#### DNA isolation

DNA stocks were thawed and DNA isolated using a “salting out” method, based on established protocols^79,80^. The thawed stocks were first centrifuged, and the supernatant removed. The pellet was resuspended in 300μL 3% SET buffer (3% SDS, 10mM EDTA, 30mM Tris) and incubated at 65°C for 15 minutes. The tube was cooled to room temperature by immersing it in room temperature water, then 2.5μg of RNAse A was added. After vortexing, the mixture was incubated at 37°C for 1 hour, after which it was cooled on ice. 150μL of 3M Sodium Acetate was then added and mixed by inversion, after which it was cooled on ice for a further 5 minutes before centrifuging at maximum speed for 10 minutes on a tabletop centrifuge. The supernatant was transferred to a new tube and DNA was precipitated by the addition of 500μL isopropanol, which was mixed by inversion and then pelleted by centrifugation for 1 minute at maximum speed. The supernatant was removed and the pellet was washed with 200μL cold 70% ethanol without vortexing before being allowed to dry inverted for 30 minutes at room temperature before the DNA was resuspended in 50μL molecular biology grade water.

#### Sequencing library preparation

The barcode locus was amplified through a 2-step PCR protocol slightly modified from Ref. ^38^. The first amplification added inline indices for sample multiplexing, universal molecular identifiers for removing PCR duplicates during analysis and Illumina-compatible adapter sequences for a second round of amplification with standard Illumina Nextera XT primers. For the first reaction, 10μL of template was mixed with 25μL of OneTaq 2x Master Mix, 1μL of 25mM MgCl_2_, 1μL each of the forward and reverse primers (at 10mM concentration) and 12μL of molecular biology grade water. Primer sequences are as described^38^. This mixture was amplified using the following conditions: (1) 94°C for 10 min; (2) 94°C for 3 min; (3) 55°C for 1 min; (4) 68°C for 1 min; (5) Repeat steps 2–4 for a total of 8 cycles; (6) 68°C for 1 min; (7) Hold at 4°C. The amplified product was purified using Ampure XP magnetic beads using established protocols (with 50μL of beads used per sample). Then, 10μL of the purified product was used as the template for a second reaction along with Illumina Nextera XT primers (1μL of 10mM stock for each primer), 1μL of 25mM MgCl_2_, 25μL of OneTaq 2x Master Mix and 12μL of water with the following reaction conditions: (1) 94°C for 5 min; (2) 94°C for 30 sec; (3) 62°C for 30 sec; (4) 68°C for 30 sec; (5) Repeat steps 2–4 for a total of 25 cycles; (6) 68°C for 5 min; (7) Hold at 4°C. The PCR products were purified using Ampure XP beads as before. 5μL of each sample was mixed to form a pool for Illumina sequencing, concentrated using Ampure beads as before with equal volume of beads as a pooled sample and size-selected via agarose gel extraction to isolate the correct amplicon before submitting for sequencing.

#### Sequencing

Populations A1 and C1 were initially sequenced on a MiSeq platform. All populations (including A1 and C1) were then also sequenced on a HiSeq platform. Data from both runs were combined for all downstream analysis of frequency trajectories. We obtained an average of 2.6 million paired-end reads per time point.

To identify and count DNA barcodes, we used a custom python pipeline BarcodeCounter2 available at https://github.com/sandeepvenkataram/BarcodeCounter2. The package first uses the BLASTn tool to identify sequences known to flank the barcode region within each read pair. If reads contain inline indices, samples can be demultiplexed. Universal molecular identifier (UMI) sequences can be extracted if present within the reads. If a sequence contains multiple barcode regions, these extracted regions are concatenated together. To account for sequencing errors, DNAClust^81^ is then used to cluster the concatenated barcodes into clusters of nearly identical sequences which presumably originated from the same DNA molecule. The output of DNAClust is a FASTA database of all unique barcode sequences present in the library. Then, BWA^82^ is used to map the barcode sequences from each sample onto the clustered FASTA database. Mapping of reads is necessary because the clustering process removes identifying information associating barcode sequences with samples from which they came. PCR duplicates are removed based on the UMI sequences, and the total number of unique reads corresponding to every barcode in the FASTA database is counted. The final output is a tab-delimited table of the read counts for every barcode in every sample. The software is designed to be user-friendly and highly customizable, with simple text files describing the input files, multiplexed samples and a sequence template describing the structure of the sequenced reads. The package is built using python3, and uses the popular BioPython package. The package has multithreading support, and can be run on both personal computers and supercomputing clusters.

When generating the database of all unique barcode sequences, we clustered sequences at 95% similarity, so that, given that the length of our barcode is 52bp, sequences with 3 or more base-pair differences were merged into the same cluster. This set of clustered sequences was generated only once using all of the time points from all sequenced populations.

We developed an iterative heuristic procedure to identify adaptive lineages from lineage tracking data and estimate their fitness.

1. **Initialization.** Neutral barcodes for iteration 1 are identified separately for every pair of consecutive time points. For a given time point pair, we define the set of neutral lineages at iteration 1 as those lineages whose frequency (a) does not exceed 10^−4^ at the earlier time point and (b) increases by less than 100-fold between cycle 1 and cycle 11.
2. **Estimation of mean fitness.** Given the set of neutral lineages at iteration *k*–1, we obtain their total frequency 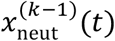 at each cycle *t*. The frequency of neutral lineages is governed by equation 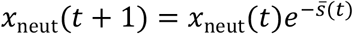. where 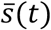 is the mean selection coefficient (per cycle) of the population at cycle *t*. Thus, we estimate the mean selection coefficient of the population at cycle *t* at iteration *k* as

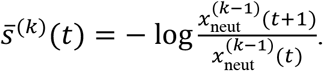 The mean fitness at cycle *t* is then defined as 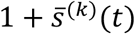.
3. **Estimation of fitness of individual lineages.** The frequency *x_i_*(*t*) of lineage *i* with selection coefficient *s_i_* (per cycle) relative to the ancestor is governed by the equation 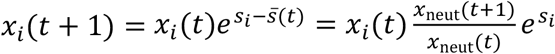. Thus, for every lineage *i* (including those that were called as neutral at the previous iteration) we estimate 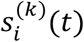 at iteration *k* at time point *t* as

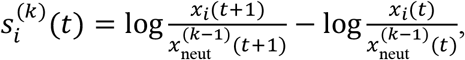

where *x_i_*(*t*) is the observed frequency of lineage *i* at cycle *t*. These estimates are made for every pair of consecutive time points between cycles 1 and 11. If *x_i_*(*t*) = 0 and *x_i_*(*t* + 1) = 0, the time point pair is excluded from the calculations. If only one of the two frequencies is 0, then this frequency is set to 0.5 / total read depth. To obtain the final estimate of the selection coefficient 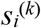 for lineage *i* at iteration *k*, we average all 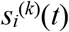, weighting each 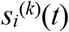 by the number of reads for lineage *i* at time *t*. We calculate the standard error 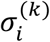 with the same weighting. The relative fitness of lineage *i* is then defined as 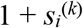.
4. **Calling adapted lineages.** Lineage *i* is called adapted at iteration *k* if 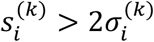.
5. **Updating the set of neutral lineages.** We calculate the coefficient of variation (CV) for each lineage *i* as 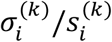. We include any lineage *i* into the new set of neutral lineages at iteration *k* if (a) its maximum frequency does not exceed 10^−4^ between cycles 1 and 11, and (b) either the CV of the lineage exceeds the median CV across all lineages or if 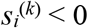 (see Figure S1A).
6. **Termination.** The procedure is terminated when the set of neutral lineages at iteration *k* differs by less than 5% compared to the set at iteration *k* – 1.

Our procedure converges to a stable set of neutral lineages within 5 iterations. We have carried out sensitivity analyses of our procedure with respect to the choice of various parameters, as well as other sanity checks, as described in the Supplementary Information (Section 1.3). Note that we report fitness values relative to the ancestral strain on a per-cycle basis, rather than the per-generation basis typically used in the literature because our cultures experience growth phases other than exponential growth^38,83^ (Figure 1, Supplementary Information (Section 1.1)).

#### Permutation tests for differences in bDFE across conditions

We randomly relabel adapted lineages as being from either the A- or C-condition, and calculate the difference in bDFE median and IQR from this permuted data. This procedure was conducted 10,000 times, to generate a random distribution of median and IQR difference values.

### Competitive fitness assays

We selected cycle 9 to isolate adapted mutants because the estimated fraction of adapted lineages in our evolving populations was large, but each adapted lineage was still at a low frequency (Extended Data Figures 2 and 3). Specifically, 92% of a typical population in the A-condition consisted of adapted lineages, with the median frequency of an individual adapted lineage being 9×10^−5^ (~ 9 cells per lineage at the bottleneck), and 73% of a typical population in the C-condition consisted of adapted lineages, with the median frequency of an individual adapted lineage being 7×10^−5^ (~ 7 cells per lineage at the bottleneck).

#### Isolation of random clones

To isolate adapted clones, frozen stocks of the monoculture and community populations from cycle 9 were thawed, plated onto standard 100mm Petri dishes with CYM + 1% agarose at a dilution of approximately 100 cells per dish, and incubated at 30°C for three days (algae do not grow at 30°C). 88 random colonies were isolated from each population, i.e., a total of 440 clones from the A- and C-condition each. Eight additional clones from each population were harvested at cycle 17 and are present in the pools described below, but they are not included in any of the analyses presented in this study. Each colony was transferred into a well of a 96-well plate (Corning 3370) with 200μL of CYM media and incubated for two days at 30°C. 10μLwas used for each of the “population”, “row” and “column” pools described below. Then, 50μL of 80% glycerol was added to each well, and the plate was stored at −70°C.

#### Barcode genotyping

The DNA barcodes of all isolated clones were identified by sequencing, using the Sudoku method^84,85^. Specifically, 10μL of each clone was pooled into 10 “population” pools (one pool for each source population), eight “row” pools and 12 “column” pools. DNA barcodes in each pool were amplified and sequenced as described above We expect that a given combination of row, column and population pools would have a single barcode in common, defining the isolate in the corresponding well of the appropriate plate. We determined all barcodes present in the intersection of each combinations of row, column population pool. If a single barcode is identified in the intersection, the corresponding well is assigned that barcode identity. If multiple barcodes or no barcode are identified, the associated clone is removed from further analysis.

#### Generation of A, C and N pools

We used the heuristic procedure described above to classify clones with identified barcodes into three groups described below: non-adapted, adapted in the C-condition or adapted in the A-condition. Clones were pooled into three libraries defined by their membership in these three groups as follows. To construct each pool, we transferred 20μL of thawed frozen stock of each isolate into a 2mL 96-deep-well plate (Corning P-2ML-SQ-C) filled with 1.8mL of growth media supplemented to 10mM ammonia. After incubating these plates at 30°C for three days, we formed each pool by combining 200μL of individual saturated cultures. Three pools were stored in 20% glycerol at −70°C.

For the sake of efficiency, clone pooling was based on an earlier version of the heuristic procedure used for the classification of lineages. The current version of the procedure (as described above) classifies lineages slightly differently. As a result, pools do not perfectly correspond to the classification of clones according to the current heuristic procedure, which is provided below. This minor discrepancy has no bearing on our results because the final classification of clones is based on fitness estimates from the competition assays (see below). The *A pool* contains 214 clones that are classified as adapted in A populations. The *C pool* contains 223 clones that are classified as adapted in C populations. The *N pool* contains 144 clones, 84 of which are classified as neutral from the BLT analysis (i.e., not adapted in either A or C populations), 29 clones that are classified as adapted in the A-condition and 31 clones that are classified as adapted in the C-condition. Thus, we measured competitive fitness for a total of 581 clones.

#### Competition assay experiment

To conduct the competitive fitness assays, we pre-cultured each of the three pools (N, A and C; see above) separately in the growth media for two days. We also pre-cultured algae for 10 days, starting from colonies. We then combined A, C and N pools in the 1:1:18 ratio. We carried out three replicate competitions in the A-condition and three replicate competitions in the C-condition. To this end, we inoculated each of the six replicates with 100μL of the combined A/C/N pool. In addition, the three C-condition replicates were inoculated with 100μL of the algae preculture (~ 10^6^ cells / mL).

All replicates were propagated in conditions identical to the BLT experiment for a total of five growth cycles. Glycerol stocks were made at the end of each growth cycle after the dilution step by centrifuging the full culture volume, removing the spent media and resuspending the pellet in 2mL of 20% glycerol + PBS. Two 1mL aliquots of this glycerol suspension were stored at −70°C. One of these aliquots was harvested for DNA extraction and barcode sequencing using protocols described in the section on the heuristic BLT analysis procedure.

#### Competition assay data analysis

Barcodes were identified and counted as described above. The resulting barcode count data were analyzed as described previously^38^ using software available at https://github.com/barcoding-bfa/fitness-assay-python. Briefly, the 84 non-adapted barcodes (as defined from the BLT analysis described above) from the N pool were used to estimate the mean fitness trajectories and the additive and multiplicative noise parameters for each pair of time points in each assay^38^. These estimates were used to estimate the fitness of every lineage for each pair of neighboring time points along with the error in the estimate. The variance of an estimate for a given pair of time points was calculated as the inverse of the read depth at the earlier of the two timepoints + the estimated multiplicative noise parameter. Inverse variance weighting was then used to combine estimates across all time point pairs to generate a single fitness and error estimate for each lineage in each replicate. Replicate estimates were combined using further inverse variance weighting to generate the final fitness estimate for each isolate in the A- and C-conditions. For each mutant in each condition, we also calculated the 95% confidence interval around the fitness estimate based on the variability in fitness measurements between replicates (assuming that measurement errors are distributed normally). Fitness estimates are provided on a per growth cycle basis, as discussed above. Validation of this analysis procedure and additional statistics are described in Supplementary Information (Section 2).

### Genome sequencing and analysis

We sequenced full genomes of 219 A-mutants, 187 C-mutants, 8 non-adapted evolved isolates and 24 ancestral isolates sampled from the inoculum population. Sequencing failed for four A-mutant and six C-mutants, leaving us with 215 A-mutants and 181 C-mutants with sequenced genomes, 8 non-adapted evolved isolates and 24 ancestral isolates (a total of 428 clones). For all remaining clones, we obtained high quality genome data (> 4x coverage, mean coverage of 24x).

DNA extraction and library preparation and sequencing. DNA was extracted using the YeaStar yeast genomic DNA extraction kit Protocol I (Zymo research #D2002) with in-house produced YD Digestion buffer (1% SDS + 50mM Na_2_PO_4_), DNA Wash buffer (80% ethanol + 20mM NaCl) and Elution Buffer (10mM Tris-HCl). 0.2μL of 25mg/mL RNAse A (Zymo research #E1008-8) was used for each sample, as well as 1 Unit (0.2μL of 5U/μL stock solution) of Zymolyase (Zymo research #E1004). Libraries were prepared using the method described by Baym et al^86^ and sequenced on the Illumina HiSeq4000 platform. Sequencing services were provided by Novogene Inc. and the UCSD Institute for Genomic Medicine.

#### Small Variant calling

Reads were first trimmed using Trimmomatic with parameters “HEADCROP:10”, “ILLUMINACLIP:NexteraPE-PE.fa:2:30:10”, “LEADING:3”, “TRAILING:3”, “SLIDINGWINDOW:4:15” and “MINLEN:36”. Reads were then mapped with bowtie2 (v. 2.3.4.3) using -sensitive parameters to the Saccharomyces cerevisiae reference genome (v. R64-2-1) with the addition of an extra “chromosome” defining the barcode locus. Reads were sorted, duplicates marked and short variants were called and filtered using GATK (v. 4.0.11.0) AddOrReplaceReadGroups, MarkDuplicates, HaplotypeCaller and VariantFiltration, respectively. Variant filtration used the filter expression “QD<10.0 || FS>20.0 || MQ<20.0 || AN>10 || AF<0.25 || QUAL<100.0 || DP<3” All other commands used default parameters. Variants were annotated with ENSEMBL Variant Effect Predictor using their command-line tool. As many variants had multiple possible annotations, coding sequence annotations (“missense variant”, “frameshift variant”, “stop gained” and “stop lost”) were prioritized over synonymous annotations, which were prioritized over upstream noncoding annotations (within 2kb of a gene) and finally downstream noncoding annotations (again within 2kb of a gene). Variants further of 2kb of any gene or those within 2kb but with no annotations were removed as likely nonfunctional. Finally, variants with less than 3 reads of support for the derived allele were removed as putative false positives.

#### Additional filtering to remove erroneous and ancestral variants

The procedure described above identified 34,720 small variants across 428 sequenced isolates. We expect that many of these variants are sequencing and/or mapping errors that our procedure failed to remove, as well as fixed differences from the reference genome present in the ancestor of our experiment. To further filter out such spurious variants, we estimate the ancestral allele frequency spectrum from 24 sequenced ancestral clones and compare it with a typical allele frequency spectrum of 24 adapted mutants (averaged over 1000 random draws of 24 adapted mutants). As Figure S15 shows, the latter has an excess of variants that are present only in one clone, as expected for de novo mutations. The fact that there is no excess of mutations present in two or more adapted clones suggests that all or most mutations observed in two or more adapted clones are not adaptive. After removing 303 such variants, each of which was detected in 110 isolates on average, we are left with 1842 mutations across 428 sequenced strains (4.30 per clone), of which 33 strains carry no detectable derived small variants (Table S2 and Data S3).

#### Copy number variant calling

To identify large copy number variants (CNVs), we generate coverage plots for each sequenced clone by averaging read depth into 1kb windows with bedtools genomecov. An example plot is shown in Figure S13A. As coverage negatively correlates with the distance to telomeres (Figure S14), we re-calculate coverage after correcting for this variation (Figure S13B). We then manually identify CNVs from these corrected coverage plots by visualizing the coverage distribution at higher resolution (Figure S13C).

We identified 176 CNVs across 167 strains (Data S3), of which 85 are whole-chromosome aneuploidies. We found no CNVs in 32 sequenced ancestral and neutral isolates (Figure S16C, Table S2), we estimate the frequency of observing a non-adaptive CNV event as at most 1/32. Thus, we expect at most 12.4 such events among 396 sequenced adaptive clones. In fact, we found 176 CNV events, which suggests that all or almost all of them are adaptive (expected FDR ≤ 0.07; Data S4).

#### Identification of driver loci carrying small adaptive mutations

To differentiate adaptive “driver” mutations among residual ancestral and erroneous variants as well as nonadaptive “passenger” mutations, we rely on the idea of genetic parallelism, i.e., the fact that loci under selection gain mutations in independent lineages more often than expected by chance^87,88^. For each gene, we define its multiplicity as the number of clones that carry a mutation in this gene. Since shorter genes require lower multiplicity to be called adaptive, we bin genes by their length into six 1kb bins plus one bin for genes with length ≥ 6kb. For each length bin *l* = 1,2, …,7, we count the number of genes whose multiplicity is *m* = 1,2, …, denoting these counts by *k_lm_*. We obtain the number of such genes 〈*k_lm_*〉 expected in the absence of selection as follows.

We randomly and independently redistribute *N* = 1718 mutations (small mutations in adaptive clones after removing multiple mutations in the same gene in the same clone) across 7226 yeast genes 1000 times, with the probability for each mutation landing in a given gene being proportional to its length + 2kb. Then, the observed excess number of mutations with multiplicity m in length bin *l* is *a_lm_* = max{0, *k_lm_* – 〈*k_lm_*〉}. These excess mutations are assumed to be adaptive. We redistribute the remaining 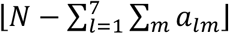 potentially non-adaptive mutations as before, and identify additional excess mutations as adaptive. We repeat this procedure iteratively until the total number of excess mutations is ≤ 1. We achieve convergence after 8 iterations and thereby obtain the final expected counts 〈*k_lm_*〉.

Next, we would like to identify specific loci that carry adaptive mutations as those with multiplicities above some threshold. Specifically, we would like to determine multiplicity thresholds *M_l_* for each length bin *l* = 1, …,7, so that all loci in that bin with multiplicities ≥ *M_l_* are called adaptive. To do so, for all *l* we calculate the expected FDR at the given multiplicity threshold *M_l_* as

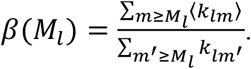

We choose the multiplicity thresholds *M_l_*, so that β(*M_l_*) ≤ β* for all *l* and for some desired β*. We use β* = 10%. We estimate the overall FDR as

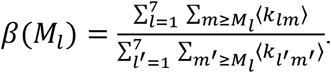

We conduct this analysis considering all 396 sequenced adaptive isolates together, as well as A- and C-mutants separately. Loci identified in any one of these three analyses are defined as putative adaptive loci. After identifying putative 63 adaptive loci this way, we assume that all discovered 185 mutations at these loci are adaptive (Data S4). An extended discussion of our analysis of adaptive mutations can be found in the Supplementary Information (Section 3).

#### Probability of genetic parallelism

We calculate the probability of genetic parallelism *P*_g_ for a set of mutants as follows. We consider every pair of mutants, and calculate the proportion that have a mutation in at least one common adaptive locus (including both small variants and CNVs). Adaptive isolates with no mutations at the identified adaptive loci are assumed to have an adaptive mutation at a locus that is not shared with any other isolate in the set. To test whether A- and C-mutants have different probabilities of genetic parallelism, we reshuffle the evolution treatment label (i.e. Alone vs Community) across all A- and C-mutants and calculate *P*_g_ for both resulting groups. We obtain the absolute value of the difference in *P*_g_ between the two groups, |Δ*P*_g_|, in 1000 such permutations, and estimate the *P*-value as the fraction of permutations where |Δ*P*_g_| exceeds the observed difference.

### Phenotyping

For all phenotypic measurements, replicate measurements were conducted using distinct samples. In no case was a single sample measured repeatedly.

#### Measurement of ancestral yeast and algae growth

Yeast and algae were thawed from frozen stocks (−80°C for yeast, LN2 storage for algae) and grown separately in standard growth conditions for one growth cycle. Communities were inoculated by mixing 100 μL of yeast with 10^4^ cells of algae into 10 mL of our standard growth culture and propagated for 1 growth cycle via 100-fold dilution. On the second growth cycle, each culture (yeast alone, algae alone and the community, with six replicates each) was characterized daily by CFU counting for the yeast, and chlorophyll fluorescence measurements for the alga. Each culture was plated onto CYM media plates (1% agarose for) and were incubated at 30°C. Colonies were counted to estimate the density of yeast in each culture at each timepoint. We estimate alga density by measuring chlorophyll b fluorescence. To that end, we transfer 200μL of each culture into a well of a black-wall clear-bottom 96-well plate (Corning) and measure fluorescence in a plate reader (Molecular Devices Spectramax i3x, excitation at 435nm and observation at 670nm). Chlorophyll fluorescence intensity measurements were converted into cell density estimates by using a calibration curve as described in the Supplementary Information (Section 5.1).

#### Selection of A- and C-mutants for phenotyping experiments

We selected 31 C-mutants and 28 A-mutants to cover a diversity of mutations and fitness values represented among all sampled adaptive mutants (see Data S4 for the number of mutants selected from each mutation class). Our reasoning for this non-random sampling was that mutants carrying a driver mutation at the same locus would have similar phenotypic values. If we selected clones for phenotyping randomly, we would have likely not observed more rare phenotypes. To account for this over-dispersion in the selection of mutants we apply the mutation weighting procedure described below.

#### Measurement of YYA

To estimate YYA, we inoculated all 60 yeast strains (including the ancestor) individually into the standard conditions (10 mL of media in 50 mL flasks) from frozen stocks and propagated them for two cycles (10 days). During the third cycle, we estimated yeast densities after 5 days of growth by plating and colony counting as described below. Correlations between replicate measurements are shown in Figure S19A.

#### Measurements of YYC and AYC

We create mutant communities as follows. On day 0, we inoculate 50 mL growth media (in 250 mL non-baffled Erlenmeyer flask) with 1 mL of CC1690 *C. reinhardtii* stock stored in a liquid nitrogen freezer and incubate for 20 days. On day 20, 100 μL of this culture are transferred to the standard conditions (10 mL of fresh media in a 50 mL flask). Also on day 20, the 60 yeast strains (including the ancestor) are individually inoculated into the standard condition from frozen stocks. On day 25, we form the mutant communities by transferring 100 μL of each yeast culture and 200 μL of the algae culture into fresh media (200 μL of alga were used instead of 100 μL because the density of algae culture was approximately 50% of that at the initiation of the BLT experiment). These mutant communities are grown for one cycle in our standard conditions. On day 30, we transfer 100 μL of each mutant community into 10 mL fresh media, as in the BLT experiment. We estimate both yeast and alga density on day 35, as described previously. Yield estimates can be found in Data S2. Correlations between replicate measurements are shown in Figure S19B,C.

#### Microscopy

To detect potential physical associations between algae and beneficial yeast mutants, we created mutant communities as described above. After 5 days of growth, communities were then mounted on glass microscope slides (Fisher Scientific 12550143), sealed with Dow Corning high vacuum grease (Amazon B001UHMNW0) and imaged on a light microscope using a 20x objective with DIC. Extended Data Figure 8 shows one representative community; the remaining 17 imaged mutant communities along with WT controls can be found in the Dryad data repository.

#### Mutant growth curve measurements

To estimate the growth parameters *r* and *K* for individual beneficial mutants, we carried out growth curve measurement experiments of individual yeast mutants and the ancestor in the A-condition. To this end, we inoculated all 60 yeast strains (including the ancestor) individually into the BLT condition (10mL of media in 50mL flasks) from frozen stocks and propagated them for two cycles (10 days). During the third cycle, we estimated yeast densities on days 10, 10.5, 11, 11.5, 12, 13, 14 and 15 by plating and colony counting as described in the section “Measurements of YYC and AYC.” The growth curves are shown in Figure S20 and the data are provided in Data S2.

We estimate *r* as the slope of the relationship between log(CFU/mL) and time (in hours) for the three measurements between 12 and 36 hours of growth. We estimate *K* as the maximum observed density (in CFU/mL). *r* and *K* estimates can be found in Data S2. Correlations between replicate measurements are shown in Figure S21.

#### Mutation weighting

Even though not all A- or C-mutants were phenotyped, we would like to make certain statistical statements about the distribution of phenotypes among all sampled A- or C-mutants. To this end, we associate each of the 59 phenotyped mutants with a single driver mutation. Mutants with multiple driver mutations are associated only with the most common driver mutation. Mutants with no identified driver mutations are associated with a unique unknown mutation. To obtain the prevalence of a given phenotypic value among all A- or C-mutants, we weight each measured phenotypic value by the number of sequenced A- or C-mutants with the same driver locus as the phenotyped mutant and divide by the total number of phenotyped A- or C-mutants.

#### Kernel density estimation

We use kernel density estimate (KDE) to determine how likely certain phenotypic trait values would occur among all A- and/or C-mutants. Specifically, we obtain the kernel density estimates *D_A_*(*y,a*) and *D_c_*(*y,a*) for the probabilities that a community formed by the ancestral alga and a random A- or C-mutant, respectively, would produce yeast yield *y* and alga yield *a*. To estimate *D_A_*, we apply the kde2d function in R with bandwidth 1 along the *x*-axis and 4/3 along the *y*-axis to the mutation-weighted yield data for the A-mutants. We analogously obtain *D_C_*(*y,a*).

#### Accounting for measurement errors in statistical tests by permutation testing

As the measurement errors in our estimates of phenotypic values (*r*, *K*, yeast yield and algae yield) are quite large, we use a permutation and resampling procedure to determine the statistical significance in various tests involving these variables. In this procedure, we permute isolate labels and resample the phenotypic values associated with each isolate from the normal distribution with the estimated mean and the estimated standard error of the mean. We carry out 1000 permutation and resampling instances in each test. To estimate the false positive rate, we count the fraction of resampled values that are significant at a chosen threshold and averaged this number over all permutations. False discovery rate (FDR) is then computed by dividing the average number of false positives by the number of observed positives.

Permutation tests of Pearson correlation significance are conducted as follows. For each permutation, isolate labels for each variable are permuted independently and phenotypic values are resampled as above. A randomized correlation coefficient is determined from each of the 1000 permutations, and significance is determined by the proportion of randomized *R*^2^ values that exceed the observed *R*^2^ value.

#### Phenotypic parallelism analysis

We quantify the degree of parallelism among a set of mutants with respect to a pair of quantitative traits *X* and *Y* by estimating the probability that, for two randomly selected mutants, trait *X* changes in the same direction in both mutants and trait *Y* changes in the same direction in both mutants. Mathematically, if one randomly selected mutant has trait increments Δ*X_i_* and Δ*Y_i_* relative to the ancestor and the other mutant has trait increments Δ*X_j_* and Δ*Y_j_*, we estimate the probability that both (Δ*X_i_)(ΔX_j_*) ≥ 0 and (Δ*Y_i_*)(Δ*Y_j_*) ≥ 0. This measure of phenotypic parallelism emphasizes the direction of change rather than the magnitude. When we compute this measure for the pair of yields of mutant communities, in which case *X* and *Y* are yeast and alga yields, we refer to it as the probability of ecological parallelism, *P*_e_. When we compute this measure for the growth phenotypes, in which case *X* and *Y* are *r* and *K*, we refer to it as the probability of phenotypic parallelism, *P*_ph_.

We obtain these probabilities of parallelism for the A- and C-mutants. We test the significance of the deviation of these probabilities from the expectation of 25% parallelism via a χ^2^-test. To determine the statistical significance of the difference between the probabilities of parallelism for the A- and C-mutants, we resample the phenotypic values of each of the 59 mutants from the normal distribution (see above) and permute mutant genotype and home-environment labels, so that the genotype and label are always associated with each other but dissociated from the phenotypic values. We then calculate the parallelism probabilities for these permuted and resampled data. We estimate the *P*-value by carrying out this permutation and resampling procedure 1000 times.

## Data Availability

All raw sequencing data is available on the US National Center for Biotechnology Information (NCBI) Sequence Read Archive (SRA) under BioProject PRJNA735257. Other input data (e.g. growth data, variant calls, community yield etc) can be found on Dryad at https://doi.org/10.6076/D14K5X.

## Code Availability

The latest version of the barcode counting software BarcodeCounter2 can be found at https://github.com/sandeepvenkataram/BarcodeCounter2.git. Analysis scripts can be found on Dryad at https://doi.org/10.6076/D14K5X.

## Acknowledgements

We thank Gavin Sherlock and Katja Schwartz for providing the barcoded yeast library, Stephen Mayfield and Frank Fields for laboratory equipment and help with algal husbandry, Rachel Dutton and Manon Morin for help with sequencing, Scott Rifkin and Jessica Bloom for help with microscopy, STARS students Jesse Yu and Sophia Rosemann for help with experiments, Justin Meyer, Alena Martsul, Shohreh Sikaroodi for technical assistance, the Kryazhimskiy, Meyer and Hwa labs, Damien Barrett, Josh Borin, Shermin de Silva, Susanne Dunker, Nandita Garud, Stan Harpole, Canan Karakoç, Holly Moeller, Dmitri Petrov, and Peter Zee for feedback on the manuscript. Sequencing was done in part at the UCSD IGM center (University of California, San Diego, La Jolla, CA). We acknowledge the San Diego Supercomputing Center for the use of the TSCC cluster for computing services. EFYH is funded by National Science Foundation CAREER grant 1846376 and Deutsches Zentrum für Integrative Biodiversitätsforschung (iDiv) grant DFG–FZT 118, 202548816. SK is funded by BWF Career Award at the Scientific Interface grant 1010719.01, Alfred P. Sloan Foundation grant FG-2017-9227 and the Hellman Foundation.

## Author contributions

conceptualization (SV, EFYH, SK), methodology (SV, HYK, SK), data acquisition (SV), analysis (SV, HYK, SK), initial manuscript (SV, SK), editing (SV, HYK, EFYH, SK), supervision (EFYH, SK), funding (SK).

## Materials and Correspondence

All raw sequencing data is available on the US National Center for Biotechnology Information (NCBI) Sequence Read Archive (SRA) under BioProject PRJNA735257. Other input data (e.g. growth data, variant calls, community yield etc) and analysis scripts can be found in Data Files S1–S4 and on Dryad at https://doi.org/10.6076/D14K5X. Strains and other biological materials are available by request to SK.

## Code availability

The latest version of the barcode counting software BarcodeCounter2 can be found at https://github.com/sandeepvenkataram/BarcodeCounter2.git. All other analysis scripts used for this study are available on Dryad at https://doi.org/10.6076/D14K5X.

## Code and data availability for reviewers

As the data and code is not currently publicly available on Dryad, reviewers can access this data through the following reviewer link: https://datadryad.org/stash/share/g4RSTbYCCwCpQxidH0dhQDKf7SsVsijY8pjzcsy7Y88

## Extended Data Figures

**Extended Data Figure 1:**
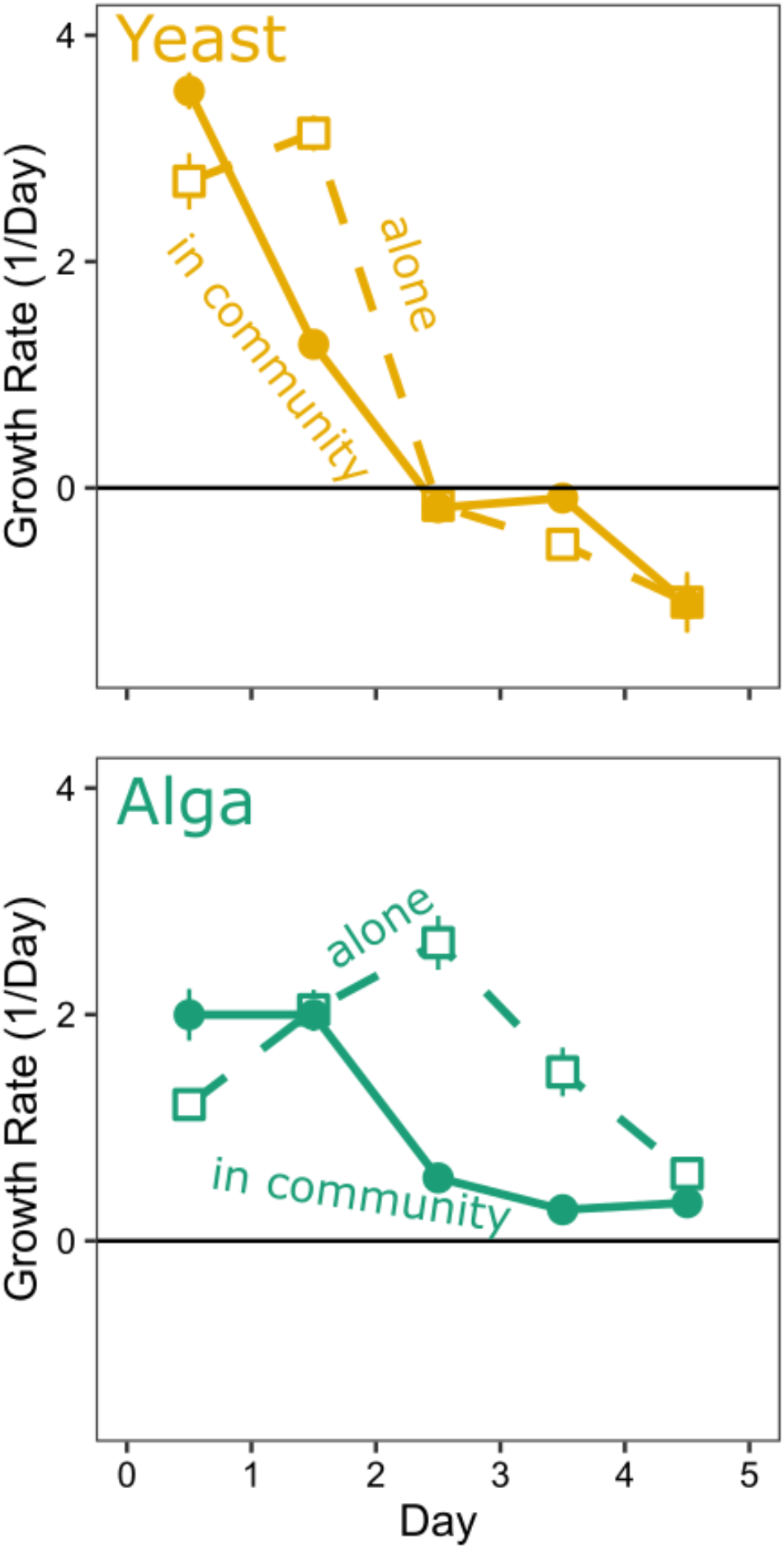
Per capita net population change for the ancestral yeast and alga. Same data as in Figure 1. Error bars show ±1 standard error of the mean.

**Extended Data Figure 2:**
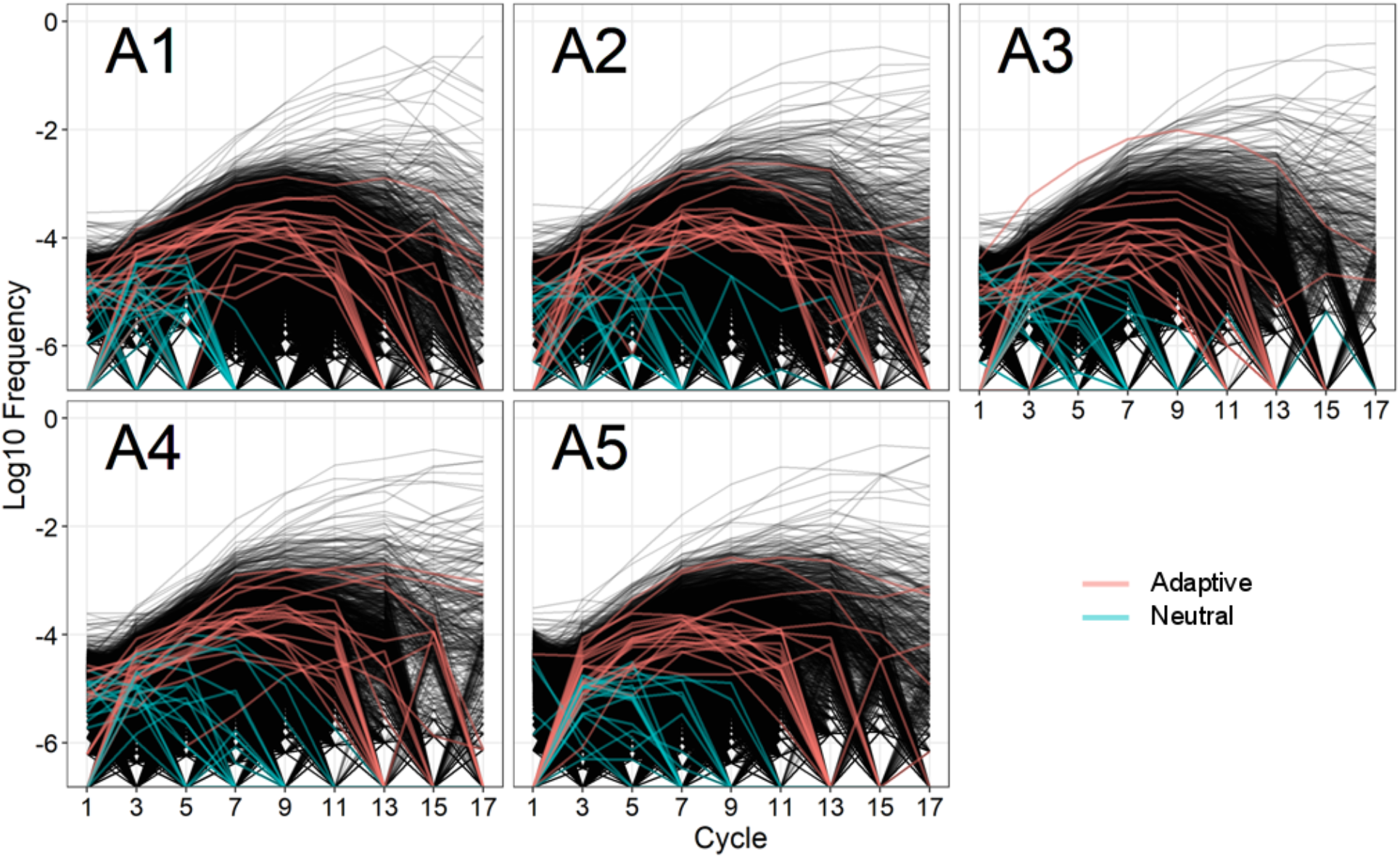
Frequency trajectories of barcoded lineages in yeast in the A-condition. Each panel corresponds to a BLT replicate population in the A-condition, as indicated. Lineage frequencies were measured at every odd cycle. Twenty random adapted lineages are shown in red, and twenty random neutral lineages are shown in blue.

**Extended Data Figure 3:**
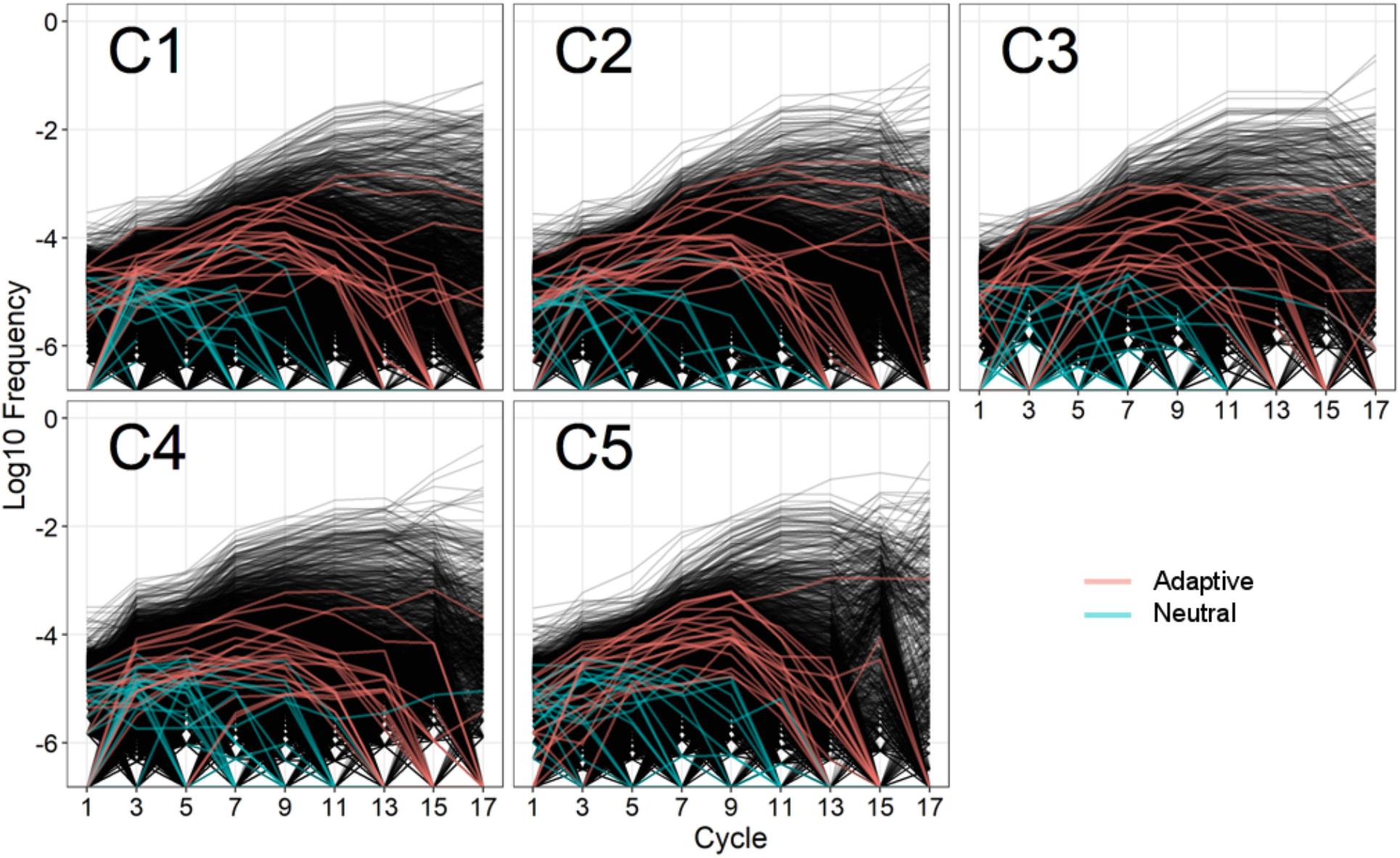
Frequency trajectories of barcoded lineages in yeast in the C-condition. Each panel corresponds to a BLT replicate population in the C-condition, as indicated. Notations are as in the Extended Data Figure 2.

**Extended Data Figure 4:**
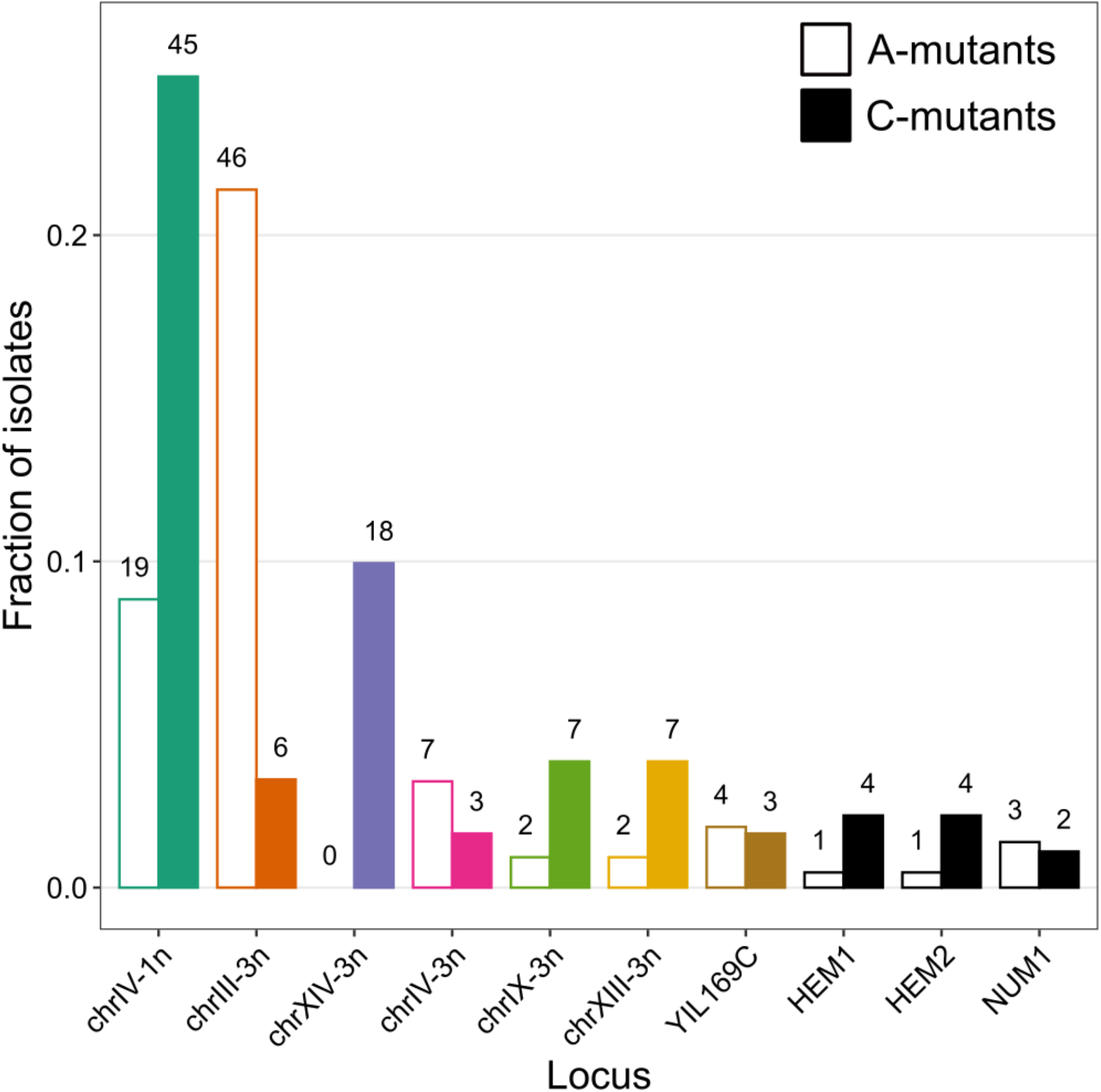
Distribution of adaptive mutations across the most common driver loci. Only driver loci with 5 or more mutations are shown (see Data S3 for the full distribution). Colors correspond to Figure 2 in the main text.

**Extended Data Figure 5:**
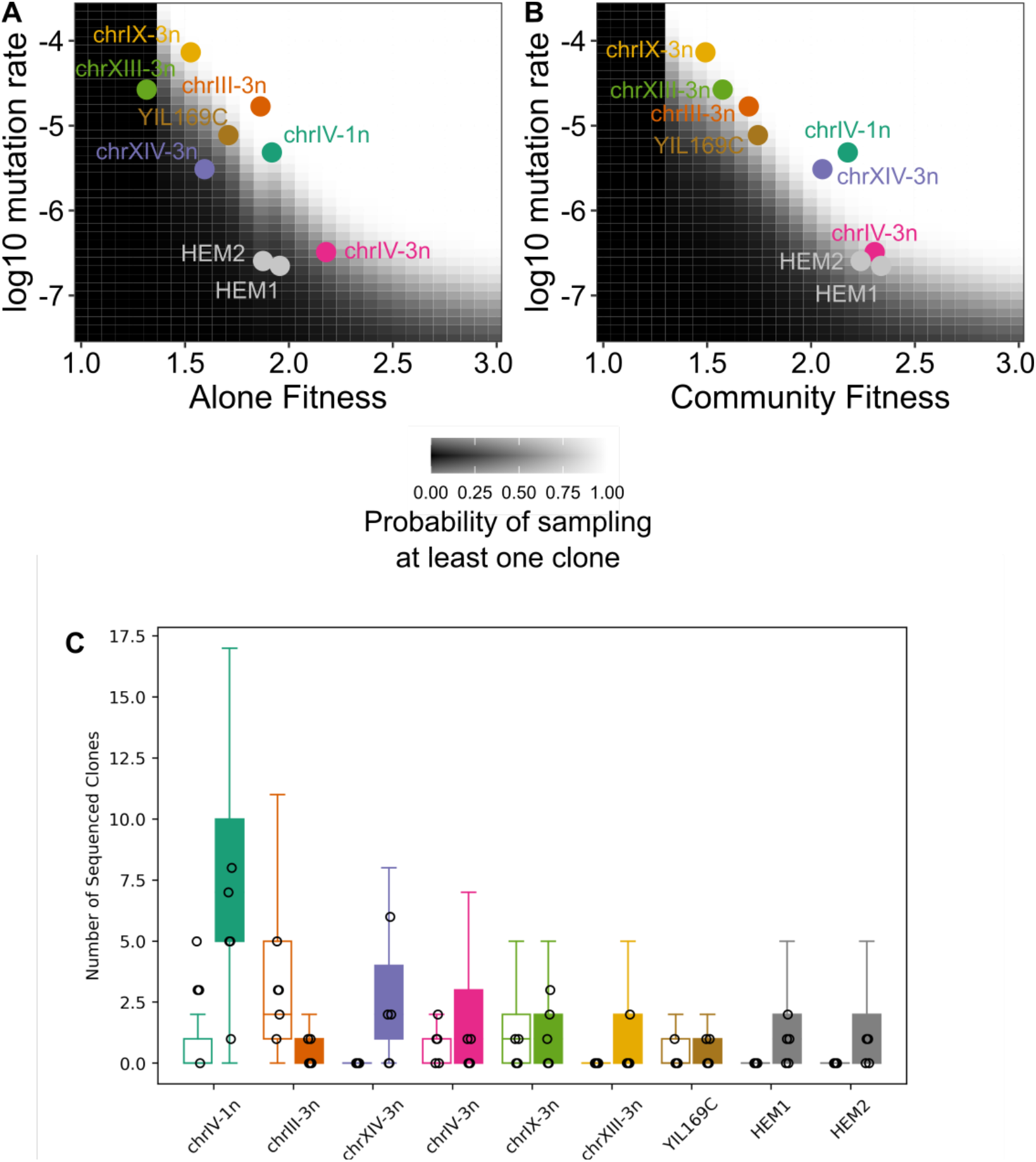
Probabilities of observing adaptive mutations at the most common driver loci in the whole-genome sequencing data. **A.** Shades of gray show the probability of sampling at least one clone with a beneficial mutation that arises at a certain rate (*x*-axis) and provide a certain fitness benefit in the A-condition (*y*-axis). The most common driver loci are shown by points (colors are the same as in Figure 2 in the main text). The estimated beneficial mutation rate and the selection coefficient for each mutation class are given in Table S3. **B.** Same as A but for the C-condition. The mutation rate for each locus is assumed to be the same in both conditions, but the selection coefficients vary. **C.** The observed number of sequenced clones per replicate population with a mutation at each locus (black points) and the numbers expected in our simulations (bars and whiskers). Bars show IQR (Q3 – Q1), whiskers show Q1 – 1.5 × IQR and Q3 + 1.5 × IQR.

**Extended Data Figure 6:**
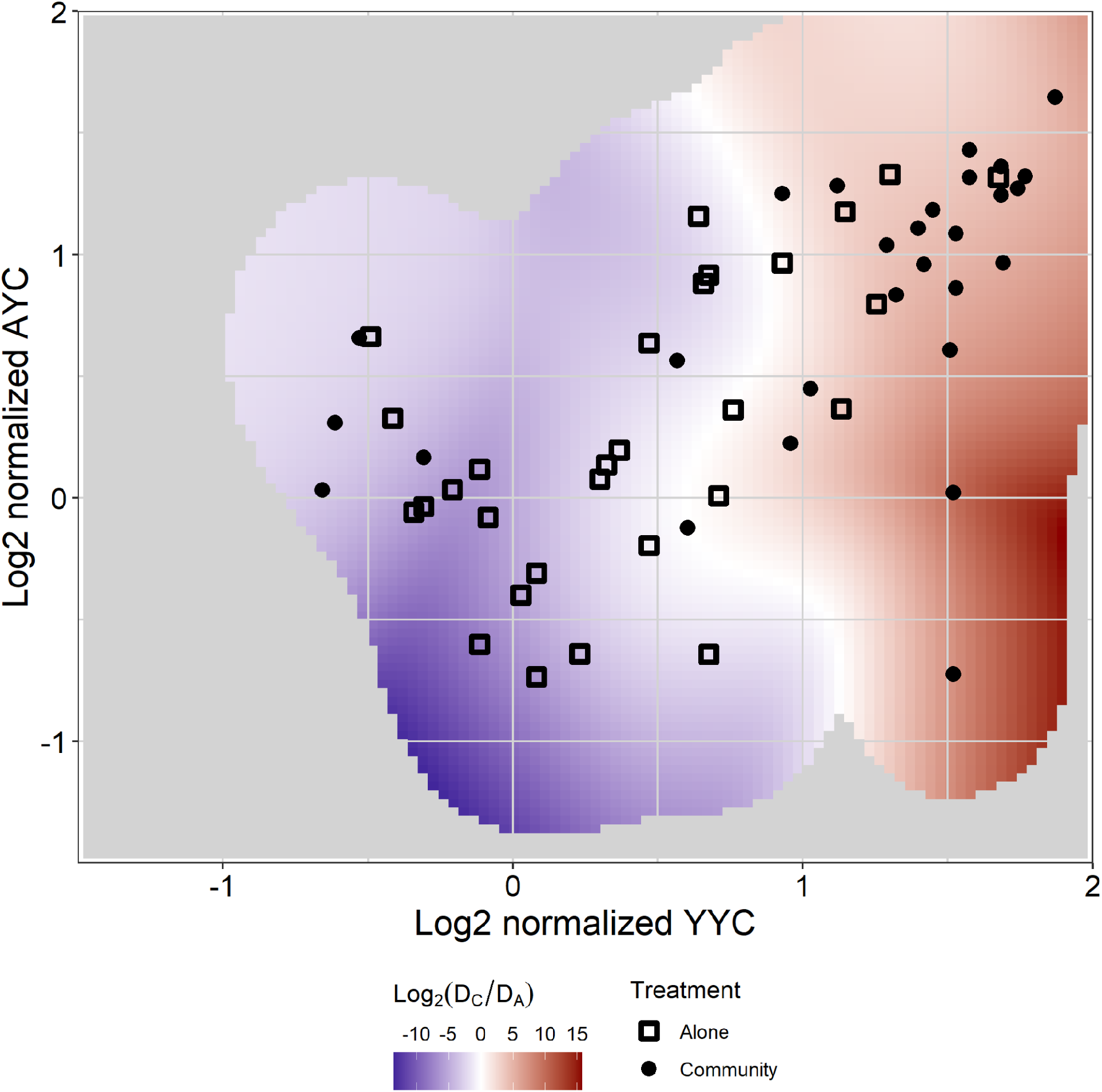
The weighted distribution of yields. The heatmap shows the ratio of probabilities D_C_ and D_A_ of observing a given pair of yields among C- and A-mutants. Data points are identical to Figure 3B in the main text. To estimate D_A_ and D_C_, each data point is weighted by the frequency of occurrence of the corresponding mutation among the A- and C-mutants, respectively (see Methods for details). Regions where either D_A_ or D_C_ falls below 0.03 are colored gray. YYC and YYA are normalized by the respective ancestral values.

**Extended Data Figure 7:**
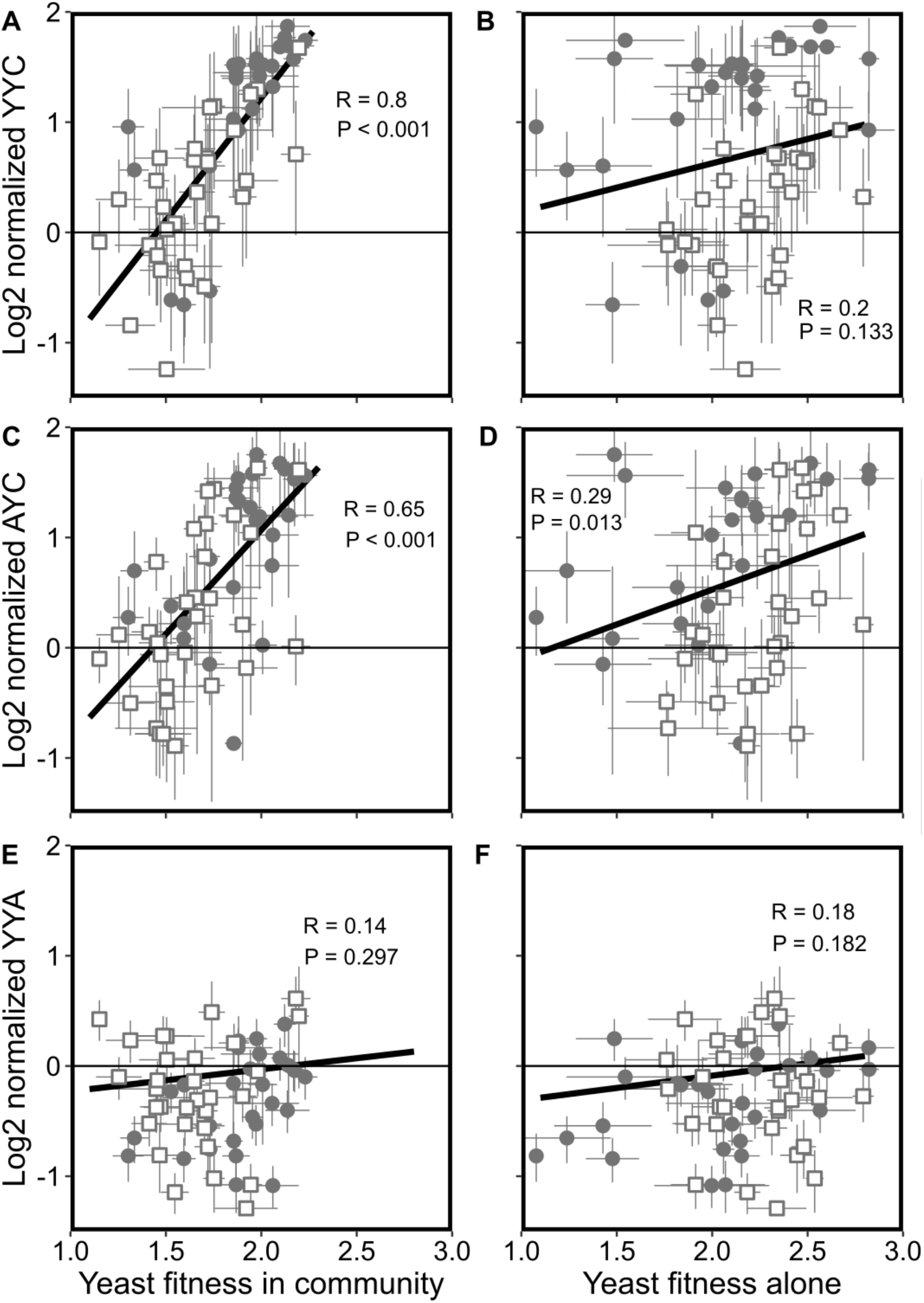
Correlations between competitive fitness and yields. Normalization is relative to the ancestor. Error bars represent ±1 standard error of the mean.

**Extended Data Figure 8:**
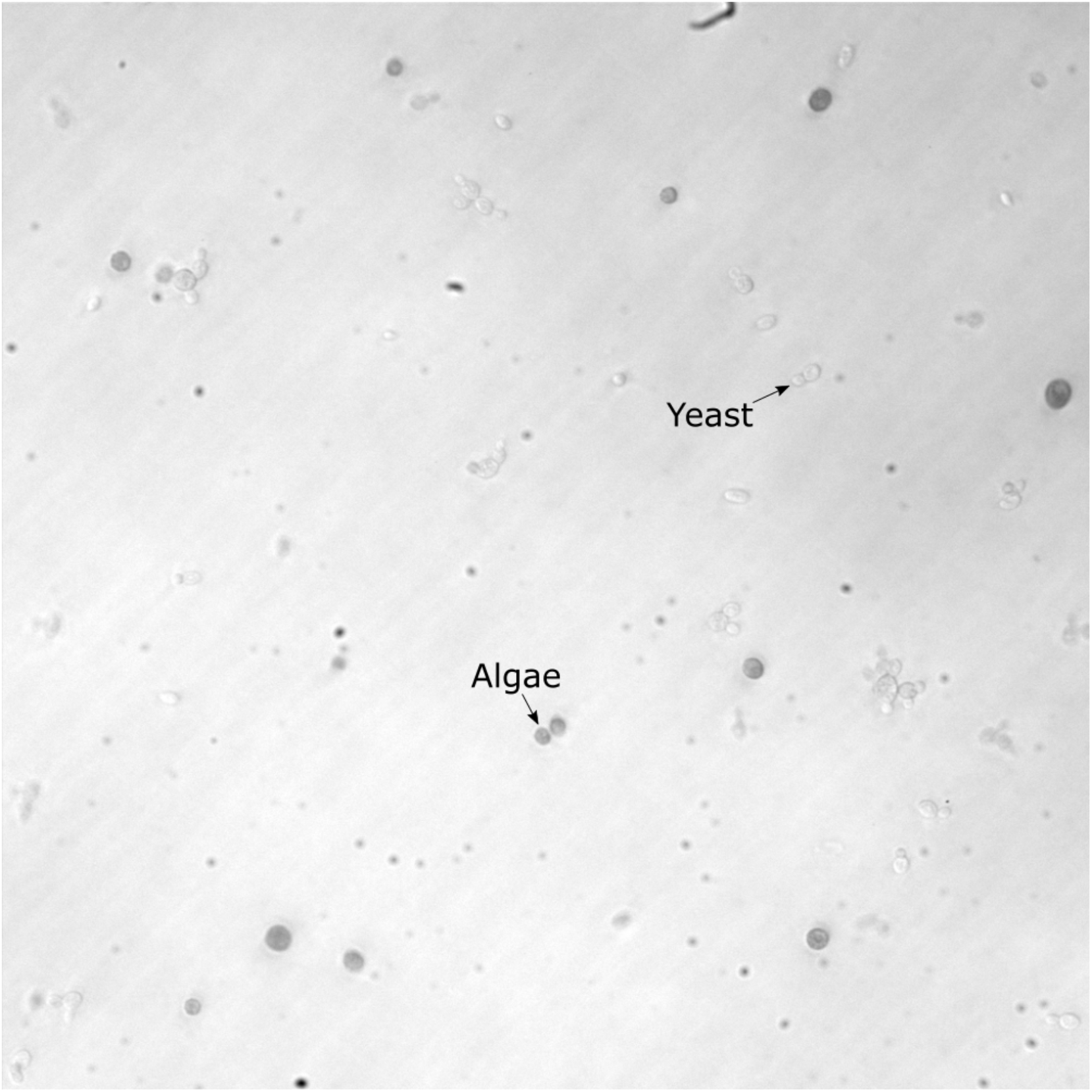
Representative microscopy image showing lack of physical associations between yeast and algae cells. Mutant culture formed by the C-mutant C2 (barcode ID 109098) is shown. Yeast and alga cells are indicated with arrows.

**Extended Data Figure 9:**
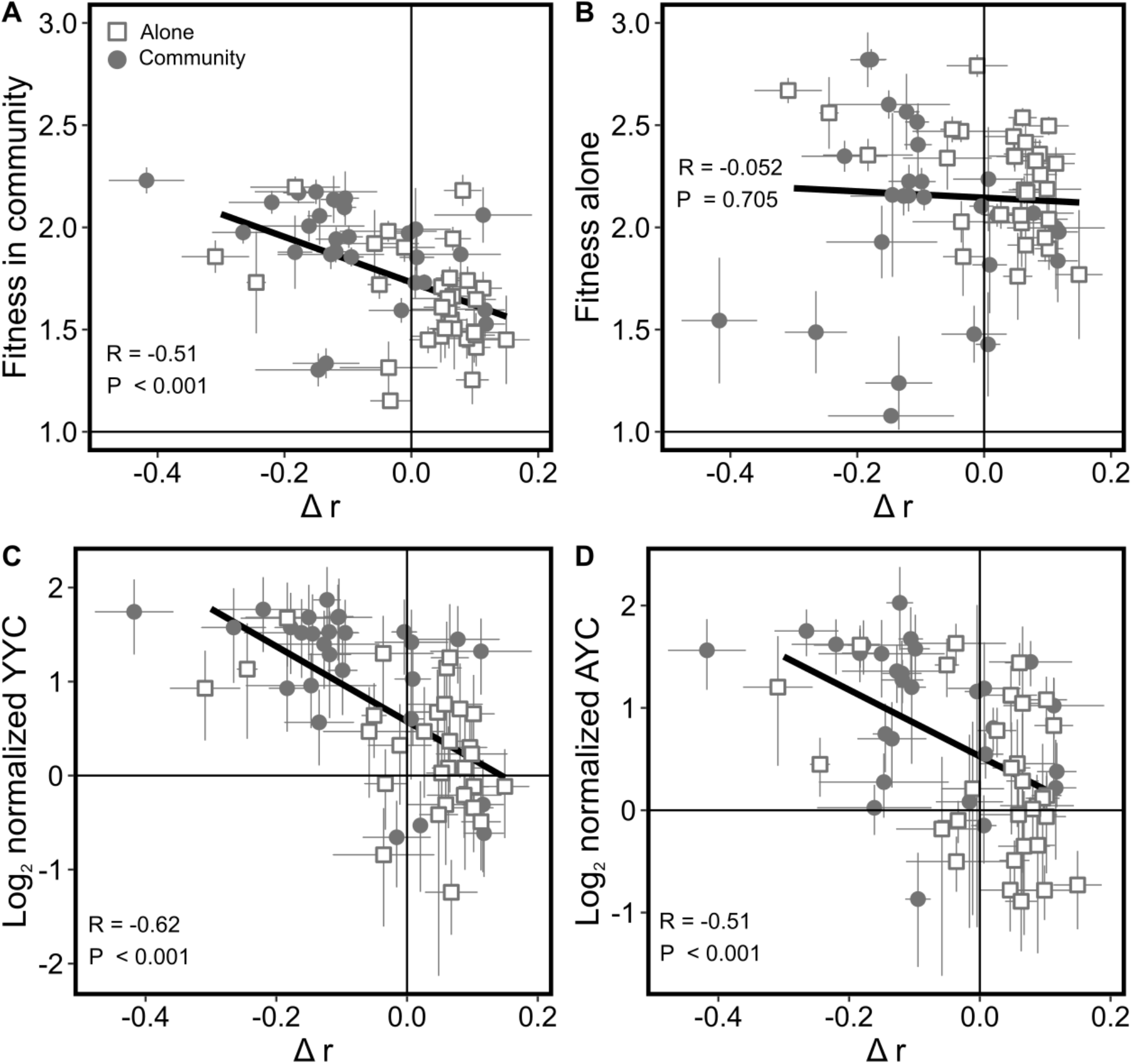
Relationship between growth rate, yields and fitness. In all panels, normalization is relative to the ancestor. Error bars represent ±1 standard error of the mean.

**Extended Data Figure 10:**
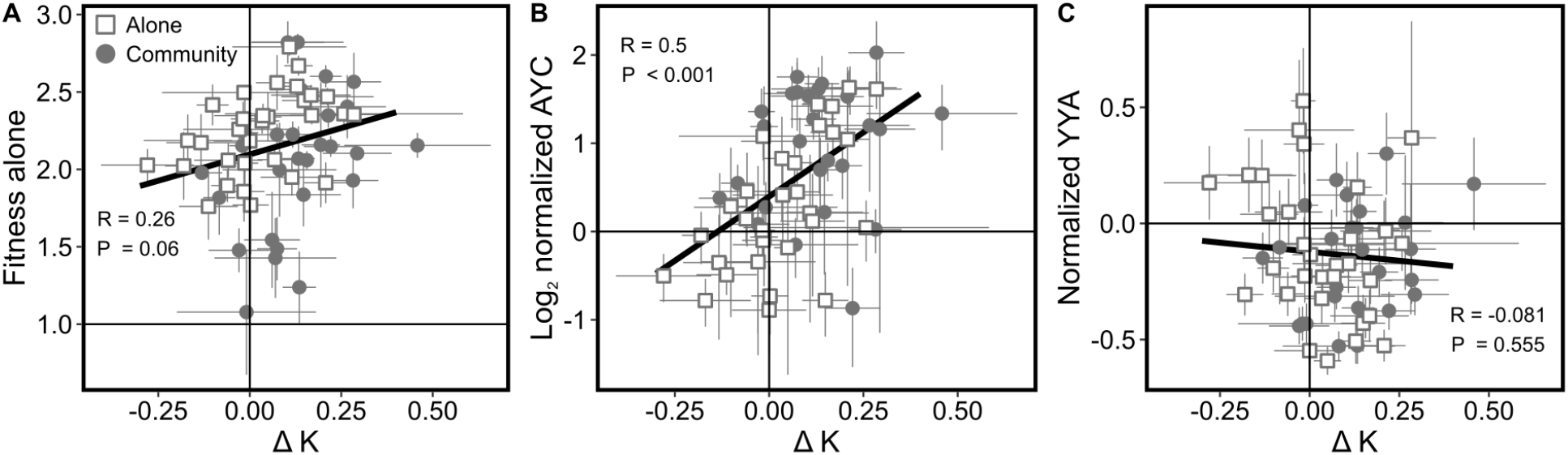
Relationship between carrying capacity, yields and fitness. In all panels, normalization is relative to the ancestor. Error bars represent ±1 standard error of the mean.

## Supplementary Materials

Supplementary Information Text

**Data S1.** Absolute species abundances over a five-day growth period.

**Data S2.** List of all sampled isolates, their fitness, phenotypes and yields.

**Data S3.** List of all identified mutations.

**Data S4.** Distribution of adaptive mutations across driver loci.

## Supplementary Information

## 1 Barcode lineage tracking (BLT) experiment and data analysis

### 1.1 Number of yeast and alga generations per cycle

#### Yeast

The rate at which new mutations arise in a yeast population and the amount of physical time it takes for these mutations to survive drift and establish depends on the number generations that yeast go through during our BLT experiment. Thus, if yeast go through substantially different numbers of generations per cycle between the A- and the C-conditions, it could cause differences in our power to detect adapted lineages, which in turn could confound our ability to compare measured bDFEs. In fact, it is possible that yeast go through different number of generations because they reach different peak densities and because they likely do not reproduce after Day 2 in the A-condition, but they likely do reproduce in the C-condition (since the alga supplies the limiting ammonium).

We can estimate the number of generations in the A-condition if we assume that there is no death during Days 1–2 and there is no growth during Days 2–5. Given that the yeast starts the growth cycle at 1.37 × 10^4^ cells per mL and reaches peak density of 4.72 × 10^6^ cells per mL on Day 2 (Figure 1, Data S1), the estimated number of generations per cycle in the A-condition is 8.4. Knowing that yeast yield in the A-condition is 9.83 × 10^5^ cells per mL, we can also estimate the average per capita death rate in the A-condition in Days 3–5 as 0.52 per day.

We can estimate the number of generations per cycle in the C-condition as follows. We again assume that there is no death in Days 1–2. Thus, given that the yeast starts the growth cycle at 2.39 × 10^4^ cells per mL and reaches peak density of 2.83 × 10^6^ cells per mL on Day 2 (Figure 1, Data S1), we estimate the number of generations in this part of the cycle to be 6.9. To estimate the number of generations in Days 2–5, we assume that the per capita death rate during this phase of the cycle is the same in the C-condition as in the A-condition, i.e., 0.52 per day. Thus, in the absence of new births, yeast would have reached density of 5.95 × 10^5^ cells per mL by the end of the cycle. Instead, we observe that yeast yield is 8.78 × 10^5^ cells per mL, which implies that 2.83 × 10^5^ new yeast cells were produced during this period, corresponding to 0.13 generations. Thus, we estimate that yeast go through on average 7.0 generations per cycle in the C-condition.

Given that yeast go through fewer generations and generally have lower population sizes in the C-condition than in the A-condition, we would expect that fewer adaptive mutations would arise in the C-condition and it would take them longer to spread. However, we actually detect slightly more adapted lineages in the C-condition than in the A-condition, suggesting that these differences do not substantially undermine our ability to identify adapted lineages.

#### Alga

Since the alga grow continuously during the entire cycle both alone and in community with yeast, we can estimate their number of generations by assuming that there is no death. Given that the alga growing alone starts at density 9.3 × 10^3^ cells per mL and reaches density 2.6 × 10^6^ cells per mL(Figure 1), we estimate that the alga goes through approximately 8.1 generations when growing alone. Given that the alga growing in community with yeast starts at density 4.83 × 10^4^ cells per mL and reaches density 7.90 × 10^6^ cells per mL, we estimate that the alga goes through approximately 7.4 generations when growing in community.

### 1.2 Justification for ignoring adaptation in alga

In the C-condition, that is, when yeast and alga are co-cultured together, yeast and alga reach final yields of 8.78 × 10^6^ and 7.90 × 10^7^ cells, respectively, with the bottleneck sizes being by a factor of 100 smaller. Numerous evolution experiments with yeast populations of comparable or smaller size across various environmental conditions have shown that adaptive mutations arise and reach high frequencies within 250 generations or less [1, 2, 3, 4, 5]. Although much less is known about the rates of adaptation in *Chlamydomonas reinhardtii*, one study reports a failure of the alga (strain CC2344) to adapt within 1000 generations of evolution at the bottleneck size of 10^5^ cells [6]. Another study reports 35% growth rate gain after 1880 generations of evolution in alga strain CC-503 cw92 mt+ [7] with a comparable population size to ours and detectable gains appeared only after about 300 generations. Our community BLT experiments last for about 66 algal generations until adapted yeast mutants are sampled, which is likely insufficient for new mutations to arise in the alga population. Furthermore, since the environment in our experiment is well-mixed and there is no evidence for physical associations between the two species (see Extended Data Figure 8), the only way the alga can modify the environment for yeast is through the medium. Thus, even if some alga mutants arose during the BLT experiment, they would not be able to reach high enough frequencies in the population to substantially alter the medium.

### 1.3 BLT data analysis

#### Sanity checks

We performed two “sanity” checks to ensure that our analysis gives reasonable results. First, mean fitness is expected to monotonically increase over time in each population. We find that mean fitness does indeed increase monotonically during the first 11 cycles in all but one populations (Figure S1B). Second, we visually inspected how well individual lineage trajectories are fitted by equation

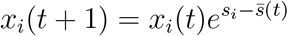

and found that these fits are generally very good (Figure S1C,D shows typical fits).

#### Sensitivity of the procedure for detecting adapted lineages with respect to the choices of parameters

The heuristic procedure for identifying adapted lineages has three key parameters: (1) the minimum lineage frequency threshold (set to 10^−4^); (2) the expansion factor used to identify neutral lineages (set to 100); and (3) the “SE parameter” which is the number of standard errors separating the lineage’s fitness from zero required to call it adapted (set to 2). We tested the sensitivity of our procedure with respect to the choice of these parameters.

We varied the minimum lineage frequency threshold and the expansion factor by an order of magnitude in either direction. We find that the number of adapted lineages that we detect varies primarily with the minimum lineage frequency threshold, as expected (Figure S2C). However, our claims regarding the differences between the bDFEs in the A- and C-condition remain reasonably robust (Figure S2A,B). The most sensitive result is the difference in the width of the bDFEs which disappears when we lower the minimum lineage frequency threshold, presumably because too many non-adapted lineages are being falsely called adapted.

We then tested how the SE parameter affects the bDFE median in each condition, while keeping the other two parameters at their chosen values. We find that the bDFE median as a function of the SE parameter has a plateau between values 1 and 2 (Figure S3). When the SE parameter declines below 1, we see a precipitous decline in the bDFE median, as would be expected from the associated growth in the number of false positives. When the SE parameter is increased beyond 2, we observe a gradual increase in the bDFE mean, as would be expected from the associated growth in false negatives. This result suggests that any choice of the SE parameter within the plateau would be appropriate, as it would strike a balance between keeping down both false positives and false negatives.

#### Precision and accuracy of the procedure for detecting adapted lineages

We developed a simulation framework to quantify the precision and accuracy of our procedure. We simulated evolution using a population initialized with 100,000 barcoded lineages, of which 2,000 had already acquired an adaptive mutation prior to the beginning of the evolutionary simulation. The fitness effects of these mutations per growth cycle were drawn from a normal distribution, with *μ* = 0.4 and *σ* = 0.2. These parameters were selected to allow for relatively weak adaptation, as a strong adaptation regime would lead to extinction of many adapted mutants. Observing adaptation in this regime allows us to get better statistics on the precision and accuracy of our method.

We simulated the evolution of our population iteratively as follows. Given that the population size of lineage *i* at the beginning of cycle *t* is 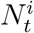, by the end of the cycle it deterministically grows or shrinks to

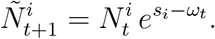

Here, *s_i_* is the fitness of lineage *i*, 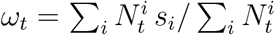 is the mean fitness of the population at time *t*. After growth, we simulate dilution, so that the size of lineage *i* at the beginning of cycle *t* + 1 is drawn from the Poisson distribution with parameter 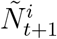. The initial size *N*_0_ of each lineage was drawn from an exponential distribution with mean 100.

The population was propagated for total 18 growth cycles. Every other cycle, barcodes were sampled and “sequenced”. We simulate this stochastic process by drawing 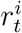 reads of barcode *i* from the negative binomial distribution with mean 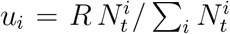, and variance *u_i_* (1 + *ϵu_i_*). The total read depth was set as *R* = 10^6^. The error parameter *ϵ* = 0.01 allows for variation greater than the mean and represents systematic sources of experimental error (gDNA extraction, PRCs, sequencing). The choice of the negative binomial distribution and parameter *ϵ* was based on Ref. [8]. We then applied our lineage calling procedure to these simulated barcode data.

Our procedure identified 2306 lineages as adapted, of which 1413 were true positives and 893 were false positives (Figure S4). We thus estimate the false positive rate to be 71%, the false positive rate to be 1% and the false discovery rate to be 39%. For the true positives, the estimated fitness correlates very well with their true fitness (Figure S5A). The median of the estimated bDFE is lower compared to that of the underlying true bDFE (0.314 versus 0.406, Figure S5B), presumably due the presence of false positives. Indeed, 93% (829 out of 893) of the false positives have estimated fitness effects below the median. As a result, for lineages above the median the FDR is 4.3%, indicating, as expected, that our procedure captures the higher-fitness lineages very well.

#### Pre-existing mutations

Some mutations could have arisen prior to the beginning of the experiment, specifically, prior to splitting the barcoded yeast library between five A and five C replicates. Such pre-existing mutations cause two issues with the downstream analysis and interpretation. First, they make it more difficult to identify causal mutations that increase fitness of adapted lineages. We discuss this problem and how we address it in Section 3. The second issue is that pre-existing beneficial mutations confound the inference of bDFEs and their comparison across treatments. Specifically, if multiple lineages (within the same population or in different replicate populations) carry the same pre-exisiting beneficial mutation and are identified as such, this mutation will be counted multiple times and will therefore inflate the size of a fitness class in the inferred bDFE.

To diagnose this problem, we note that there are two types of pre-existing mutations. Those that arose after the transformation that introduced barcodes into our strain (type I) and those that arose before this transformation (type II). All pre-existing mutations of type I are linked to different barcodes. Therefore, these mutations can be identified by comparing barcodes of lineages identified as adapted in different populations. We found that pairs of populations have on average 416 (15%) adapted lineages with a common barcode, whereas only 23 (0.8%) would be expected by chance (Figure S8). Barcodes are more often shared between A populations than other types of population pairs (22% vs 13%, t-test *P* = 10^−10^). We suspect that this is because the majority of adapted mutants have higher relative fitness in the A-condition than the C-condition (Figure 2B in the main text), such that pre-existing mutations are more likely to reach high frequencies and be detected in the A-condition. We eliminate pre-existing beneficial mutations of type I from our bDFE analysis by constructing these distributions from lineages with unique barcodes.

The same pre-existing mutation of type II can be linked to multiple barcodes and is indistinguishable in the barcode data from multiple independently arising mutations. However, pre-existing mutations of type II can be identified in the genome sequencing data. As discussed in the methods section on genome sequencing and analysis, we find no evidence of such mutations, suggesting that they must be rare in our populations.

#### Additional analyses

The median standard error in our estimates of fitness of adaptive mutants was 0.02, and median CV was 0.08 (Figure S1A). The standard deviation of the bDFEs is much larger, 0.22 for A populations and 0.33 for C populations, suggesting that there are multiple classes of adaptive mutations available to yeast in both conditions. We confirm that the observation of a multimodal bDFE is not an artefact of combining the bDFEs of all replicate C populations (Figure S6). Furthermore, the fitness estimates have fairly consistent error distributions across all replicate populations (Figure S7).

## 2 Competitive fitness assays

We obtained barcode frequency data at cycles 1, 2, 3, 4 and 5 (Figure S9). Fitness estimates of individual lineages were concordant between replicates (Figure S10), with relatively low estimation errors (Figure S11). Estimated fitness of sampled clones in both conditions along with errors and confidence intervals are provided in Data S2.

### Determining valid clones for further analysis

Of the 581 clones that were pooled in the competition assays, we filtered out many clones from downstream analysis via a number of filters. First, clones that were sampled at cycle 17 were excluded from further analysis, as were clones that lacked a valid fitness measurement in either environment or had variance in fitness measurement in either environment ≥ 0.5. These filters removed 151 of 581 from consideration, leaving us with 430 clones for further analysis.

### Calling adapted clones

We use the competition assay data to call adapted clones. A clone is called as adapted in a given environment if its estimated competitive fitness in that environment is more than two standard errors greater than 1 (2.3% FDR based on normal distribution). We identified 401 clones as adapted in the A-condition (see Data S2). 221 of them were sampled from the A-condition and we refer to them as the A-mutants. We identified 402 clones as adapted in the C-condition (see Data S2). 189 of them were sampled from the C-condition and we refer to them as the C-mutants. There are 16 clones that are not adapted in either A or C condition, 13 clones that are adapted in the C-condition but not in the A-condition and 12 clones that are adapted in the A-condition but not in the C-condition. Of the 16 clones not adapted in either the A or C conditions, 12 come from lineages determined not to be adaptive in the BLT analysis, which we define as our neutral clones (Data S2). The genomes of 215 A-mutants, 181 C-mutants and eight neutral mutants were later sequenced, as described in the Methods and Section 3.

### Relationship between fitness estimated in competition assays and in BLT experiments

The correlation between fitness of adapted mutants (i.e., A- and C-mutants) estimated in the competition assays and those estimated from the BLT data is reasonably good (Figure S12, *R* = 0.57, *P* = 8.77 × 10^−37^), but there are also some systematic differences. Specifically, competition assays over-estimate BLT fitness in the A-condition and under-estimate it in the C-condition (Figure S12). We discuss possible reasons for these discrepancies below.

When we place mutants onto the bDFE in the non-home environment (as in Figure 2A in the main text) and when we estimate the probability of sampling a mutation in the non-home environment (see Section 4), we need to know the BLT fitness of mutants in their non-home environment. We have no direct measurements of BLT fitness of A- or C-mutants in their non-home environment, but we do have the non-home fitness estimates of all mutants in competition assays. However, directly substituting BLT fitness for competitive fitness would lead to biases due to the aforementioned discrepancies between the two estimates. To correct for these discrepancies, we linearly regress BLT fitness of A-mutants against their competitive fitness in the A-condition and we regress BLT fitness of C-mutants against their competitive fitness in the C-condition (see Figure S12). We then use these regressions to estimate the BLT fitness of C-mutants in the A-condition and A-mutants in the C-condition.

### Difference in adaptive mutant fitness between conditions

Mutant’s fitness were called as significantly different between A and C-conditions if the two 95% confidence intervals did not overlap. We discovered 178 such clones (108 A-mutants and 70 C-mutants), all of which were adaptive in both environments.

### Possible reasons for the slight discrepancy between BLT and competitive fitness estimates

We can think of at least three possible reasons for this discrepancy. One possible reason is that we use a different set of reference lineages in the BLT experiments and competition assays. It is possible, for example, that some of the lineages that we use as reference in the competition assay are not in fact neutral in one or both of the conditions.

Another possible reason is that fitness is weakly frequency-dependent (this is expected in batch culture experiments [9]). Frequency dependence can manifest itself in a discrepancy between BLT and competition assays because lineages are present at much higher frequencies in the competition assays than in the BLT experiments. Specifically, the median frequency of an adapted clone (as defined from the competition analysis) in the competition assay is 4 × 10^−4^ at the initial time point, whereas the median frequency of an adapted lineage is 3 × 10^−6^ at the initial time point (the frequency of the adapted mutant driving the lineage frequency must be even lower since many if not most adapted lineage also initially contain non-adapted individuals).

A third possible reason is that the media composition in the BLT experiments and in competition assays could be somewhat different at later stages of the growth cycle because population compositions are different and the media composition is determined by the collective metabolism of all variants present in the population.

Given the overall concordance of fitness estimates, we decided that dissecting these relatively minor effects was not particularly important within the scope of the present work.

## 3 Genome sequencing and analysis

### Small mutations

The distributions of derived small variants per clone is shown in Figure S16A and their summary statistics are given in Table S2. These data show that the majority of small variants detected in the evolved clones are still likely ancestral or erroneous. We can estimate the expected number of adaptive small variants as follows. A typical adapted isolate carries 1.18 more mutations compared to an ancestral isolate (Figures S15, S16, Table S2). However, not all of these mutations may be adaptive. Indeed, given that small indels and single-nucleotide mutations occur at rate 3 × 10^−3^ per genome per generation [10], we expect 0.18 of such mutations to have occurred on the line of descent of any isolate sampled at cycle 9 (~ 60 generations). Based on this estimate, a typical adapted clone is expected to carry 1.18 – 0.18 = 1.00 adaptive mutation.

One potential problem with this estimate is that our yeast strain may have a somewhat different mutation rate than the strain used in Ref. [10]. We can obtain an alternative estimate for the number of adaptive mutations per clone by comparing adapted clones with the neutral ones. We find that a typical neutral clone carries 1.96 ± 0.88 more mutations than a typical ancestral isolate^1^ (Table S2), which is greater than 1.18 extra mutations carried by a typical adapted clone. Thus, it is possible that a typical adapted clone carries no beneficial small mutations.

The problem with the second estimate is that some of the clones identified as neutral may in fact carry adaptive mutations^2^. Therefore, we use both methods to obtain bounds on the number of adaptive mutations carried by a typical adapted clone and conclude that a typical adapted clone is expected to carry between 0 and 1 small adaptive mutations. Carrying out the same calculations for the A- and C-mutants separately, we estimate that a typical A-mutant carries between 0 and 1.34 small adaptive mutations (so that between 0 and 288 out of 1008 small mutations found in A-mutants are expected to be adaptive), and a typical C-mutant carries between 0 and 0.61 small adaptive mutations (so that between 0 and 110 out of 717 small mutations found in C-mutants are expected to be adaptive; Table S2).

Using genetic parallelism, we identify 185 mutations at 63 loci as adaptive (on average 0.47 mutations per clone), consistent with our expectation. 76 of these mutations at 39 loci are found in 60 A-mutants (on average 0.35 mutations per clone) and 109 mutations at 56 loci are found in 81 C-mutants (on average 0.60 mutations per clone; Table S2).

### Total number of identified adaptive mutations per clone

After combining small mutations and CNVs together, we find that a typical A-mutant carries 0.74 identified adaptive mutations with 100 A-mutants having no identified adaptive mutations (Data S3). A typical C-mutant carries 1.11 identified adaptive mutations with 46 C-mutants having no identified adaptive mutations (Data S3). Since all A- and C-mutants gained fitness in their home environment, each of them must have at least one adaptive mutation. Therefore, the “unknown” sectors in Figure 2C in the main text refers to the numbers of A- or C-mutants without any identified adaptive mutations.

### Pre-existing mutations

We use genetic information to test for the prevalence of preexisting mutations of type II, i.e., those that arose prior to the integration of the DNA barcodes, so that multiple lineages may carry an adaptive mutation identical by descent. If such mutations were prevalent, we would expect to observe an excess of adapted clones carrying identical genetic mutations, compared to ancestral isolates, but this is not the case (Figure S15). In fact, an identical small mutation is found less frequently in multiple adapted clones than in multiple ancestral clones, suggesting that some of the pre-existing genetic variation may have been deleterious.

CNV events are not suitable for this analysis because they occur at very high rates, and thus we have no way of ascertaining whether or not two identical CNV events are identical by descent. Indeed, many CNVs are aneuploidies, which occur at rate 9.7 × 10^−4^ per generation [10]. The only consistent segmental CNVs are ChrIV-1n and ChrIV-3n. ChrIV-1n mutations are localized to identical breakpoints across 64 mutants; these breakpoints are concordant with highly similar repetitive regions, YDRWTy2-2/YDRCTy1-2 and YDRWTy2-3/YDRCTy1-3. There are three different classes of ChrIV-3n amplifications across 10 mutants; at least some of these breakpoints also appear to be localized close to known repetitive regions, including YDRCTy2-1, YDRCTy1-1 and YDRWTy2-2. Thus, we suspect that these recombination-driven segmental events also occur at high rates and are not necessarily indicative of pre-existing variation.

## 4 Simulations of evolutionary dynamics and the estimation of rates of adaptive mutations

We found that the sets of A- and C-mutants are genetically distinct (Figures 2C in the main text, Extended Data Figure 4 and Data S4), despite all of them being more fit than the ancestor in both environments. In this section, we show that this somewhat puzzling observation can be explained by the differences in the evolutionary dynamics in the concurrent mutation regime [11].

There are two key differences between the A- and C-conditions. First, the bDFEs are different (Figure 2A in the main text), resulting in different increases of population’s mean fitness over time. Second, the fitness rank orders of adaptive mutations are also different (Figure 2A in the main text). These two facts imply that in the concurrent mutation regime the chances for a given mutation to escape drift while rare and reach a certain frequency and be sampled can be substantially different in the two conditions. Next, we develop a quantitative version of this argument.

As described in Section 3, we classify all discovered adaptive mutations into mutation classes by the type of CNV or the gene in or near which the mutation occurred (Data S4). Let 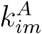 and 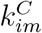 be the number of sequenced A- or C-mutants that carry a mutation from class *m* sampled from the replicate population *i* = 1, 2, 3, 4, 5 (see Table S3). We would like to know whether the differences between 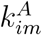 and 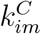 can be explained by the observed differences in the bDFEs and by differences in the fitness benefits provided by mutations of class *m* in the A- and C-conditions.

The challenge is that the probability Pr {*k*; *s, U*} of observing *k* mutants of a certain type in a sample from a given population depends not only on the selection coefficients s of adaptive mutations of this type (which we have measured, as described in Section 2) but also on the unknown rate *U* at which such mutations arise. Therefore, we first find the mutation rates that fit our data and confirm that these rates are biologically plausible. We then ask how well the expected numbers of A- and C-mutants carrying specific mutations match the observed numbers 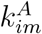 and 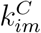, given the estimated mutation rates.

### 4.1 Data

#### Exclusion of pre-existing mutations

A number of lineages are identified as adapted in multiple populations, indicative of pre-existing mutations of type I (see Methods for more details). Since such mutations do not carry information about the mutation rate, we exclude them from this analysis. That is, numbers 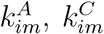 given in Table S3 and in Extended Data Figure 5 are the numbers of sampled and sequenced clones with unique barcodes.

#### Selection coefficients

Fitness of individual mutants in the BLT experiments are estimated based on the competition assay data, as described above (see Section 2), and selection coefficients 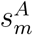 and 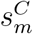 are taken as averages over all mutants carrying a mutation from a given mutation class (Table S3).

#### Mean fitness trajectories

Our model described below takes into account the fact that the probability of a newly arisen beneficial mutation to survive genetic drift depends on how the mean fitness of the population changes over time. Thus, when simulating the evolutionary dynamics, we use the empirical mean selection coefficient trajectories 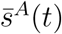 and 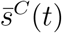 in the A- and C-conditions, respectively. 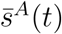 and 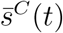 are computed by first averaging the values of the mean selection coefficient (estimated in as described in the Methods) over all five replicates and then fitting the logistic function to these data points (see Figure S1B).

### 4.2 Model

To keep the model as simple as possible, we assume that our population has the constant size *N* = 2 × 10^6^ in both A- and C-conditions, mutations from the mutation class *m* arise at (an unknown) rate *U_m_* per division in both conditions. We assume that all mutation from class *m* confer the same (known) fitness benefits 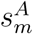 and 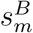 relative to the ancestor in the A- and C-condition, respectively (Table S3). We also assume that all mutants are sampled at time *T* = 60 generations after the beginning of the experiments in both A- and C-conditions, which roughly corresponds to cycle 9, assuming log_2_ 100 = 6.64 doublings per cycle (see Methods).

To estimate the mutation rate *U_m_* for mutation class *m* we define the log-likelihood function

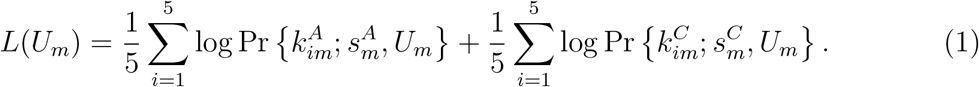

The sampling probability Pr *{k; s, U*} depends on the number of lineages that independently arose in the population and that carry a mutation of the focal type, as well as on the frequencies of these lineages at the sampling time point *T*. It is difficult to obtain an analytical expression for this probability because it requires integrating over all these nuisance parameters. We therefore compute the sampling probability numerically, using population dynamic simulations described below.

#### 4.2.1 Evolutionary dynamics simulations and the estimation of the sampling probability

As mentioned above, difference in the bDFE between the A- and C-conditions may contribute to the observed differences in the genetic composition of sampled mutants. The bDFE indirectly affects the survival probability of a newly arisen beneficial mutation by altering how the mean fitness of the population changes over time. Thus, we design our simulations so that they match the average empirical mean selection coefficient trajectories in the A- or C-condition (see Methods and Figure S1B). Matching these trajectories in simulations of the full Wright-Fisher where all mutation classes segregate simultaneously is difficult. Instead, we simulate the arrival and spread of mutations of each mutation class separately, while accounting for changes in mean fitness using our logistic fit of the observed mean fitness trajectories (Figure S1B). This approach also allows us to use branching process approximations described below in Section 4.2.2, which greatly speed up the calculations.

Since in this section we are concerned with mutants of one mutational class in one environment, we omit the subscript *m* and superscripts A/C. In other words, we assume that the mutations arrive at rate *U* per individual per generation and have selection coefficient *s*, and we sample 88 mutants from this population at time point *T* = 60.

In our simulations, we allow for new mutations to arise between 1 cycle prior to the beginning of the experiment up to cycle 9. Thus, we divide the time interval between –6.6 and *T* = 60 generations into 260 to 1300 segments of length Δ*t* = 0.01/*s* generations. For each time segment (*t_i_*, *t_i_* + Δ*t*), we draw the number of new mutants arising in the population in that segment from the Poisson distribution with rate *N U* Δ*t*. Each of these mutants survives until the sampling time point *T* with probability *P*_surv_(*T; s, t_i_*), given by equation (2) below. For each mutant that survives, we draw its establishment time *τ* from the distribution given by equation (3) below. We then set the mutant’s frequency at the sampling time point to *n*(*T*; *s_m_, τ*)/*N*, where *n* is given by equation (4) below. At the end of each simulation run, we obtain a list of frequencies of all independently arisen mutants carrying mutation of type *m*. We assume that each independently arisen mutants is linked to a distinct barcode. We then randomly sample 88 clones from the whole population and discard those that do not carry the focal adaptive mutation. If multiple clones are sampled from the same lineage, only one is retained. We further randomly sub-sample these clones with 97.5% success probability, simulating the small whole-genome sequencing failure rate, which results in the final number *k* of sampled clones that carry a mutation from the focal mutation class. We estimate the probability of sampling *k* clones, Pr {*k; s, U*}, by running 10^4^ simulations and recording the fractions of simulations where we observe *k* sampled clones.

#### 4.2.2 Branching process approximations for mutant growth dynamics

Consider a mutant that arises at some point *t*_0_ < *T* on the ancestral background. The early population dynamics of such mutant lineage can be modeled as a branching process [11]. In particular, Desai and Fisher [11] used the branching process approximation to derive the probability that a mutant with selection coefficient s that arose at time zero in the ancestral population (i.e., whose mean selection coefficient is zero) has not gone extinct by time *t* and the related probability that the mutant “establishes” at time *τ* (see equations (11) and (17) in Ref. [11]). However, this model ignores the fact that the mean fitness of the population is changing over time while the focal mutant is still at low frequency. In our experiment, mean fitness changes very rapidly, as can be seen in Figure S1B, which can significantly alter mutant’s survival probability and its establishment time. Assuming that the mean selection coefficient trajectory 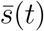 is known (see Section 4.1), we can model mutant dynamics analogous to Ref. [11] but with a generalized birth-death model with growth rate 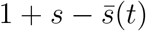 and death rate 1. In this model, the probability that a mutant that arose at time *t*_0_ survives until time *t* is given by

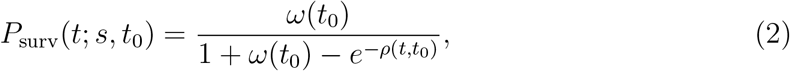

where

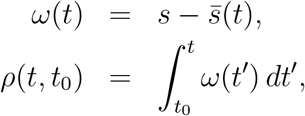

provided that *ω*(*t*_0_) > 0. Conditional on surviving, the probability that this mutant establishes at time *τ* is given by

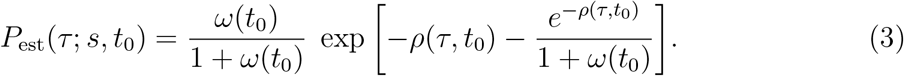

Once the mutant establishes at time *τ* and provided that it is still beneficial, i.e., *ω*(*τ*) > 0, its subsequent dynamics are essentially deterministic, so that its size *n*(*t*; *τ*) at time *t* is approximately given by

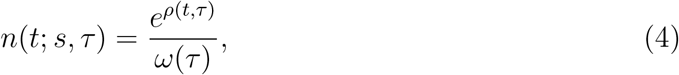

which is analogous to equation (14) in Ref. [11] in the limit *t* ≫ 1/*s*.

#### 4.2.3 Estimation of mutation rates by maximum likelihood

For each mutation class *m*, we compute the likelihood function *L*(*U_m_*) using equation (1) for 40 discrete values of *u_m_* = log_10_ *U_m_* between –7 and –3 and interpolate between them using a polynomial of degree 4. The likelihood function for chrIII-3n mutations is shown as an example in Figure S17. We find the maximum of this function as a point where the polynomial approximation of L has a zero first derivative with respect to *u_m_*. We estimate the standard error (SE) as the inverse of the square root of Fisher information [12].

### 4.3 Results

We estimated the rates of driver mutations at 9 loci that were mutated in at least five adapted clones, after excluding gene NUM1 where all mutations are likely pre-existing (see Extended Data Figure 4, Table S3 and Data S4). Our estimates are generally consistent with those derived from a published mutation accumulation experiment by Zhu et al [10]. In particular, our estimate of the rate of small adaptive mutations in genes HEM1 and HEM2 is ~ 3 × 10^−7^ per generation, consistent with per basepair mutation rate of 1.67× 10^−10^ estimated by Zhu et al [10]. Gene YIL169C is an exception, with an estimated rate of adaptive point mutations of 7.76 × 10^−6^ per generation. We estimate the rates of large CNV events to be ~ 10^−5^ per generation, again consistent with the genome-wide rate of aneuploidies of 1.04 × 10^−4^ estimated by Zhu et al [10]. Our estimate of the segmental duplications ChrIV-3n is much lower (3.24 × 10^−7^ per generation), also consistent with Ref. [10], although they do not provide a quantitative estimate of such events.

The probability that at least one sequenced mutant from all five replicate populations has a mutation with a given selection coefficient s and mutation rate *U* is shown in Extended Data Figure 5A,B, for both A- and C-conditions. As expected, this function is different between conditions because the mean fitness dynamics are different (Figure S1B). Finally, we directly compare the observed numbers of mutations of each mutation class with the expected numbers in the A- and C-conditions and find a reasonable match (Extended Data Figure 5C).

## 5 Phenotyping

### 5.1 Calibration curve for the measurement of alga cell density

We generated a calibration curve for interpreting fluorescence measurements in terms of alga cell number as follows. We grew alga in our standard growth conditions both alone and in a community with the ancestral yeast. On each day of the growth cycle, we counted alga cells using a haemocytometer and we took a fluorescence measurement using the a plate reader, as described in Methods. In addition, we generated a 2-fold dilution series starting with the saturated alga cultures at the end of the growth cycle both alone and in the community. We also obtained haemocytometer counts and fluorescence measurements for the alga grown in a 59 mutant communities (see Methods in the main text). We combined all these data to obtain a single the calibration curve by plotting the fluorescence against haemocytometer counts, both log-transformed (Figure S18). We found a good positive correlation (*R* = 0.96). We used the following equation to convert fluorescence measurements into cell density,

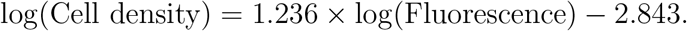

### 5.2 Analysis of plausibility of natural selection favoring high- *K*/low-*r* mutants

Our measurements of *r* and *K* for the adapted yeast mutants suggest that the increases in *K* at the expense of *r* and increases in *r* at the expense of *K* can both be adaptive in both A- and C-conditions. Theory suggests that high-*K* mutants can be favored by natural selection when the population is close to starvation [13, 14], but mutants with higher *K*, and especially those with lower *r*, are rarely found in evolution experiments. We therefore wanted to test whether the mutants with our measured *r* and *K* values (especially those with higher *K* and lower *r* than the wildtype) can plausibly invade the wildtype yeast population.

To this end, we constructed a simple coupled logistic growth model. We consider two strains, strain 1 (wildtype) and strain 2 (mutant) whose per capita growth rates are *r_i_* and carrying capacities are *K_i_*, *i* = 1, 2. The dynamics of the population sizes *N_i_* (*i* = 1, 2) are then described by equations

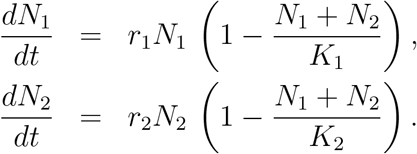

We set *r*_1_ = 0.168 h^−1^ and *K*_1_ = 5.5 × 10^6^ for the wildtype and we set the mutant values *r*_2_ and *K*_2_ based on our measurements given in Data S2. We set the initial population sizes of the wildtype and mutants to be 10^4^ and 10^2^ individuals, respectively, and simulate the growth of these strains for 120 hours (the length of our standard growth cycle). We determine that a mutant can successfully invade if its frequency increases by the end of the growth cycle.

If our model perfectly captured our experimental conditions, all of the mutants would be able to invade, since all of them are experimentally found to be more fit than the wildtype. In fact, we find that only 37 out of 59 (63%) mutants are able to invade (Figure S22). As expected, five mutants that have both *r* and *K* lower than the wildtype could not invade in our model. Similarly, eight mutants with higher *K* but low *r* and nine mutants with higher *r* and lower *K* could not invade, suggesting that our model does not capture some processes and traits that are likely important in our experiment (e.g., it does not capture the dependence of growth and death rates on resource concentrations). Nevertheless, 16 out of 24 mutants with higher *K* and lower *r* are able to invade the wildtype in this simple model, which supports our argument that selection can favor high *K* in our conditions even if it comes at the expense of decreasing *r*.

### 5.3 Relationship between *r*, *K* and fitness

As described in the main text, we found that each *r* or *K* individually explain about 26% of variation in competitive fitness in the C-condition but do not explain any statistically significant variation in the A-condition. However, it is possible that a linear combination of both variables would improve our ability to predict fitness in both conditions. To this end, we consider a multiple regression model Fitness ~ *r* + *K*. We report the results of this regression analysis in Table S4. We find that *r* and *K* jointly do not explain any variation in fitness in the A-condition, but they jointly explain 37% of variation in fitness in the C-conditions, which is significantly more than each of them explains individually. Both *r* and *K* contribute approximately equally to fitness. Furthermore, we find that decreasing *r* increases fitness even if *K* is being held constant. This surprising observation could be explained if one or more unobserved traits (other than *r* and *K*) were important for competitive fitness and if there was a trade-off between *r* and such unobserved trait.

## 6 Possible function effects of mutations in *HEM1*, *HEM2* and *HEM3* genes

Three critical components of the heme biosynthesis pathway, HEM1, HEM2 and HEM3 are putative targets of adaptation in this study, and provide a substantially larger fitness benefit in the C-condition than in the A-condition (see Data S3 and Data S4). Most mutations found in these genes probably compromise the function of the encoded enzymes and lead to a decreased “siphoning” of succinyl-coenzyme A (sCoA) from the TCA cycle and the production of less heme and/or decoupling of aerobic-driven regulation from aspects of central metabolism. Although yeast is already capable of performing fermentation under ambient oxygen concentration, we speculate that these mutations enables yeast to ferment under even higher oxygen concentrations that results from alga photosynthesis.

### 6.1 HEM1

*HEM1* encodes the mitochondrial 5-aminolevulinate synthase (ALAS) that catalyzes the first committed step of porphyrin through the condensation of glycine and sCoA using the cofactor pyridoxal 5’-phosphate (PLP):

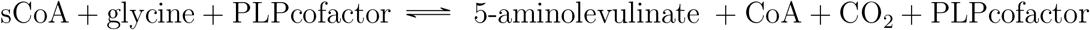

ALAS links heme/cytochrome production with the TCA cycle and aerobic respiration via sCoA, and thus plays a key role in cellular energetics [15]. ALAS functions as a homodimer (with subunits referred to as A and B below). A crystal structure of *S. cerevisiae* HEM1 is available (https://www.rcsb.org/3d-view/5TXR/1).

We found 5 adaptive mutations in the *HEM1* gene, four of which occurred in the C-mutants (Data S3 and Data S4). A nonsense mutation at residue 166 (out of 548) in HEM1/ALAS leads to a truncated coding sequence and presumably loss of function. All other mutations alter amino acids that are conserved in all sequenced *S. cerevisiae* strains, which we speculate may also significantly compromise normal HEM1/ALAS function, as described below.

**His107Pro** mutation does not seem to be involved in any substrate or co-factor binding directly, but a change from histidine to a proline, with significant backbone confirmational constraints may have serious consequences for the folding (and thus function) of the enzyme. Bracketing this position (107) nearby is Arg91, which plays a critical role in sCoA binding, and Asn121, which forms an alpha-carboxylate hydrogen bond with the glycine substrate [16]. Moreover, His107 (of subunit B) forms important structurally stabilizing side-chain interactions with Glu111 (subunit B) and Lys142 (subunit A) (see Figure S3 of [16]), which may very well be disrupted with a proline substitution (lacking a side chain).

**Asn152Lys and Asn157Lys** mutations occur in a region of the ALAS protein that becomes ordered upon PLP cofactor binding. N152 plays a direct role in coordinating sCoA [16], hydrogen bonding with the carboxylate (COO-) moiety of the sCoA succinyl group. Mutation from asparagine to a positively charged lysine may both disrupt the formation of this hydrogen bond and prevent a key side-chain interaction between Asn152 (subunit A) and Arg91 (subunit B) that stabilizes key structural elements needed for PLP cofactor binding (we expect the side chain of mutation Lys152 to repel that of Arg91 since both are positively charged). Asn157Lys is a few amino acids downstream from Ile153, which also plays a role in carboxylate/sCoA coordination like Asn121 [16]. Asn157 is adjacent to Ile158 (subunit A) that forms stabilizing main-chain and side-chain interactions upon PLP cofactor binding with Asn95 and Asn 97 (both on subunit B); mutation to Lys may disrupt these interactions, along with that of nearby stabilizing interactions between Arg98 (subunit B) and Ala147 (subunit A), due to positive charge repulsion between Arg98 and Asn157Lys (see Figure S3 of [16]).

**Gly344Cys** mutation occurs at a site that does not directly bind substrate or co-factor but is each bracketed by amino acids that do play key active site roles and could also have structural consequences as conformational flexibility is likely lost with the change away from glycine. Gly344 is flanked (although several amino acids away) by K337 that forms a critical covalent pyridoxyl-lysine bond, and F365 which delineates the sCoA substrate-binding pocket [16]. Gly344 resides on the juncture between a beta-sheet and alpha-helix motif in the protein structure, with little space for a side chain. Mutation of this site to a cysteine may alter backbone flexibility and the folding of key structural elements. Gly344 is also in close proximity to Cys182 that forms stabilizing hydrogen bonds with amino acids Asn129, Thr275, and Gly 276; having another cysteine nearby may very likely interfere with that hydrogen bonding network.

### 6.2 HEM2

HEM2 encodes for the cytoplasmic/nuclear, homo-octameric aminolevulinate dehydratase (ALAD; porphobilinogen synthase), which catalyzes the second step in heme biosynthesis:

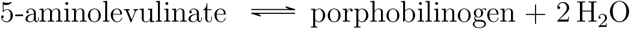

A crystal structure of *S. cerevisiae* HEM2 is also available (https://www.rcsb.org/3d-view/1AW5/1).

We found 5 adaptive mutations in the *HEM2* gene, four of which also occurred in the C-mutants (Data S3 and Data S4). Two frame-shift mutations lead to premature stop codons ~ 100 out of 342 amino acids. The three remaining mutations Ile129Phe, Val132Phe, and Pro264Arg are predicted to have moderate impact.

**Ile129** resides on a beta-strand with a side chain embedded in a relatively hydrophobic environment. It is unclear what the consequences of mutation Ile129Phe might be, as phenylalanine is a comparably bulky hydrophobic side chain to isoleucine that would at first glance seem to be a conservative substitution [17]. However, several aromatic amino acids are in the vicinity, including Trp30 and Tyr127; Ile129Phe may form aromatic ring-stacking interactions with these residues to disrupt fold structure.

**Val132** is similarly on a beta-strand with a side chain embedded in another nearby hydrophobic pocket and interface with other non-polar amino acids on alpha-helix. The Val132Phe mutation could disrupt the tight packing at this interface as phenylalanine is substantially bulkier than valine. Moreover, Tyr168 is nearby which hydrogen bonds a key water molecule that is hydrogen-bonded to several other backbone atoms flanking Val132; the Val132Phe mutation may also disrupt this by interacting with Tyr168 through aromatic stacking interactions. Pro264 resides in a sharply kinked beta-turn between beta-strand and alpha-helix elements, which may be important for constraining and enabling key interactions of flanking residues with amino acids distributed across different structural elements: Val262/Ser265/Tyr287, Ser265/Glu292, and Lys263/Tyr287 (see https://www.rcsb.org/3d-view/1AW5/1).

**Pro264Arg** mutation is pronounced not just for the loss of backbone constraint provided by the imino acid proline [17], but it introduces a large positively charged side change that may form “inappropriate” interactions to disrupt monomer fold and subsequently oligomerization; these include interactions with nearby: negatively charged amino acids: Glu292 and Glu313 aromatic amino acids (through cation-pi interactions) Phe211 (normally interacting with Tyr268 via aromatic stacking), Tyr216 (normally interacting with the backbone of Phe211), Tyr268 (normally interacting with Glu313), and Tyr287 (normally interacting with Lys263 via cation-pi interactions).

### 6.3 HEM3

*HEM3* encodes for the cytoplasmic/nuclear enzyme, porphobilinogen deaminase/hydroxymethylbilane synthase (HMBS), which catalyzes the third step in heme biosynthesis involving four molecules of porphobilinogen to make a linear hydroxymethybilane molecule that looks like a heme/tetrapyrole when “wrapped around”:

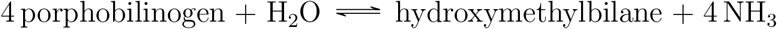

We found 4 adaptive mutations in the *HEM3* gene, three of which occurred in the C-mutants (Data S3 and Data S4). However, since no structure exists for *S. cerevisiae* HEM3 protein and since the homology with the human ortholog for which the structure does exist (https://www.rcsb.org/3d-view/5M6R/1) is rather low (~ 40% similarity), predicting the functional effects of these mutations is challenging.

## 7 Data availability

All raw sequencing data is available on the US National Center for Biotechnology Information (NCBI) Sequence Read Archive (SRA) under BioProject PRJNA735257. Other input data (e.g. growth data, variant calls, community yield etc) and analysis scripts can be found on Dryad at https://doi.org/10.6076/D14K5X.

The latest version of the barcode counting software BarcodeCounter2 can be found at https://github.com/sandeepvenkataram/BarcodeCounter2.git.

## 8 Supplementary Figures

**Figure S1.**
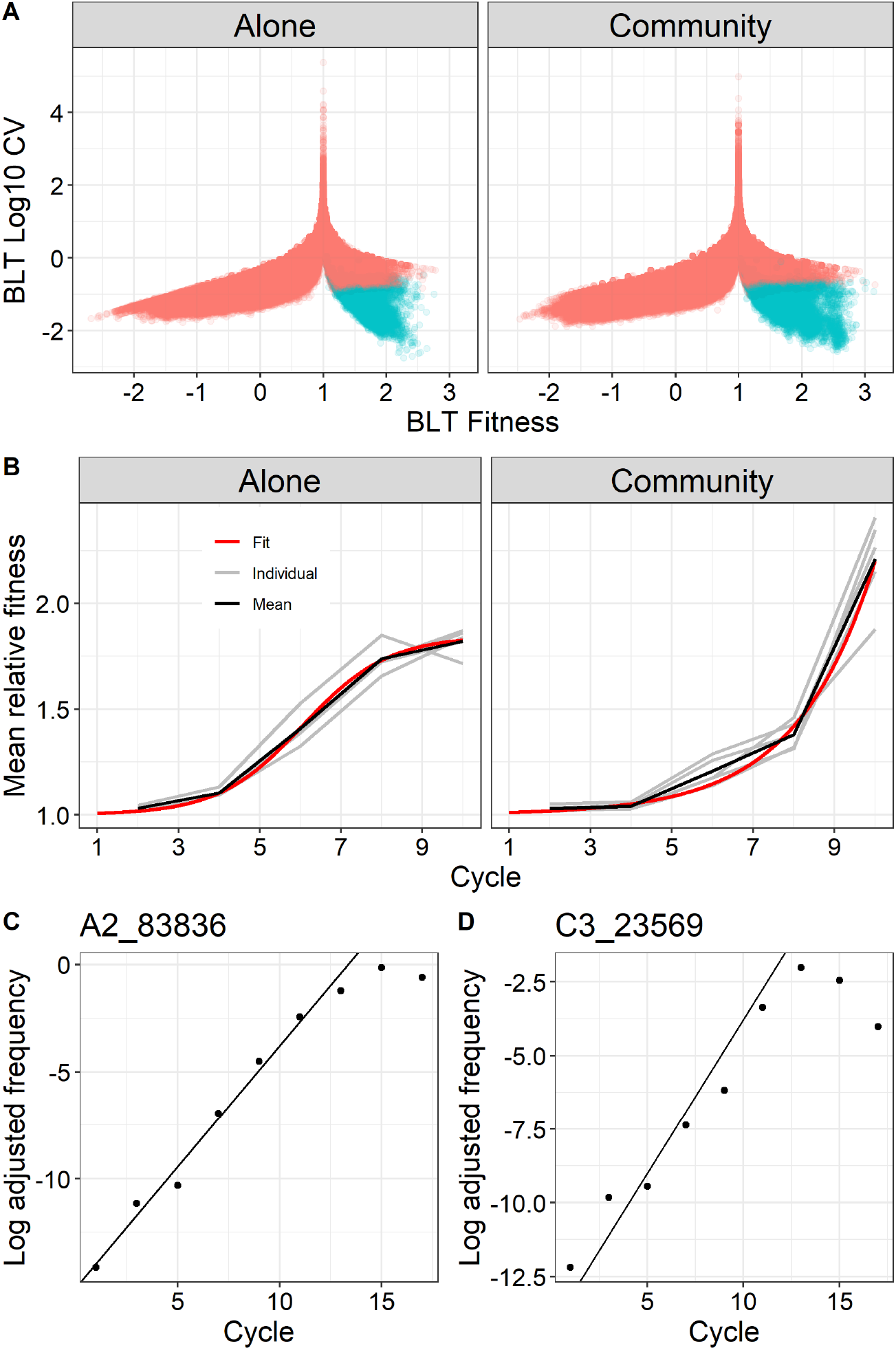
Estimation of fitness of adapted lineages in the BLT data. **A.** The mean lineage fitness (*x*-axis) is plotted against the coefficient of variation (CV) of the estimate (*y*-axis). Blue (red) points indicate lineages identified as adapted (non-adapted). **B.** Mean fitness of each population over time. Individual populations are shown in grey, the average mean fitness trajectory over all populations is shown in black, and a logistic fit to this average trajectory is shown in red. **C, D.** Examples of selection coefficient estimation for one lineage in one A population (C) and one C population (D).

**Figure S2.**
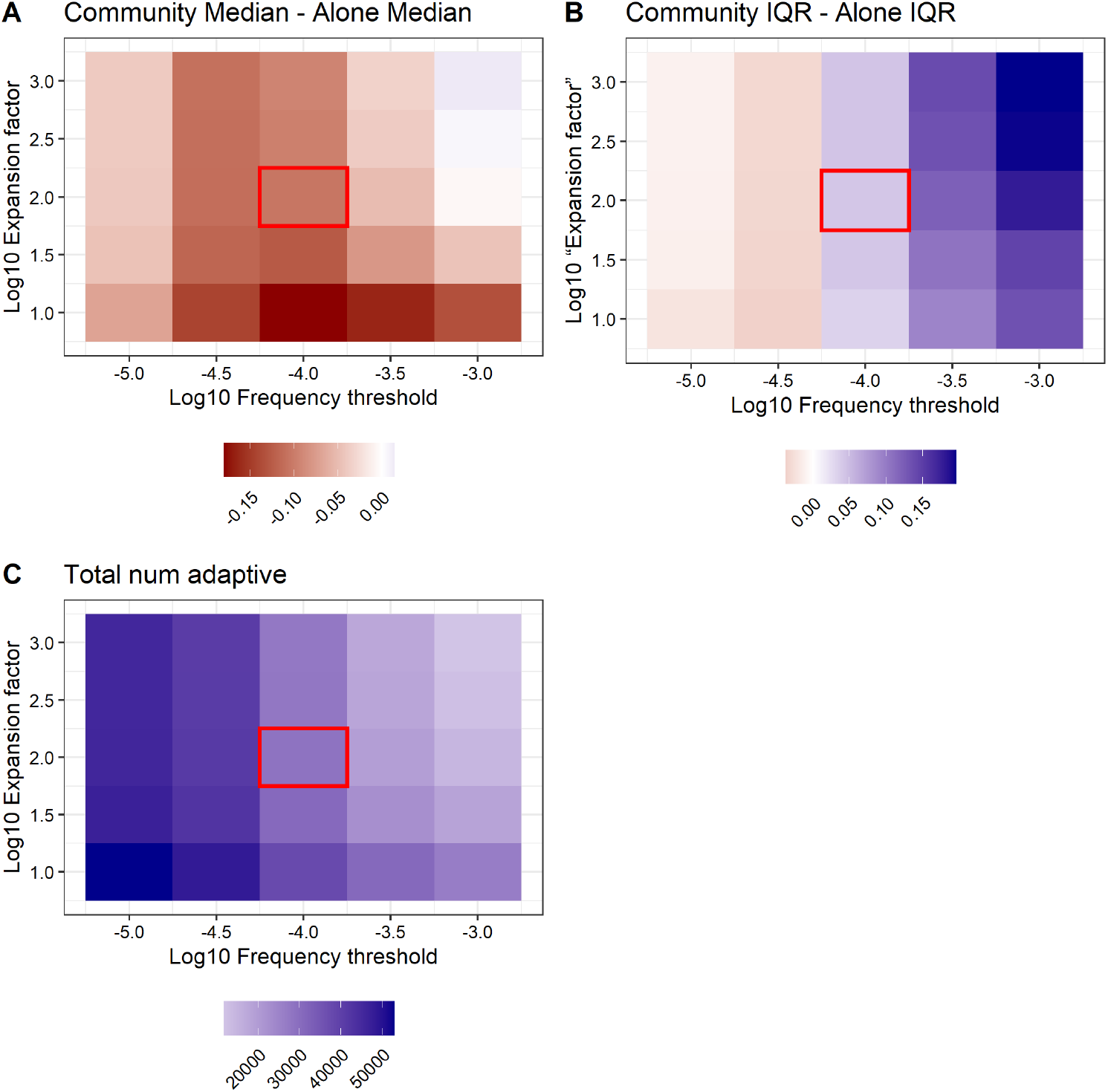
Analysis of sensitivity of the procedure for calling adapted lineages with respect to the minimum frequency threshold and the expansion factor. **A.** Sensitivity of the difference in the bDFE means. **B.** Sensitivity of the difference in the bDFE interquartile interval (IQR). **C.** Number of called adapted lineages as a function of the minimum frequency threshold and the expansion factor. The red square on each panel highlights the parameter regime used for all of our remaining analyses.

**Figure S3.**
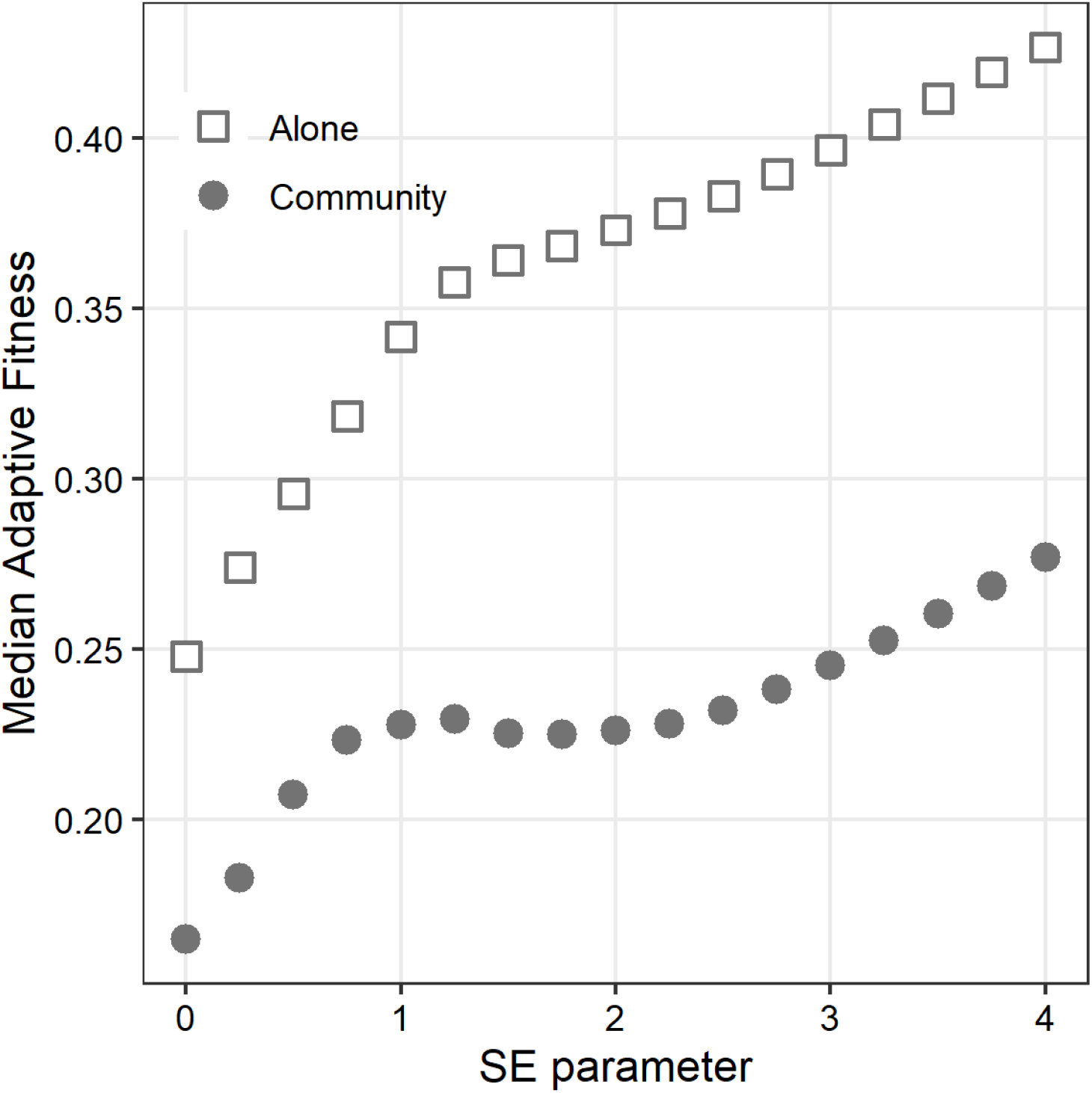
Analysis of sensitivity of the procedure for calling adapted lineages with respect to the SE parameter. Median of the pooled bDFE as a function of the SE parameter.

**Figure S4.**
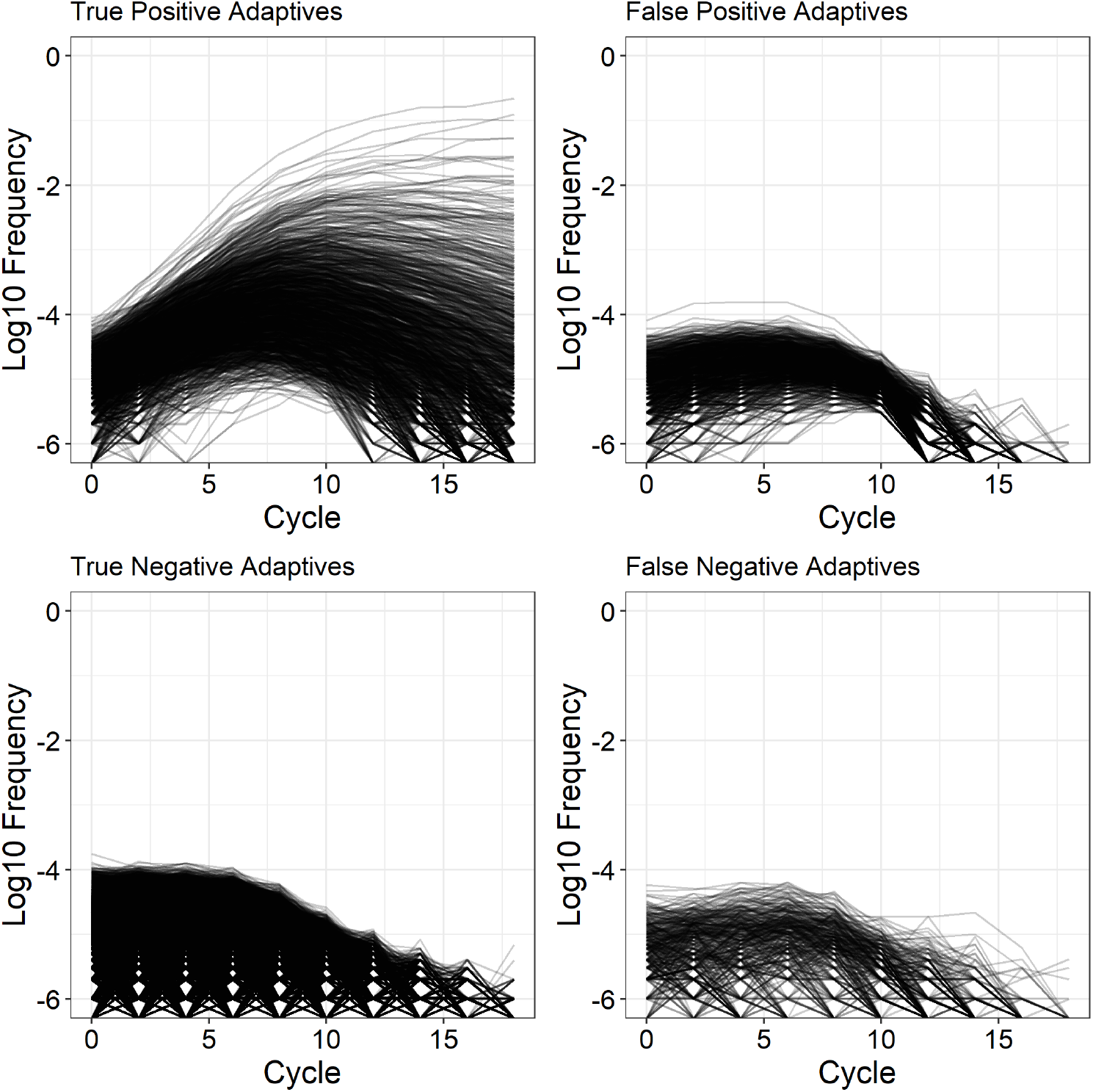
Precision and accuracy of the procedure for calling adapted lineages. The panels show the frequency trajectories of simulated lineages classified by our heuristic procedure (see Section 1 for details).

**Figure S5.**
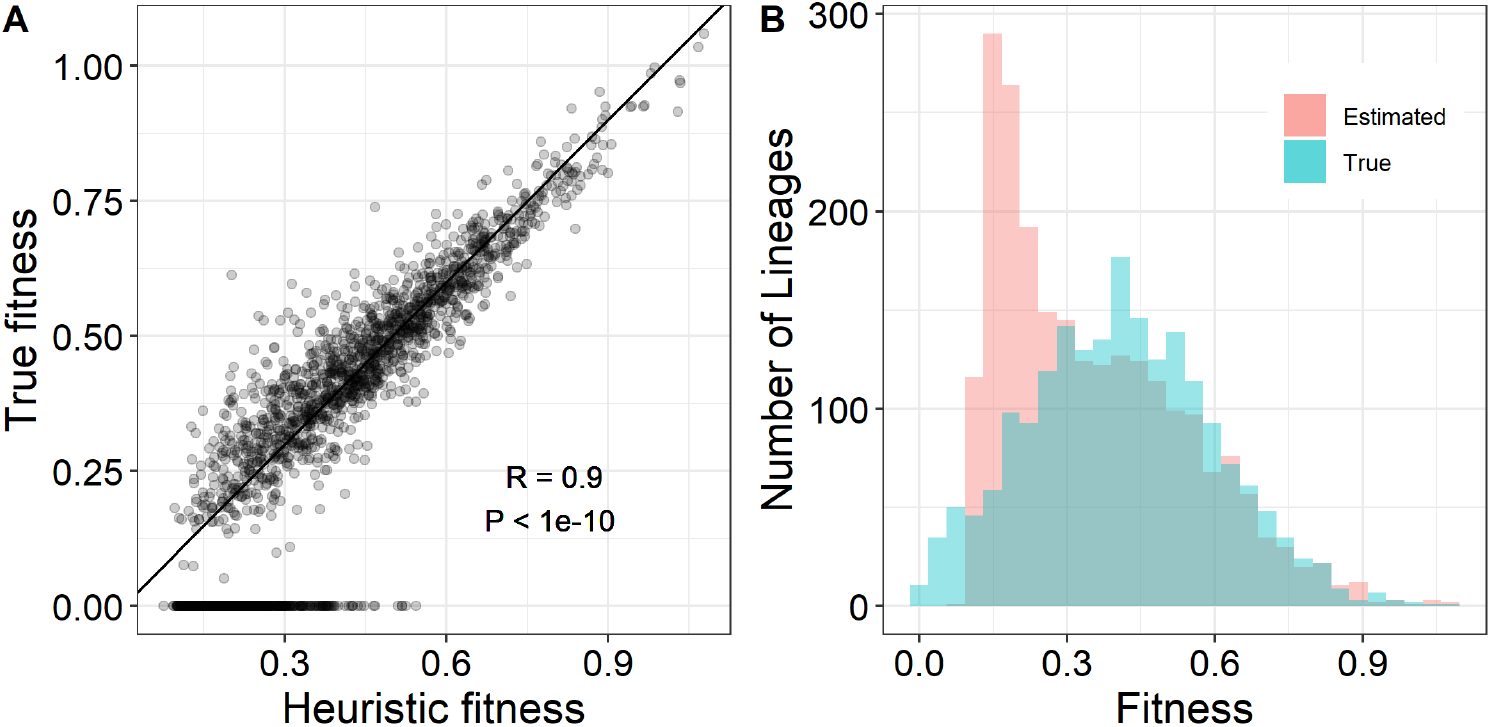
Accuracy of BLT fitness estimates and the shape of the bDFE. **A.** Relationship between the true fitness of a lineage (*x*-axis) and its fitness estimated from a simulated BLT experiment (*y*-axis; see Section 1 for details). The solid line shows the diagonal. The correlation coefficient and the associated P-value are calculated only for the true positives. **B.** True and estimated bDFEs.

**Figure S6.**
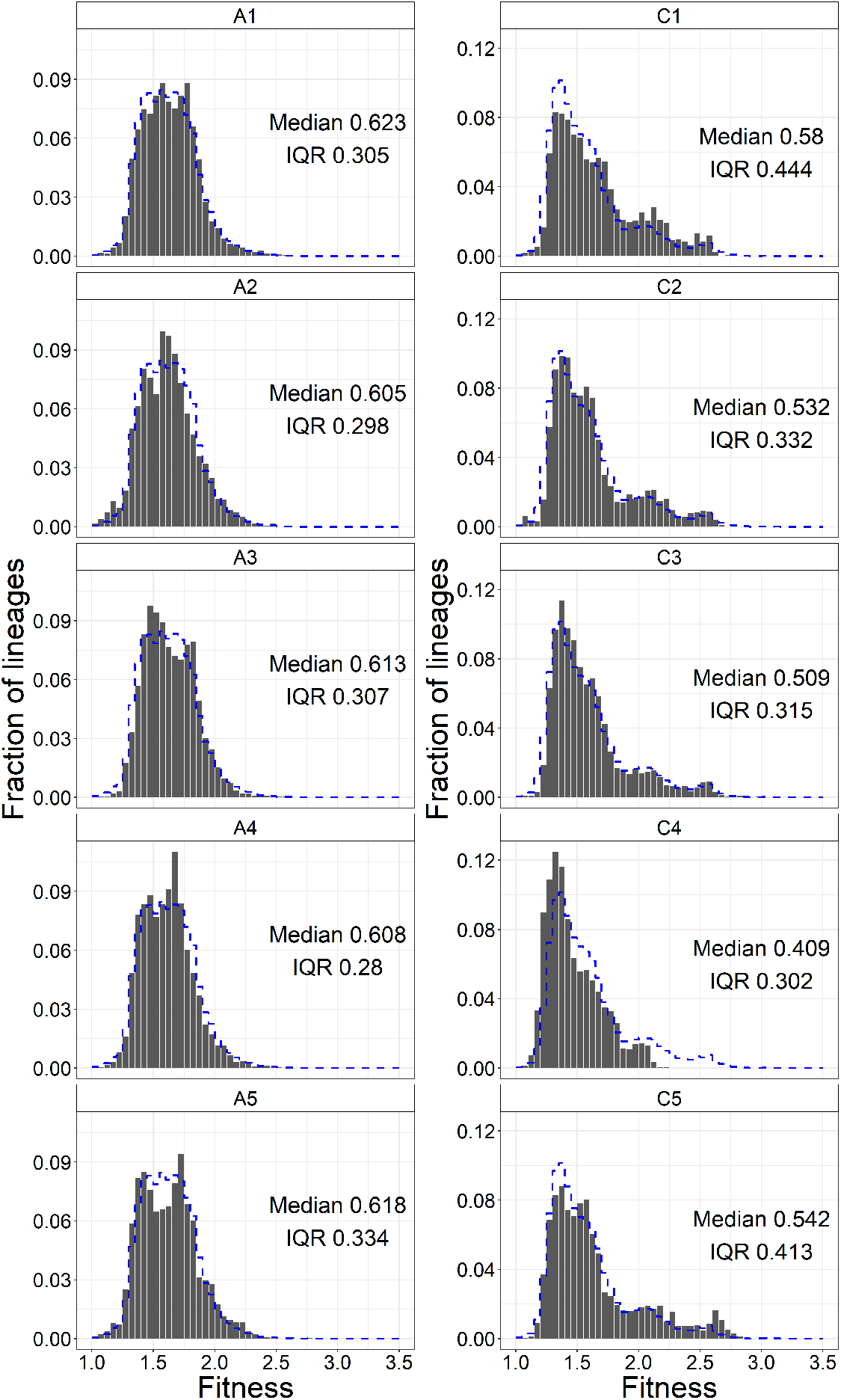
bDFEs estimated for replicate populations individually. The blue dashed line shows the pooled bDFE in the same condition (same as in Figure 2A).

**Figure S7.**
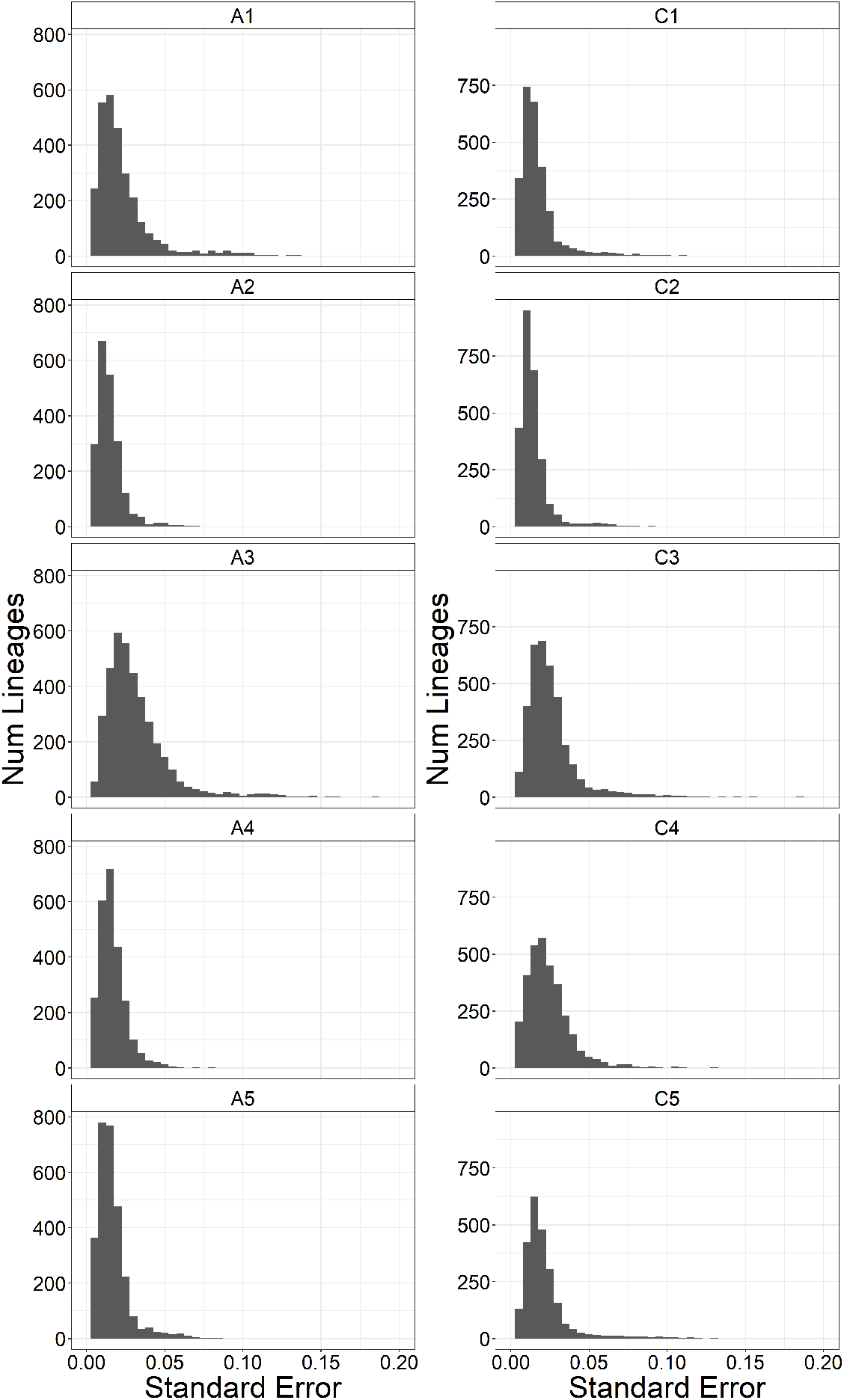
Distributions of standard errors for fitness estimates of adapted lineages for all replicate populations.

**Figure S8.**
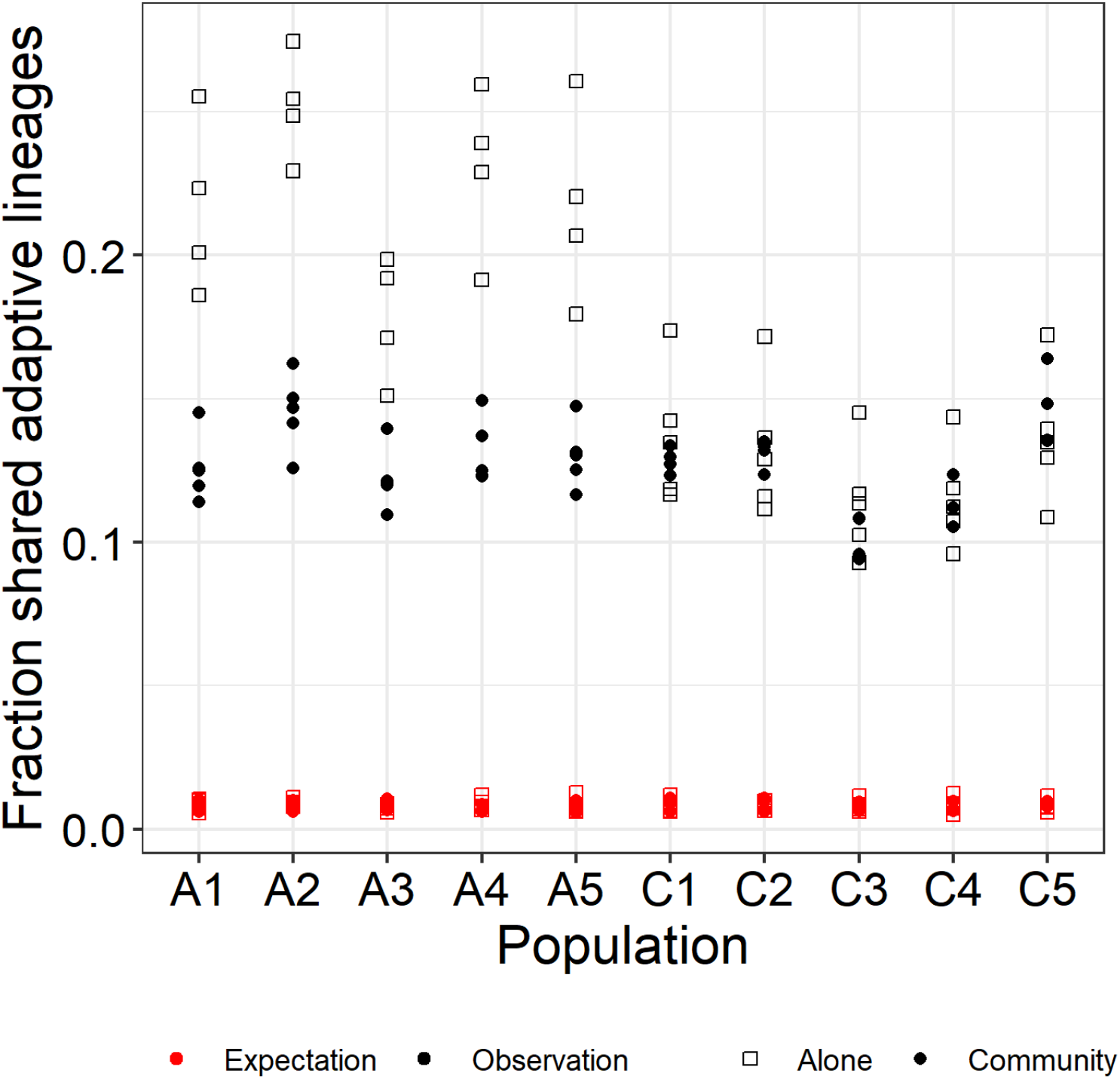
Evidence for pre-existing mutations in the BLT experiments. For each population on the *x*-axis, we plot the fraction of lineages identified as adapted in that population which are also identified as adapted in every other population (*y*-axis, black points) as well as the expectation for this fraction (red points). The expected overlap between populations *i* and *j* is calculated by comparing the observed adaptive lineages in population *i* to a random set of lineages from population *j* that reach detectable frequency. All lineages are considered equally for sampling.

**Figure S9.**
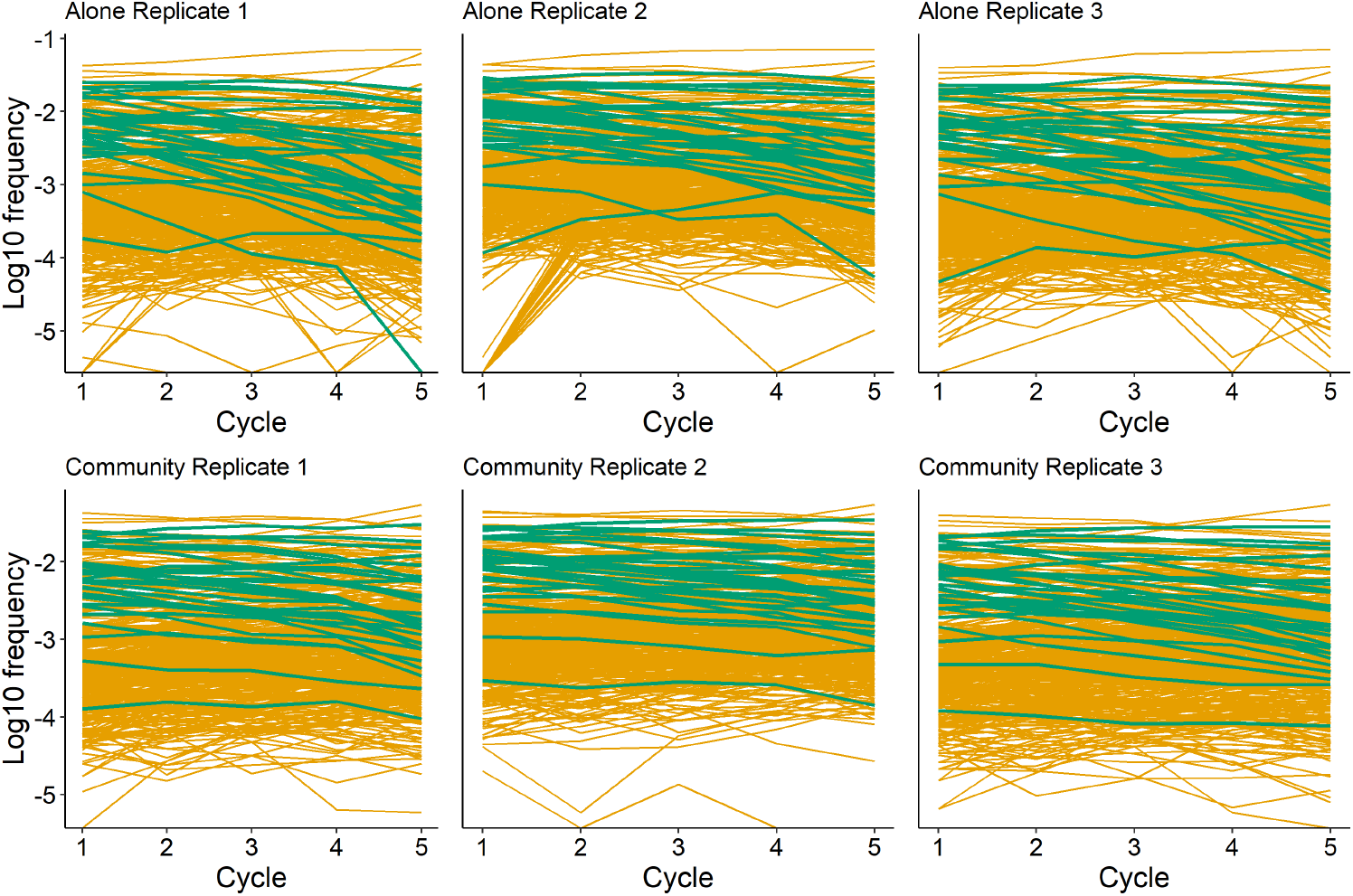
Lineage frequency trajectories in the competition assay. Reference lineages are shown in green, all other lineages are shown in orange.

**Figure S10.**
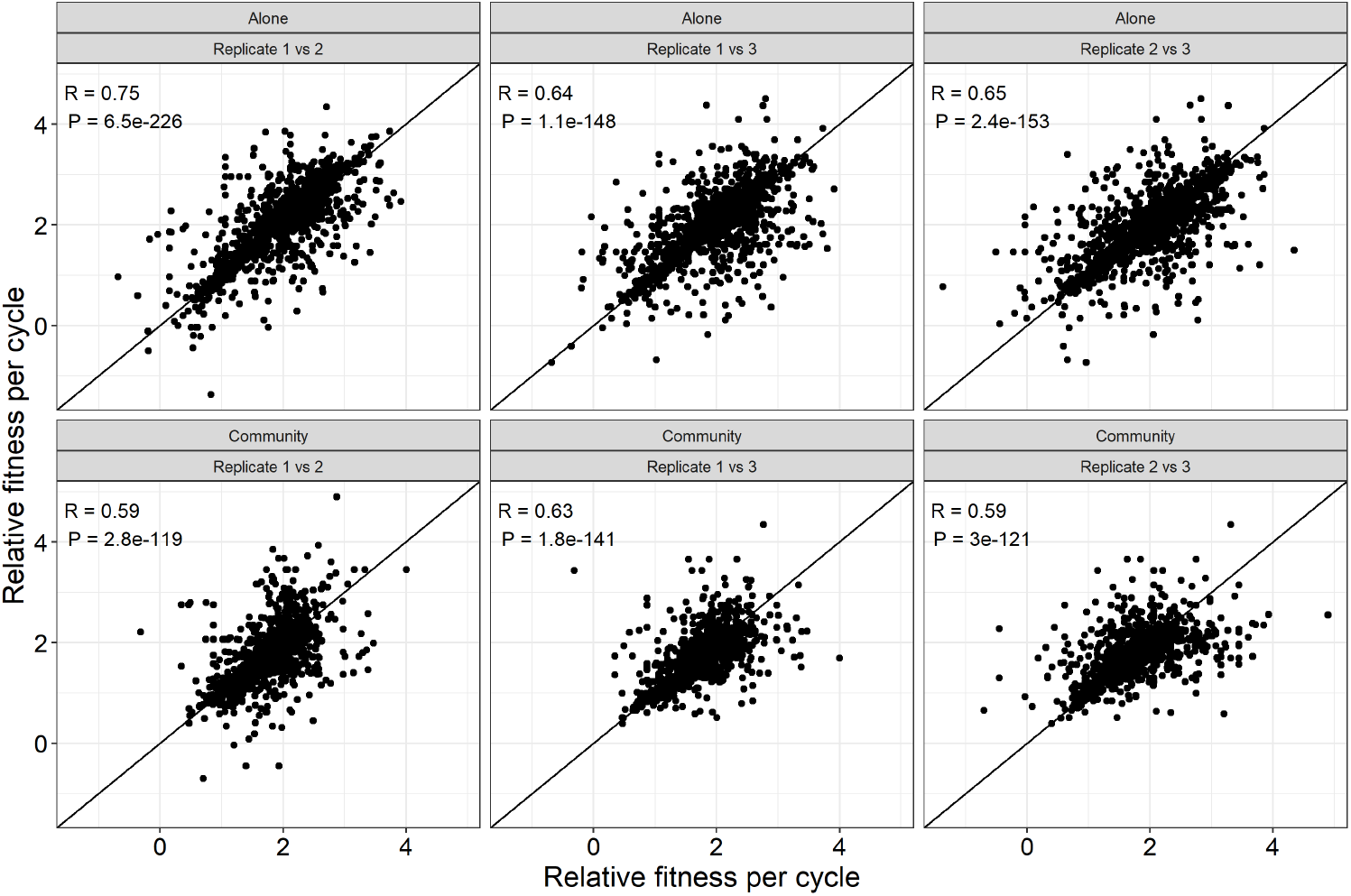
Correlation between fitness estimates across replicates of the competition assay. Line shows the diagonal.

**Figure S11.**
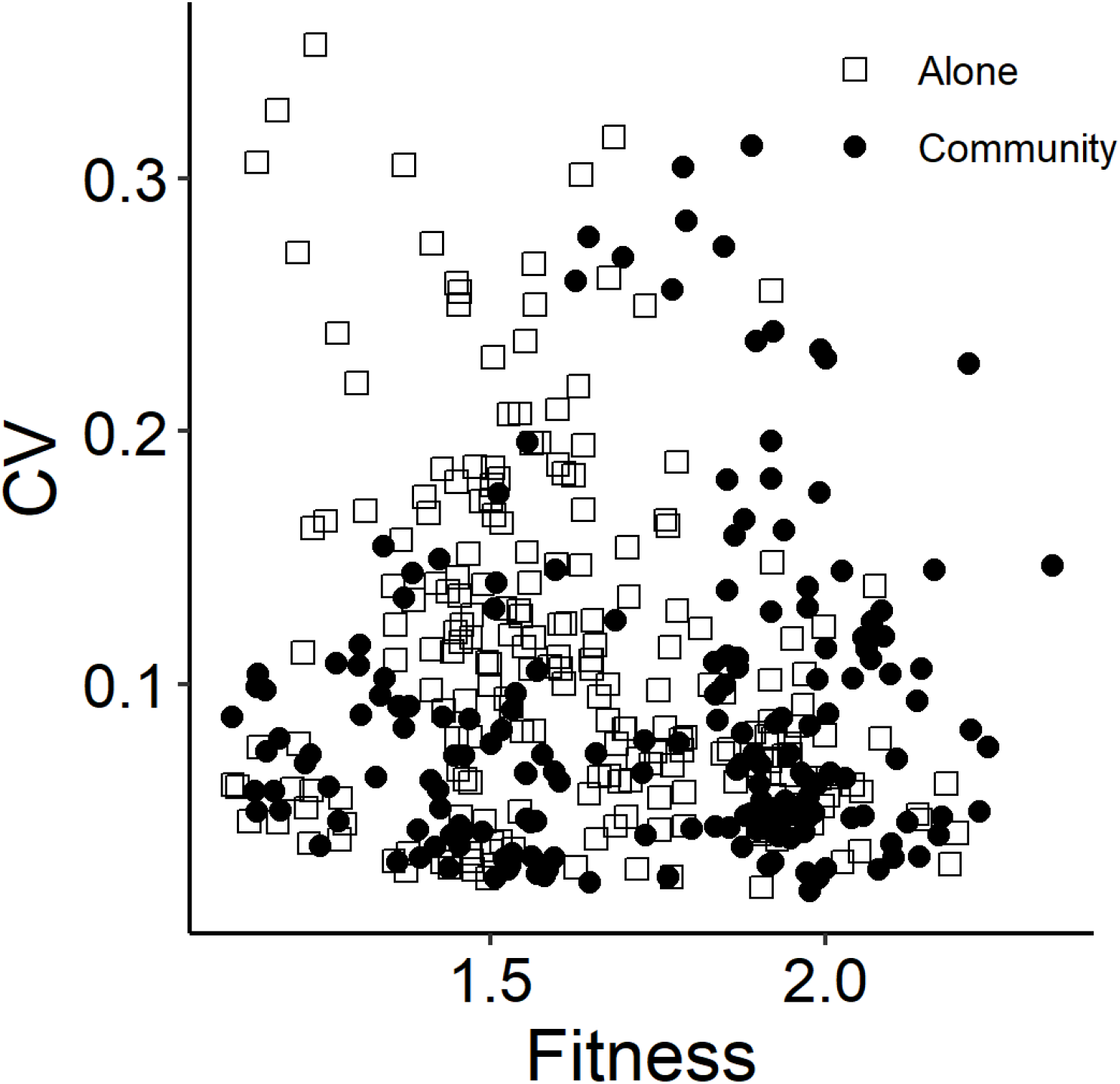
Mean and the coefficient of variation of the fitness estimates in the competition assay.

**Figure S12.**
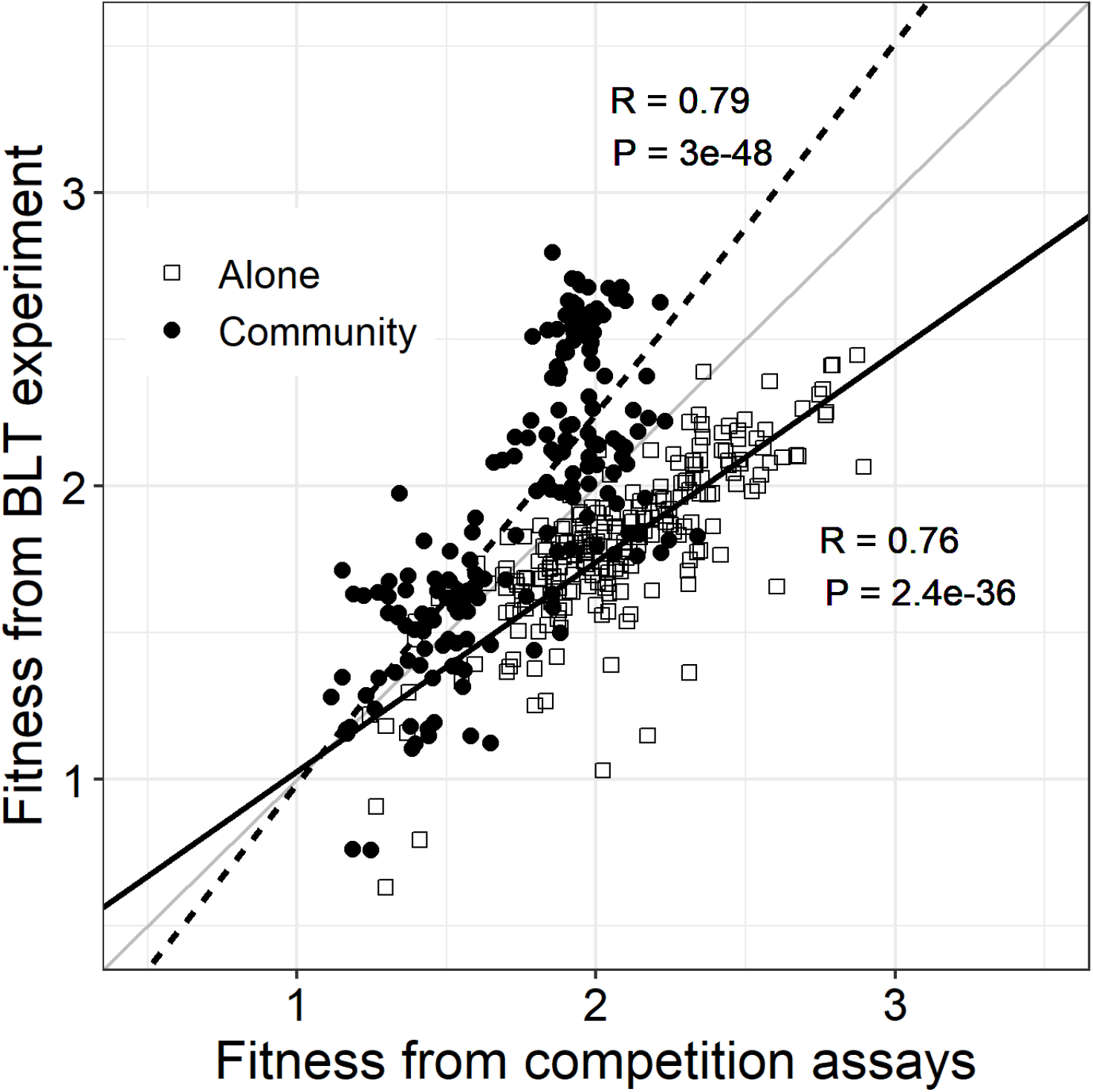
Correlation between BLT and competitive fitness estimates. Solid and dashed lines show linear regressions for the A-mutants (in the A-condition) and C-mutants (in the C-condition), respectively.

**Figure S13.**
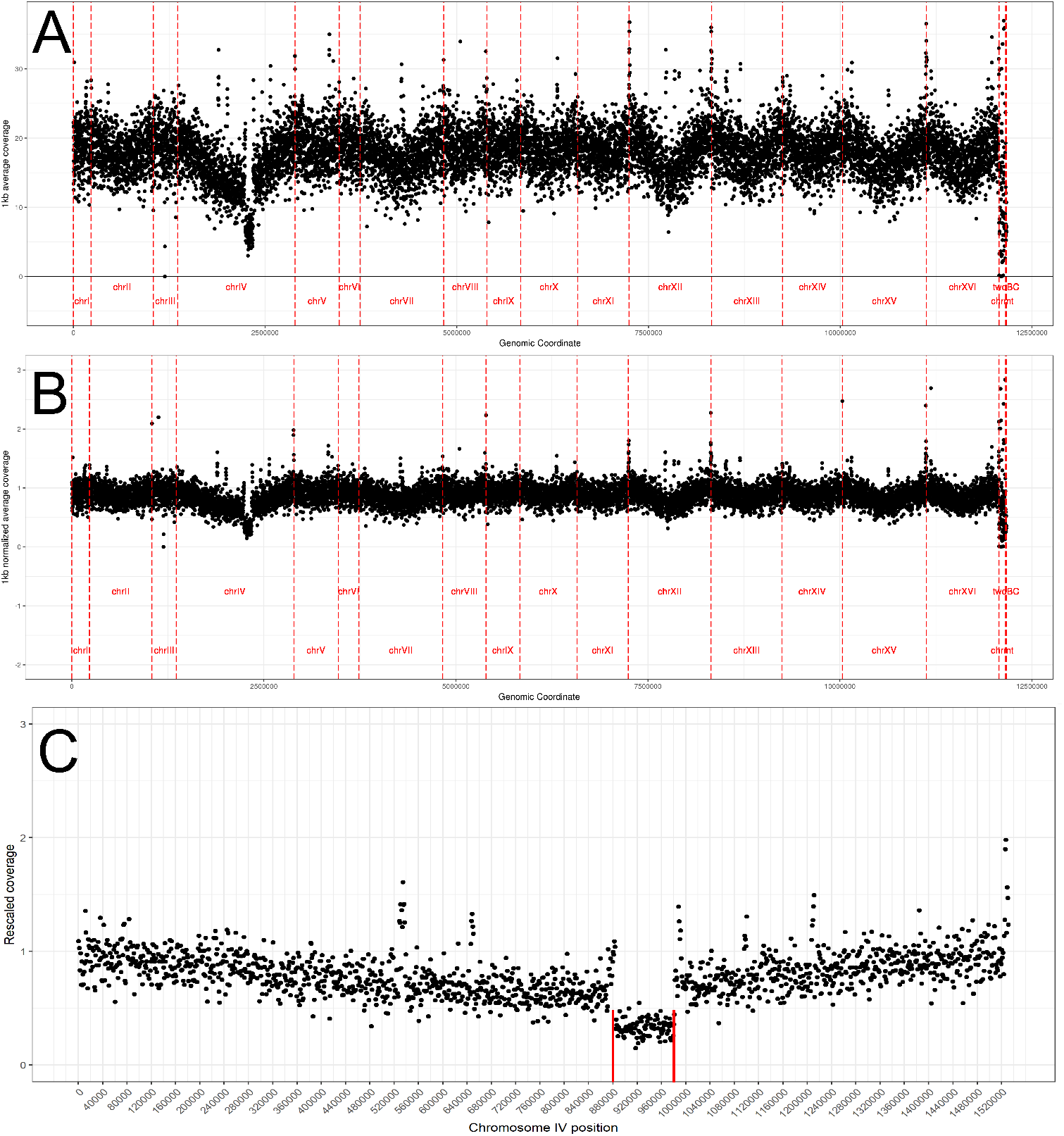
An example illustrating the detection of large CNVs in the genome sequence data. **A.** 1kb-window coverage plot for clone 386817 from population C1. **B.** Same data, after normalization using the relationship between coverage and chromosomal position shown in Figure S14. **C.** Focusing on chrIV to manually identify breakpoints for the chrIV-1n mutation (red lines).

**Figure S14.**
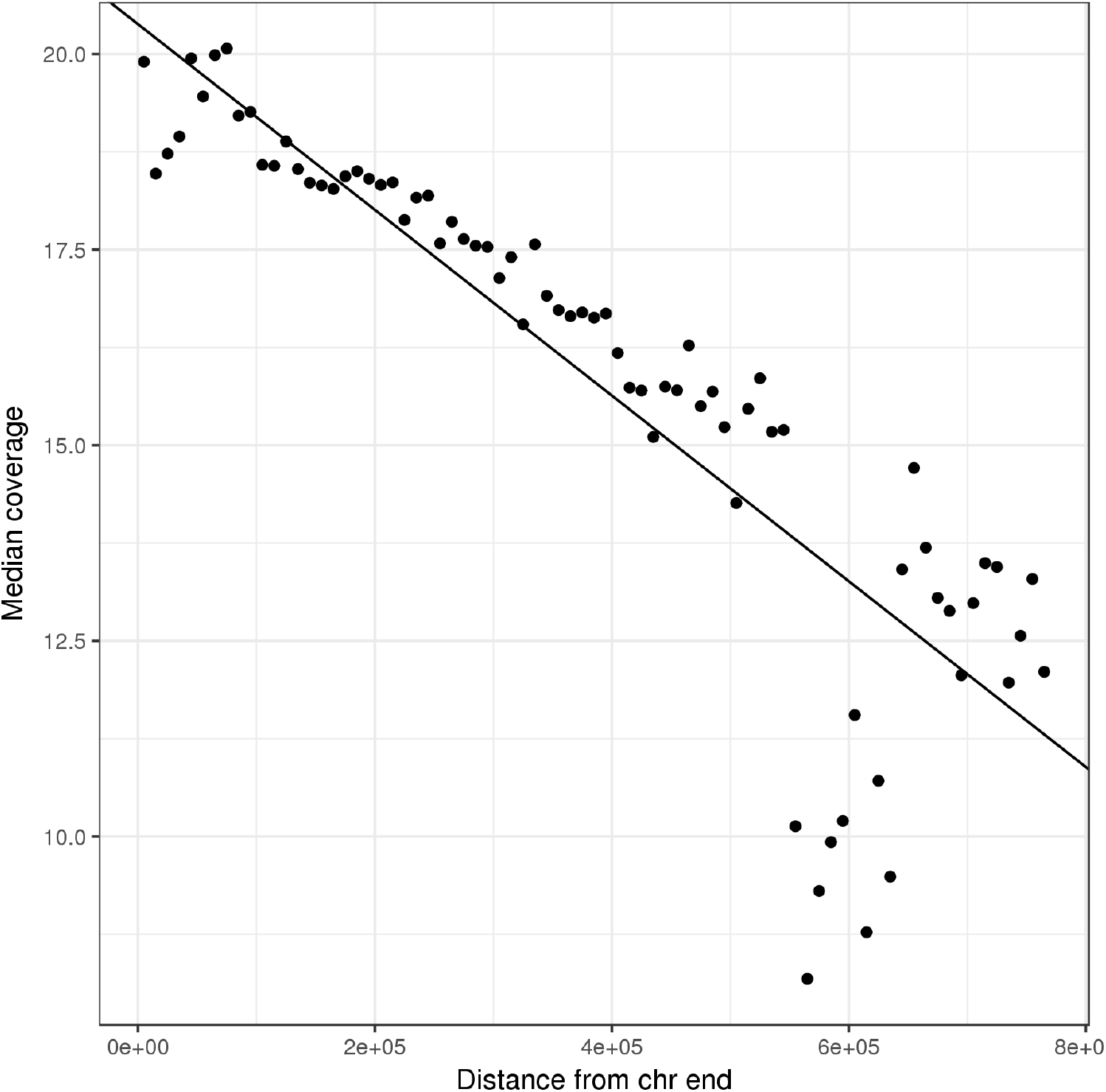
Relationship between coverage and chromosome position. Mean coverage in 1kb windows for clone 386817 from population C1 is shown vs distance of locus from the end of a chromosome.

**Figure S15.**
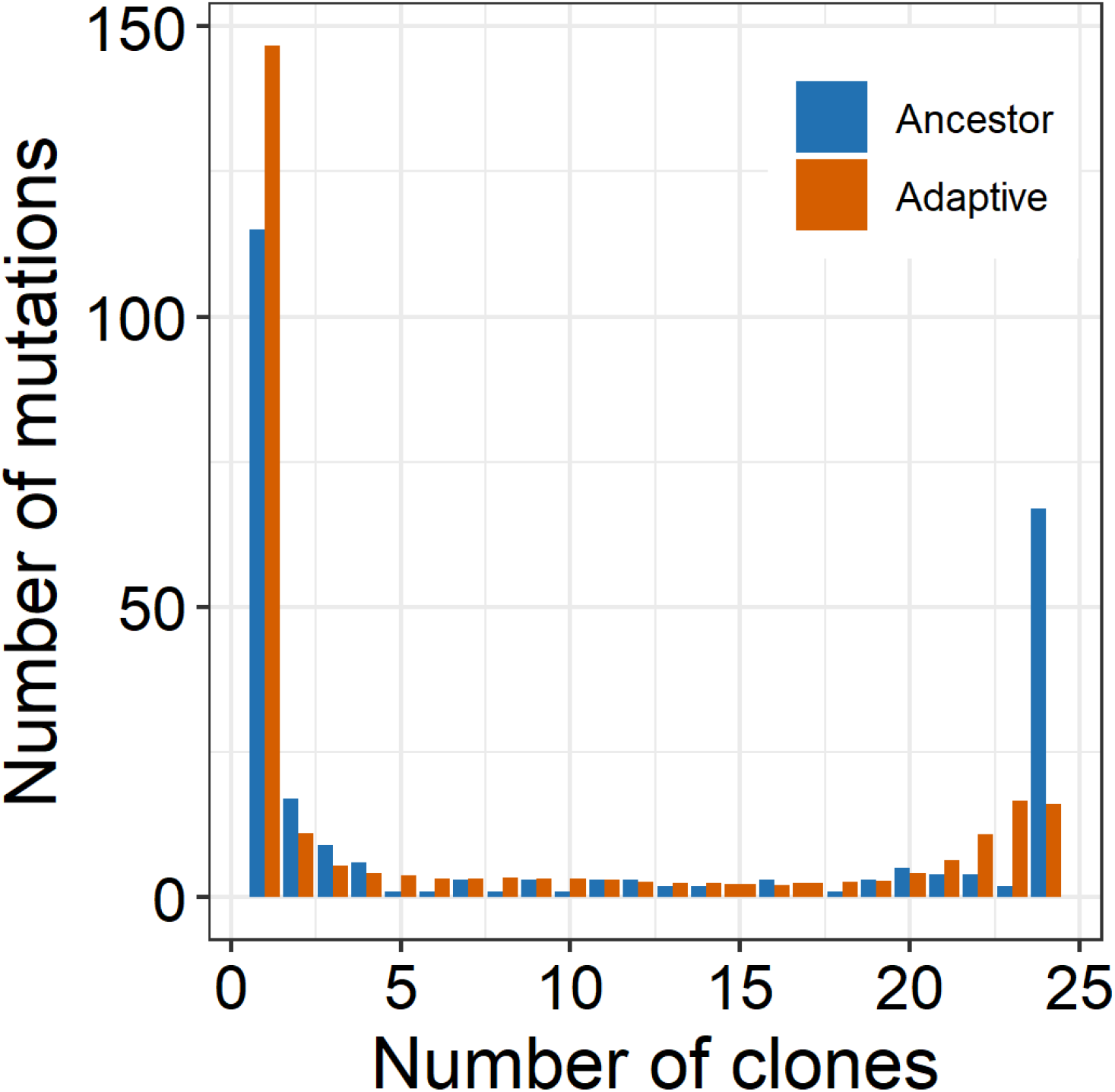
Allele frequency spectrum for ancestral and adapted isolates. Each bar shows the number of small mutations (point mutations and small indels) found in a given number of ancestral or adapted clones. The distribution for adapted clones is averaged over 1000 random draws of 24 clones.

**Figure S16.**
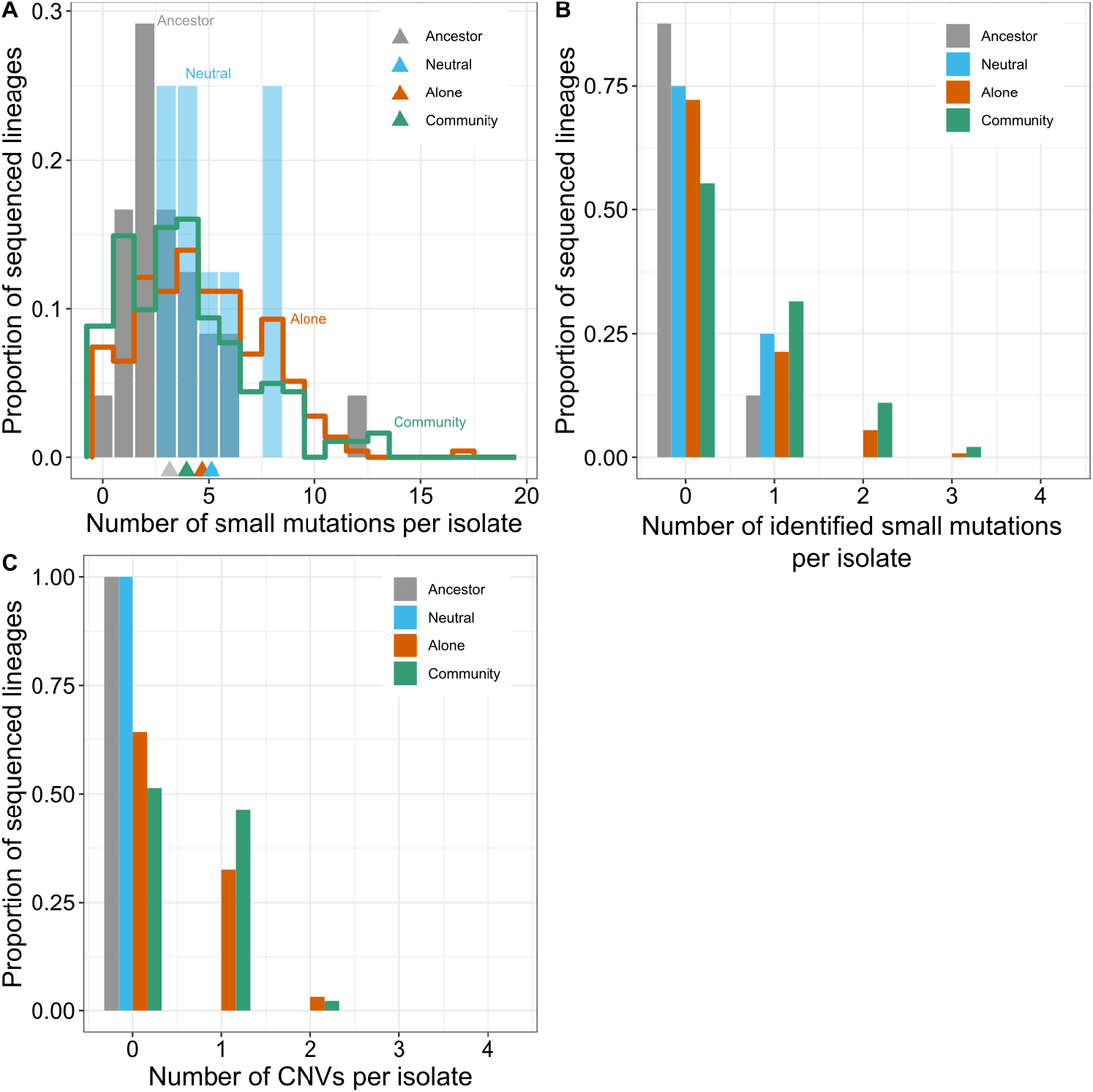
Distribution of mutations per clone. **A.** Number of small mutations detected per sequenced isolate. Triangles on the *x*-axis indicate the means of each distribution (see also Table S2). **B.** Distribution of the number of identified adaptive small mutations per isolate. **C.** Distribution of the number of CNVs per isolate.

**Figure S17.**
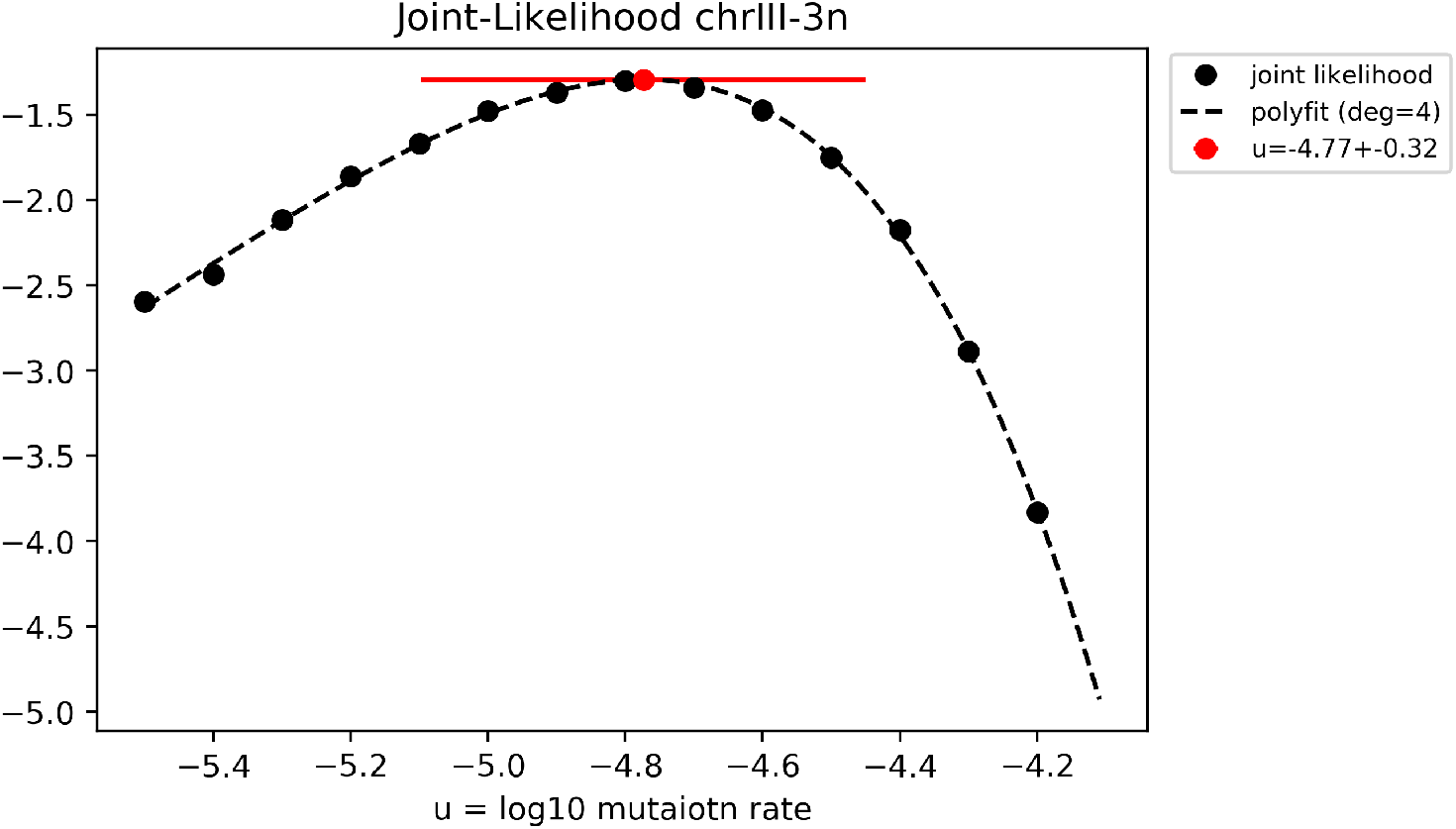
Estimation of the rate of beneficial mutations. The log-likelihood function for the estimation of the chrIII-3n beneficial mutation rate is shown as an example. Red point indicates the identified maximum value. Horizontal red bar shows the standard error calculated based on Fisher Information.

**Figure S18.**
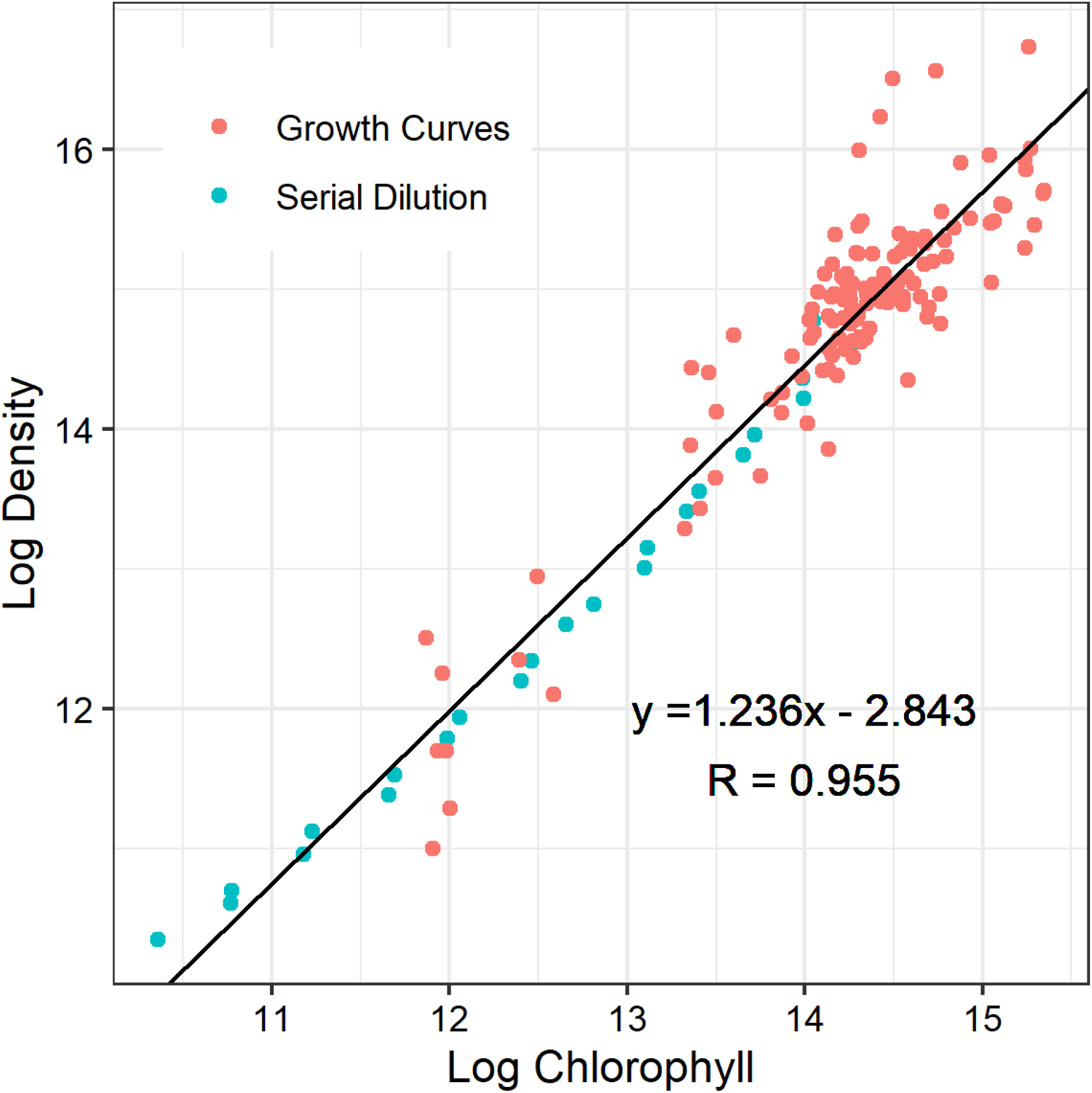
Calibration curve for the alga cell density estimation. *y*-axis shows the natural logarithm of density of alga cells measured by haemocytometer, *x*-axis shows the natural logarithm of chlorophyll fluorescence. The measured alga cultures were obtained either from growth curve experiments (red) or from a serial dilution experiment (blue; see Section 5.1 for details). The black line shows the regression line used for converting chlorophyll fluorescence measurements into cell density estimates.

**Figure S19.**
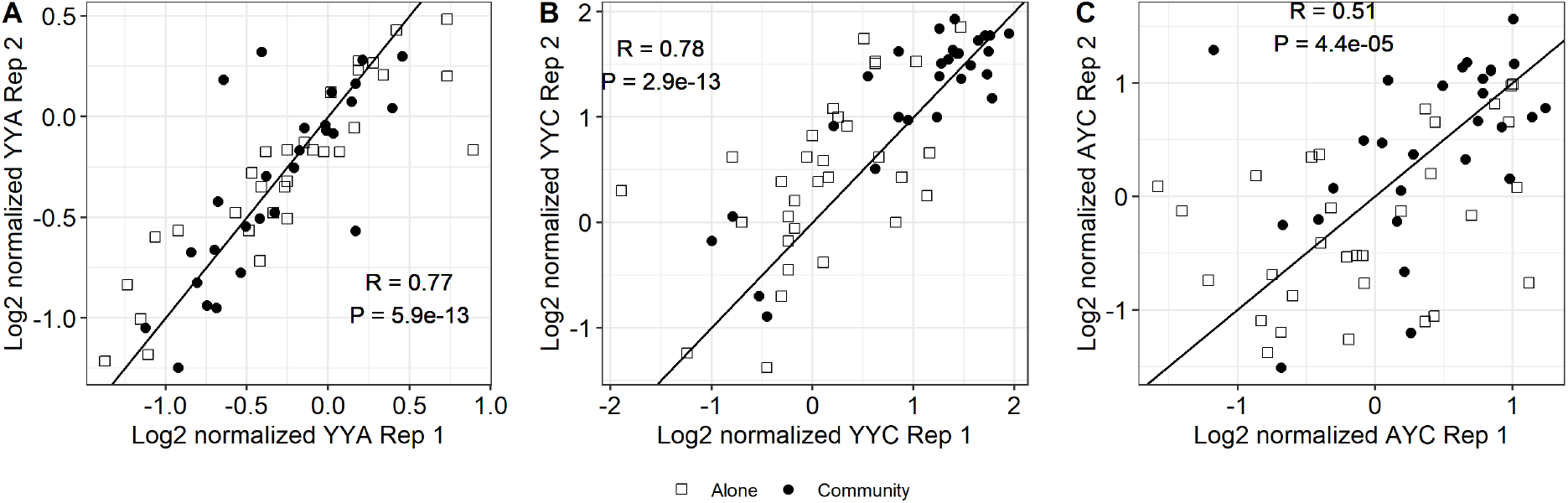
Correlation between yield measurements across replicates. The solid line shows *y* = *x*. Normalization is calculated relative to the ancestral strain.

**Figure S20.**
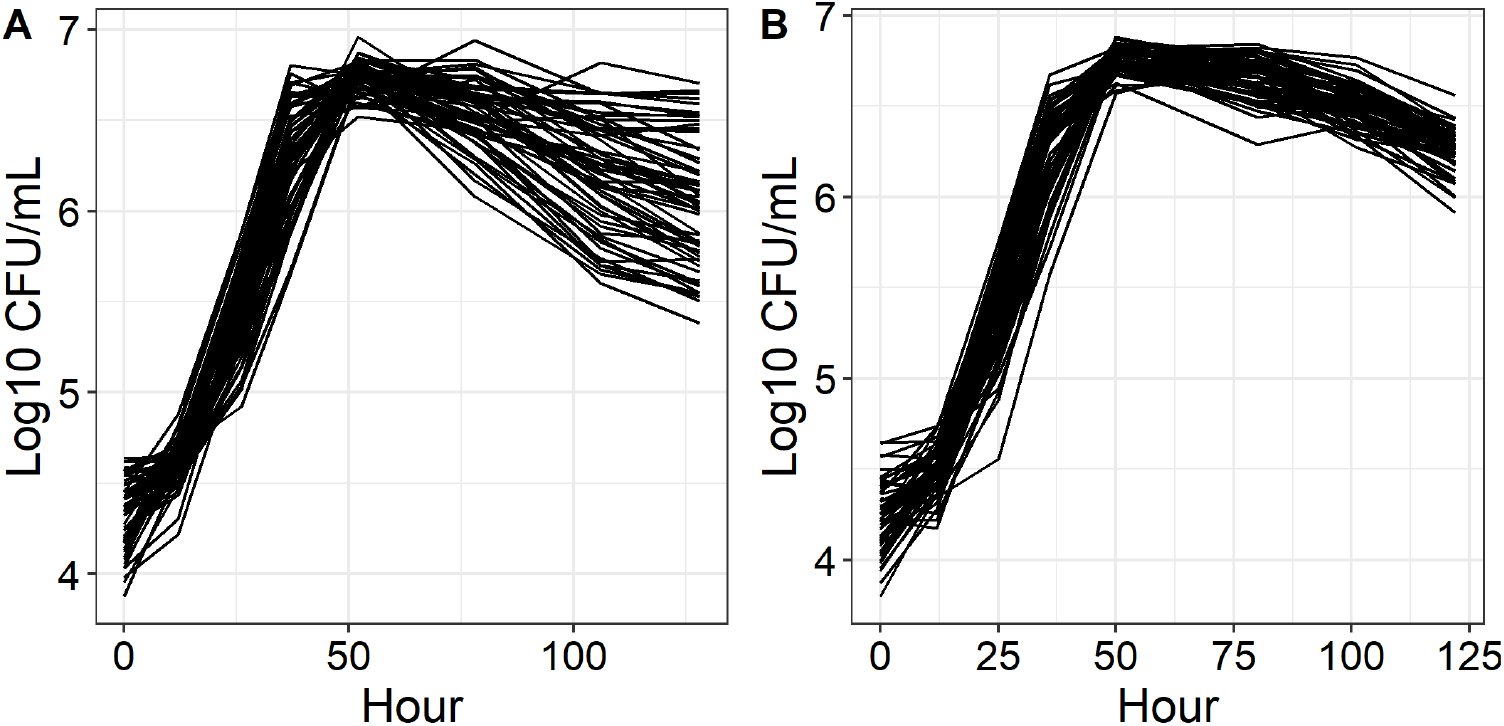
Mutant growth curves in the A-condition. Growth curves for the 59 adapted yeast mutants are shown, from which *r* and *K* values are estimated as described in Methods in the main text. **A.** Replicate 1. **B.** Replicate 2.

**Figure S21.**
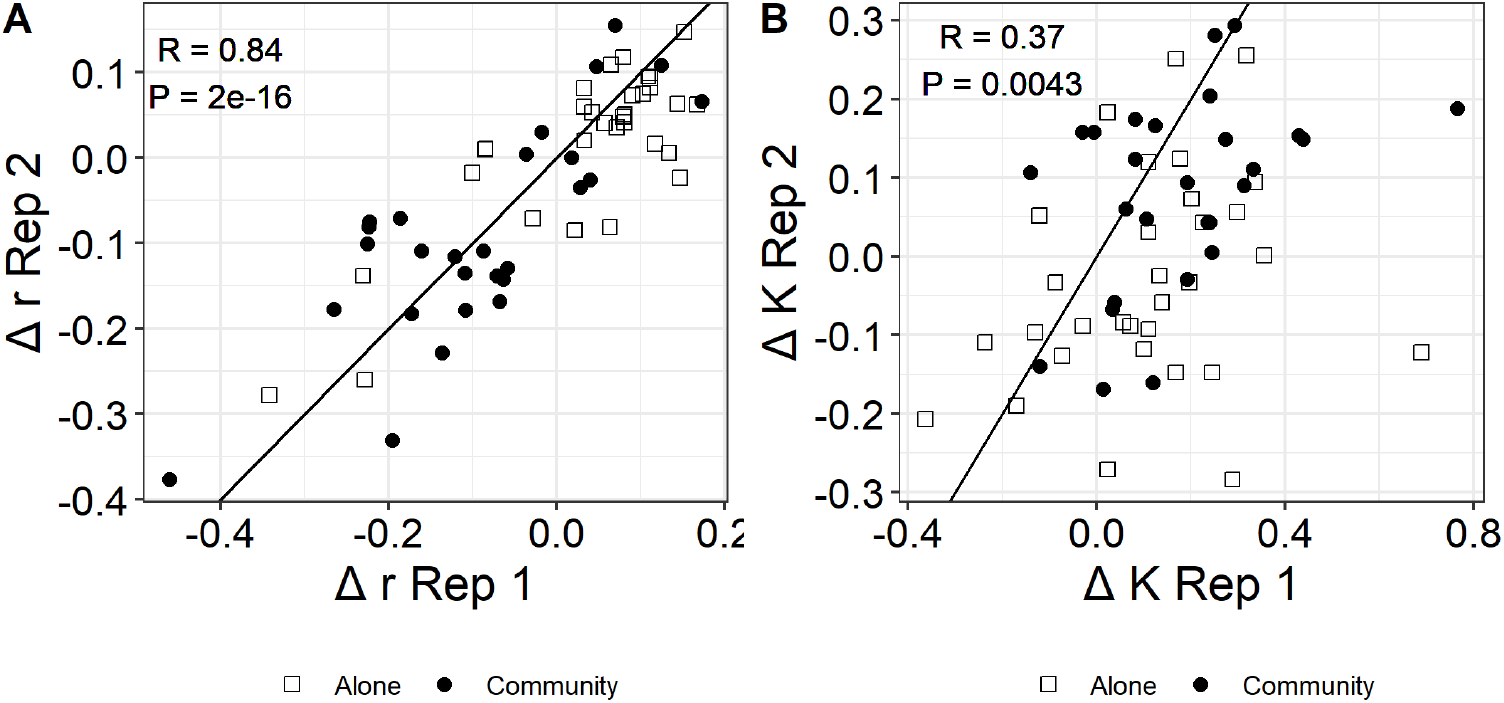
Correlation between replicates for *r* and *K* measurements.

**Figure S22.**
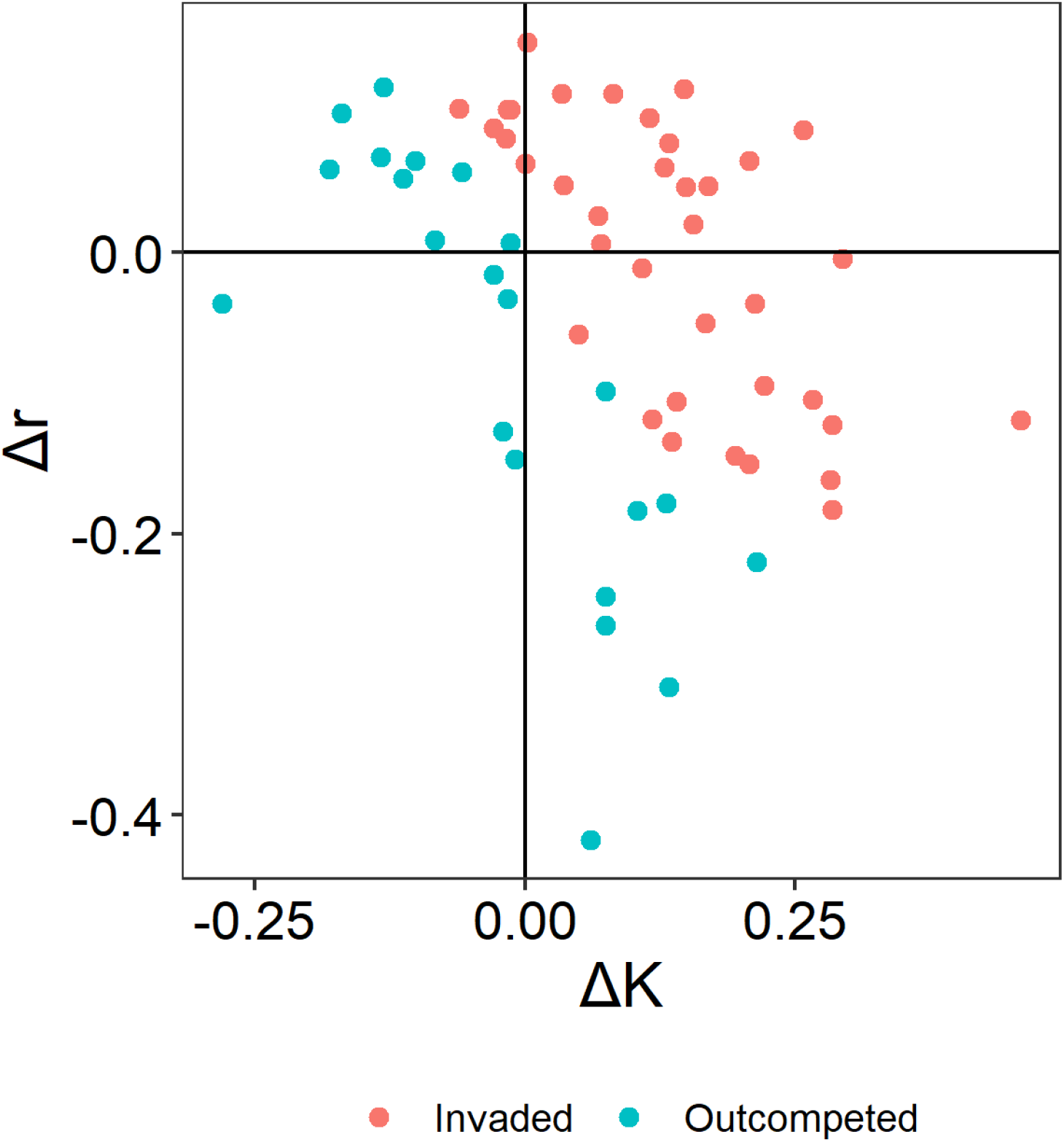
Mutant invasion analysis. Yeast mutants that are able to invade the wildtype population in the logistic growth simulations are shown in red, those that are not able to invade are shown in blue (see Section 5.2 for details).

## 9 Supplementary Tables

**Table S1.** *P*-values for comparison of absolute abundances and net population change rates across conditions. P-values are obtained from t-tests and corrected for multiple testing using the Benjamini-Hochberg procedure.

| Measurement | Day | Yeast | Alga |
| --- | --- | --- | --- |
| Absolute<br>abundance<br>(Figure 1) | 0 | $10^{-3}$ | $3 \times 10^{-3}$ |
| | 1 | $6 \times 10^{-4}$ | $8 \times 10^{-4}$ |
| | 2 | $3 \times 10^{-3}$ | $5 \times 10^{-5}$ |
| | 3 | $10^{-5}$ | $9 \times 10^{-4}$ |
| | 4 | 0.05 | $10^{-5}$ |
| | 5 | 0.73 | $10^{-5}$ |
| Growth rate<br>(Extended Data<br>Figure 1) | 0-1 | 0.07 | 0.03 |
| | 1-2 | $7 \times 10^{-5}$ | 0.85 |
| | 2-3 | 0.85 | $10^{-4}$ |
| | 3-4 | $5 \times 10^{-4}$ | $10^{-3}$ |
|  | 4-5 | 0.97 | 0.07 |

**Table S2.** Statistics of mutational counts among sequenced isolates.

| Isolates |  | Small variants |  |  |  | CNVs |  |
| --- | --- | --- | --- | --- | --- | --- | --- |
| Type | Number | Total | Per isolate | Adaptive |  | Total | Per isolate |
|  |  |  |  | Expected | Identified |  |  |
| Ancestral | 24 | 76 | $3.17 \pm 0.51$ | – | 3 | 0 | – |
| Neutral | 8 | 41 | $5.13 \pm 0.72$ | – | 2 | 0 | – |
| A-mutants | 215 | 1008 | $4.69 \pm 0.20$ | (0, 288) | 76 | 84 | $0.39 \pm 0.04$ |
| C-mutants | 181 | 717 | $3.96 \pm 0.22$ | (0, 110) | 109 | 92 | $0.51 \pm 0.04$ |
| Total | 428 | 1842 | – | – | 190 | 176 | – |

**Table S3.** Numbers of isolates carrying the most common adaptive mutations and the estimated mutation rates. Only clones with unique barcodes are shown. Column “Seq. isolates” shows the total number of sequenced isolates with unique barcodes sampled from each culture. Last row shows the estimated *u_m_ =* log_10_ *U_m_*.

| Culture | Seq.<br>isolates | Mutation class $m$/Locus | | | | | | | | | |
| --- | --- | --- | --- | --- | --- | --- | --- | --- | --- | --- | --- |
|  |  | chrIV-1n | chrIII-3n | chrXIV-3n | chrIV-3n | chrIX-3n | chrXIII-3n | YIL169C | HEM1 | HEM2 |  |
| A1 | 13 | 0 | 2 | 0 | 0 | 1 | 0 | 0 | 0 | 0 |  |
| A2 | 17 | 3 | 3 | 0 | 2 | 0 | 0 | 0 | 0 | 0 |  |
| A3 | 12 | 3 | 3 | 0 | 1 | 0 | 0 | 0 | 0 | 0 |  |
| A4 | 19 | 3 | 5 | 0 | 1 | 0 | 0 | 0 | 0 | 0 |  |
| A5 | 18 | 5 | 1 | 0 | 0 | 1 | 0 | 1 | 0 | 0 |  |
| C1 | 25 | 8 | 0 | 2 | 1 | 3 | 2 | 1 | 1 | 0 |  |
| C2 | 16 | 1 | 1 | 6 | 0 | 2 | 0 | 0 | 0 | 1 |  |
| C3 | 21 | 5 | 0 | 0 | 0 | 0 | 0 | 0 | 2 | 1 |  |
| C4 | 11 | 5 | 0 | 0 | 0 | 0 | 0 | 0 | 1 | 1 |  |
| C5 | 24 | 6 | 1 | 2 | 1 | 1 | 0 | 1 | 0 | 0 |  |
| | $s_m^A$ | 0.916 | 0.855 | 0.593 | 1.18 | 0.527 | 0.315 | 0.708 | 0.955 | 0.874 | |
| | $s_m^C$ | 1.17 | 0.689 | 1.05 | 1.3 | 0.492 | 0.575 | 0.744 | 1.34 | 1.24 | |
| | $\hat{u}_m$ | -5.32 | -4.77 | -5.51 | -6.49 | -4.14 | -4.58 | -5.11 | -6.65 | -6.59 | |

**Table S4.** Multiple regression analysis of fitness against life-history traits. *β_r_* and *β_K_* are standardized regression coefficients for *r* and *K*, respectively. *P*_r_ and *P_K_* are *P*-values indicating whether adding *r* or *K* to the model significantly improves the fit above and beyond the single-variable model.

| Condition | $R^2$ | $P$ | $\beta_r$ | $P_r$ | $\beta_K$ | $P_K$ |
| --- | --- | --- | --- | --- | --- | --- |
| A | 0.067 | 0.14 | 0.15 | 0.72 | 0.73 | 0.05 |
| C | 0.374 | $2 \times 10^{-6}$ | -0.81 | 0.002 | 0.72 | 0.002 |

## Footnotes

1 This number is larger than 0.18 expected based on the yeast mutation rate, but the difference is not statistically significant (*P* = 0.08, t-test, expected *μ* = 0.18).

2 This is confirmed by the fact that we find two mutations at driver loci in the neutral clones (see Figure S16 and Table S2).

